# Mechanics of MTOC clustering and spindle positioning in budding yeast *Cryptococcus neoformans*

**DOI:** 10.1101/690651

**Authors:** Saptarshi Chatterjee, Subhendu Som, Neha Varshney, PVS Satyadev, Kaustuv Sanyal, Raja Paul

## Abstract

The dynamic process of mitotic spindle assembly depends on multitudes of inter-dependent interactions involving kinetochores (KTs), microtubules (MTs), spindle pole bodies (SPBs), and molecular motors. Before forming the mitotic spindle, multiple visible microtubule organizing centers (MTOCs) coalesce into a single focus to serve as a SPB in the pathogenic budding yeast, *Cryptococcus neoformans*. To explain this unusual phenomenon in the fungal kingdom, we propose a ‘search and capture’ model, in which cytoplasmic MTs (cMTs) nucleated by MTOCs grow and capture each other to promote MTOC clustering. Our quantitative modeling identifies multiple redundant mechanisms mediated by a combination of cMT-cell cortex interactions and inter-cMT coupling to facilitate MTOC clustering within the physiological time limit as determined by time-lapse live-cell microscopy. Besides, we screen various possible mechanisms by computational modeling and propose optimal conditions that favor proper spindle positioning - a critical determinant for timely chromosome segregation. These analyses also reveal that a combined effect of MT buckling, dynein pull, and cortical push maintains spatiotemporal spindle localization.

## INTRODUCTION

Spatio-temporal dynamics of organelle-constituents relies primarily on the active organization of microtubules (MTs). MTs are semiflexible polymeric filaments having rapidly polymerizing plus ends and slowly polymerizing minus ends. The growth and shrinkage of MTs are governed by intrinsic dynamic instability parameters [1]. In animal cells, the centrosome is a major MT nucleating center which forms a radial MT array [2–4] along with other membrane organelles such as the Golgi apparatus facilitating many noncentrosomal, radial MT networks [2, 5–9]. The cellular structures that harbor the ability to nucleate MTs and organize a radial network are often referred to as Microtubule Organizing Centers (MTOCs) [2, 10].

In the ascomycete budding yeast *Saccharomyces cerevisiae*, spindle pole bodies (SPBs), embedded on the nuclear envelope (NE) are analogous to centrosomes. The SPBs are enriched with distinct *γ*-tubulin receptors that organize the cytoplasmic MTs (cMTs) and nuclear MTs (nMTs) [11, 12]. The MT nucleation system in *S. cerevisiae* involves *γ*-tubulin small complexes (*γ*-TuSCs) and their receptors Spc110 and Spc72 [2, 12]. This MT nucleation system provides SPBs distinct nuclear and cytoplasmic domains indicating that a single MTOC can accommodate nucleation sites for both cMTs and nMTs [2, 10, 13].

Two major fungal phyla, Ascomycota and Basidiomycota [14–16] shared a common ancestor more than 500 million years ago [17]. While extensive studies have been carried out in the context of cell division in hemiascomycetous budding yeasts such as *S. cerevisiae*, the process of chromosomal partitioning in organisms belonging to Basidiomycota remains under-studied [14–16, 18]. A relatively better model basidiomycetous budding yeast is *Cryptococcus neoformans*, a human pathogen. This pathogen is responsible for life-threatening diseases, including fungal meningitis, and emerges as the 5th leading cause of mortality among the immuno-compromised individuals [19].

Like *S. cerevisiae*, cells divide by budding in *C. neoformans* as well (Fig. 1A, Movie M1). The major sequence of mitotic events from the formation of bud until chromosome segregation in *C. neoformans* can be categorized in the following temporal order: (a) Bud inception at the end of G1/onset of S phase [16], (b) clustering of KTs as a single punctum at the periphery of the nucleus after bud initiation [16] (Fig. 1A, Movie M1), (c) nuclear migration into the daughter bud followed by spindle formation; (d) localization of the spindle near the bud-neck junction/septin ring (Fig. 1A) and eventually (e) equal nuclear division mother and daughter bud (Fig. 1A, Movie M1).

**FIG. 1.**
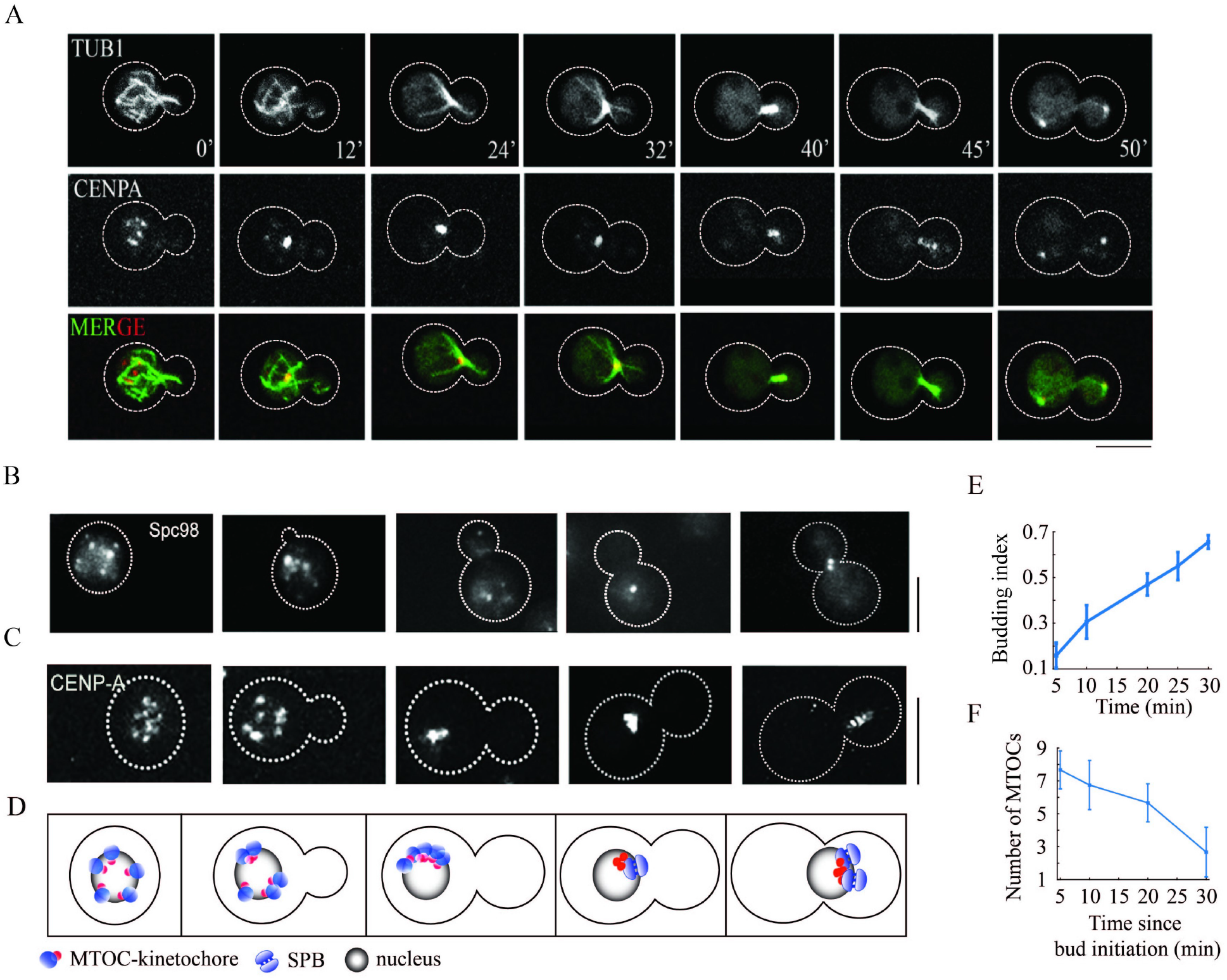
Cell cycle stages in *C. neoformans* wild-type cell as observed from live-cell imaging. (A) Time series snapshots showing the localization of MTs (GFP-Tub1, depicted in green in the MERGE panel) and KTs (mCherry-CENP-A, depicted in red in the MERGE panel) at different stages of the cell cycle in a wild-type cell. The cell was subjected to time-lapse video microscopy for 60 min. The timestamp (in min) for each image is annotated in the uppermost TUB1 panel. Bar, 5 *µ*m. (B-C) Clustering of MTOCs and KTs during mitosis in *C. neoformans*. Images of cells showing the localization of (B) Spindle pole body protein, Spc98, and (C) KT marker CENP-A at different stages of the cell cycle in the wild-type cells. Bar, 5 *µ*m. The snapshots of the co-localization of Spc98 and CENP-A (using a different strain) were shown in [20]. (D) Schematic depicting the concomitant happening of the processes shown in (B) and (C). (E) Progression of bud growth with time. (F) Number of MTOCs in the cell as a function of time estimated from the bud size.

While the overall process of nuclear division is conserved across Ascomycota and Basidiomycota, from a coarse-grained viewpoint, several intriguing differences exist in the spatiotemporal dynamics of MTOCs, KTs, nuclear migration, the specific location for spindle formation and nuclear division [14, 16, 21, 22]. We note these distinct differences in the following: (a) A single MTOC is found embedded on the NE at the onset of the cell cycle in ascomycetous budding yeast *S. cerevisiae* [23]. Here the MTOC serves as the SPB. At the G1/S phase, SPB duplicates and the KT-MT connections begin to form. However, the NE never breaks down, leading to closed mitosis [24, 25]. In contrast to *S. cerevisiae*, where the SPB being the only MTOC, several MTOCs are present in the basidiomycete budding yeast *C. neoformans* during interphase [14, 20, 26] (Fig. 1B). Moreover, it undergoes semi-open mitosis marked by the transient breakdown/rupture of NE during metaphase to anaphase transition [15, 27]. (b) In *S. cerevisiae*, the KTs are grouped into a single cluster and KT attachments to the SPBs by MTs are maintained through most of the cell cycle [24, 28–30]. Contrary to that, in *C. neoformans*, KTs are unclustered during interphase [15, 16, 20]. As the cell advances through the cell cycle, the KTs gradually cluster into a single punctum, plausibly via various MT-mediated interactions. Interestingly, a previous study [20] as well as our experiments (Fig. 1B-1D) identify that the KTs colocalize with the MTOCs. The clustering of KTs into a single punctum and the clustering of MTOCs into a single SPB happen concomitantly (Fig. 1B-1D). (c) In *S. cerevisiae*, prior to the chromosome segregation, the nucleus migrates to the proximity of mother-daughter bud neck junction [14, 16, 31, 32]. In *C. neoformans*, the sequence of events characterizing nuclear migration before the division is somewhat different from those of *S. cerevisiae*. First, the nucleus entirely moves to the daughter bud (Fig. S1). Second, the SPB duplication occurs either in the daughter bud or when the nucleus is migrated close to the bud-neck junction (namely the septin ring) as depicted in Fig. 1B, Fig. 1D. Subsequently, SPB biorientation, spindle formation, and localization of the spindle structure *inside the daughter bud near the septin ring* pave the way for proper division [14–16]. Interestingly, the features of having multiple MTOCs and nuclear division in the daughter bud are also characteristics of another basidiomycete yeast, *Ustilago maydis* [33]. Unlike budding yeast, the fission yeast *Schizosaccharomyces pombe* has a cylindrical rod-shaped geometry. Interestingly, *S. pombe* also possesses multiple interphase MTOCs (iMTOCs) and SPBs, which nucleate MTs. During interphase, fission yeast forms 3-5 MT bundles along the long axis of the cell organized by astral microtubules nucleated from duplicated SPBs and interphase MTOCs (iMTOCs). This antiparallel MT organization plays a crucial role in force generation, and nucleus positioning within the cell [34, 35]. On another instance, in *C. neoformans* the MTOCs fuse together to form the mature SPB while remaining attached to the NE. Nevertheless, the presence of multiple MTOCs in a basidiomycete budding yeasts *C. neoformans* and *U. maydis* could be reminiscent of fission yeast. Thus, *C. neoformans* may combine features of both budding and fission yeasts. While framing a mechanistic model of mitotic division in budding yeast *C. neoformans*, we took a cue from our previous studies [14, 16] and extended the ideas of various force interactions from another budding yeast *S. cerevisiae* for its geometric similarity and similarities in cell cycle stages that lead to chromosome segregation.

Self-assembly of MTOCs is orchestrated by MT mediated ‘search and capture’ mechanism [14, 16]. Essential characteristics of the clustering mechanisms are shared across a diverse set of organisms as well as several organelle assemblies (e.g., Golgi assembly and stacking, mitochondrial assembly, multi-centrosomal clustering [36] etc.) and are not necessarily organism-specific [8]. The mechanics of MTOC clustering before the formation of the SPB in *C. neoformans* remains elusive and deciphering the same experimentally is challenging. The complete clustering in wild-type cells occurs within *∼* 25 min since bud initiation (Fig. 1E-1F). In this context, we attempted to simulate possible mechanisms of MTOC clustering within the physiological time limit as determined experimentally by time-lapse microscopy.

In budding yeast *S. cerevisiae*, location for the mitotic division is predetermined even before forming the spindle [14, 22]. The spindle positioning within a confined geometry can often be mapped onto a centering problem across biological systems. For example, force transduction via the active force generators at the cell cortex leads to proper alignment and positioning of the spindle in HeLa cells. It is followed by symmetric division where the cortical pull plays a key role [37]. Similar spindle localization at the cell center is also observed in *C. elegans* embryo during early prophase [38]. In *C. neoformans*, the spindle is formed and stabilizes near the septin ring within the daughter bud [16] (Fig. 1A, Fig. S1, Movie M1). The event is preceded by the migration of the nucleus [16, 39] as determined from the statistics in Fig. S1. Timely nuclear migration into the daughter bud requires directed force generation toward the daughter bud. Since the proper positioning of the spindle and its alignment are crucial determinants of the faithful nuclear segregation [14, 39], the stable spindle localization near the septin ring inside the daughter bud [16] (Fig. 1A, Fig. S1, Movie M1) poses a natural question: what are the force balance conditions necessary for such localization?

In experiments, it is challenging to quantitatively estimate or tweak forces on the spindle in *C. neoformans*. Thus, modeling has been used to supplement/explain the experiments concerning various aspects of spindle mechanics [16, 40–48]. Previously, force balance models have been used to replicate the observed aspects of spindle dynamics [49–52]. The ‘closest to experiment’ approach would be to design simulation using an agent-based model [14, 16, 42, 44, 47, 48, 53], where all objects (e.g., MTs, motors, and other organelles) are simulated as agents having their movements dictated by laws of mechanics.

In this study, we utilized an agent-based model with MTs, MTOCs, and nucleus simulated as agents obeying laws of mechanics within a typical budded cell geometry (Fig. 2A) [14, 16]. Despite the enormous advantages of such an agent-based model, it has certain limitations. In general, the possible number of molecular motors’ combinations in various locations of the spindle at different cell cycle stages is too great (discussed in [54]) and far from clear, particularly in *C. neoformans*. The exact mechanics of the collective motor activity is significantly complex and yet to be well understood. Therefore, for simplicity, we chose not to simulate the molecular motors explicitly. Instead, we opted for a description with constant motor densities where a particular motor-driven force on an MT is proportional to the predetermined motor density. To better understand the spindle positioning, we also set up a simple one-dimensional analytical model with closed-form expressions for various averaged forces (Fig. 2B-2C). By screening these forces, we estimate the plausible force balance for proper spindle positioning in confinement, mimicking a budded cell [36, 49, 55]. These two models not only complement the primary experimental observations of spindle localization in *C. neoformans*, but reasonably qualitatively corroborates with each other. In other words, the benchmarking of the analytical model is supported by the agent-based model and vice versa. Overall, our study highlights that several mechanistic processes can facilitate efficient clustering of MTOCs, either independently or in harness with the other. Furthermore, screening the outputs of *in silico* models uncovers that proper spindle positioning near the septin ring requires MT buckling from the cell cortex in *C. neoformans*.

**FIG. 2.**
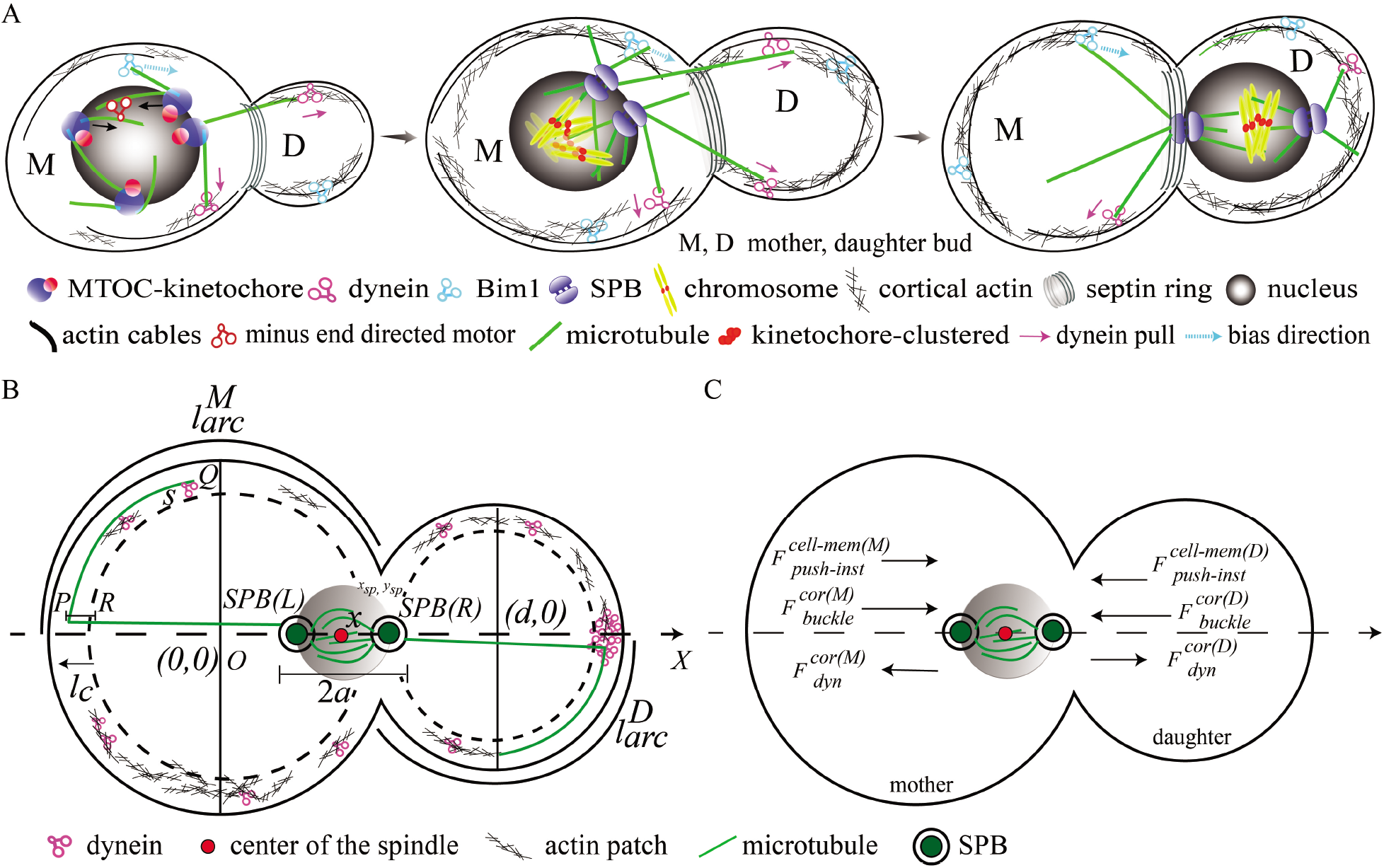
Representative schematic diagrams illustrating the computational and mathematical model. (A) Model schematic depicts the cell cycle stages in *C. neoformans* which are simulated utilizing the computational model. Plausible interaction of cMTs with the cell cortex and on the nuclear envelope are shown. Arrows indicate the direction of forces. (B) Schematic of the one-dimensional mathematical model for spindle positioning in *C. neoformans*. The origin *O*(0, 0) is located at the center of the mother bud, while the center of the daughter bud is located on the axis of symmetry (namely the *X* axis) at (*d*, 0). The instantaneous position of the spindle is denoted by *x* measured from the origin. *l*_*c*_ denotes the cortex’s width, filled with mesh-like actins and dyneins. *SP B*(*L*) and *SP B*(*R*) denote the leftward and rightward SPBs, and the green lines represent microtubules. In the mother bud, the section ‘PR’ is the ‘uncurled’ MT segment in the cortical region, and the segment ‘PQ’ is the ‘curled’ MT segment undergoing sliding within the mother cortex where cortical dyneins pull on both the segments. A similar interaction occurs in the daughter cortex as well. Localized cortical dynein patch in the daughter cortex around the axis of symmetry signifies the differential spatial distribution of dynein in the mother and daughter bud [16]. (C) The direction of the forces acting on the spindle in the mathematical model.

## RESULTS

### Developing *in silico* modeling framework

#### Computational Modeling

We consider the spindle dynamics occurring inside cellular confinement mimicking a budded cell (Fig. 2A). The mother bud size (radius *R*_*M*_) is fixed, whereas the daughter bud (radius *r*_*D*_) grows with time. Prior to the SPB formation, all the MTOCs (spheres of radii *r*_*MT OC*_ *∼* 0.1 *µ*m) are located on the surface of the outer NE (non-deforming spherical nucleus of radius *r*_*nuc*_) and nucleate cMTs. The movement of the MTOCs is constrained on the surface of the NE. At the onset of the simulation, we considered 14 MTOCs [16, 26]. Each MT is modeled as a cylinder of zero thickness with its length regulated by four dynamic instability parameters (Table S2 in the Supporting Material): catastrophe and rescue frequency (*f*_*c*_, *f*_*r*_); growth and shrinkage velocity (*v*_*g*_, *v*_*s*_). After complete clustering of MTOCs into the SPB and SPB duplication (Fig. 1B-1F, 2A), the nMTs from both the SPBs grow inside the chromosomal volume and capture KTs. Meanwhile, the cMTs that reach the cell cortex, majorly experience instantaneous cortical push 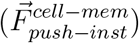, push due to buckling against cell membrane 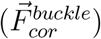, cortical dynein mediated pull 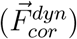 and Bim1 mediated bias towards septin ring 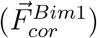 (Fig. 2A, Fig. S2A-S2C). Note that the +TIP protein Kar9 binds to the type 5 myosin Myo2 and the kinesin Kip3. Kar9 plays a significant role in early (pre-anaphase) spindle positioning by forming Bim1–Kar9–Myo2 complexes in *S. cerevisiae* [39, 56–60]. But, it has been reported previously that the *C. neoformans* genome lacks a clear homolog of Kar9 [61]. In the absence of Kar9, we probed for the function of Bim1 (which is a +TIP protein and interacts with Kar9) in generating an effective cortical bias at the plus-ends of the cMTs [16]. Note that the functions of motor and MT-binding proteins are not well worked out in *C. neoformans*. Previous study [62] shows that the partial rescue of the Bim1Δ filament phenotype is consistent with the observations in *S. cerevisiae* and supports the hypothesis that Bim1 acts on the MT cytoskeleton in *C. neoformans*. To that effect, we noted the effective bias towards the septin ring as Bim1 mediated bias/force. Further, the cMTs grazing along the NE experience additional forces (Fig. S2D-S2E); one of them is 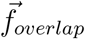 at the MT-MT overlap on NE. The 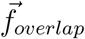 is proportional to the mean number of minus end-directed motors per unit length (*λ*_*ovl*_) at the overlap (Fig. S2E). Similarly, forces between KTs and nMTs have been computed (for details, see the Supporting Material). The instantaneous positions of all the objects (nucleus, MTOCs, SPBs, KTs) are updated by solving corresponding Stokes equations (Eq. S1-S4 in the Supporting Material). The detailed simulation procedure is described in the Supporting Material.

The agent-based stochastic MT dynamics simulation is written in FORTRAN. The data analysis and plotting have been carried out in MATLAB (The MathWorks, Natick, MA) and Gnuplot. A single simulation run takes few minutes of real-computation-time until a stable spindle positioning is attained starting from MTOC clustering (in Intel(R) Xeon(R) CPU having clock speed 2 GHz, RAM 32 GB).

#### Mathematical Modeling

##### Framework

We formulated a mathematical model to study spindle positioning. We frame the mother and the daughter bud as two intersecting circles of radii *R*_*M*_ and *r*_*D*_ respectively (Table S2). The daughter bud’s center is chosen to be ‘*d*’ distance away from the mother’s center. In this framework, the spindle’s spatial movement is allowed along the line joining the centers, regarded as the axis of symmetry. For simplicity, our analytical model is one-dimensional, where all the forces are projected along the axis of symmetry, namely the ‘X’ axis (Fig. 2B-2C). The spindle is chosen to be a rigid rod of length 2*a* with two perfectly bioriented SPBs at its ends lying on the X-axis. We further assume that the cMTs from the SPB facing the mother (daughter) bud cell cortex, namely *SPB*(*L*) (*SPB*(*R*)), interact solely with the mother (daughter) bud cell cortex (Fig. 2B-2C). In the following, we discuss the reasons behind these assumptions. The spindle is nearly aligned with the axis joining the mother and daughter bud center (Fig. 1A). The geometric obstruction imposed by the solid nucleus, the relatively smaller aperture of the bud-neck junction, and the spindle alignment parallel to the axis joining the mother and daughter bud center prevent the cMTs from a SPB to interact with both mother and daughter bud cell cortex in an isotropic manner. The cMTs emanating from the SPB facing the mother (daughter) bud cell cortex majorly interacts with the mother (daughter) cell cortex only (Fig. 2A).

Further, in the model, we considered exponentially decaying spatial dependence of forces, a straightforward and widely used choice [36, 49, 55]. We derived closed-form expressions for various MT-mediated forces (instantaneous cortical push, cortical dynein pull, the force due to MT buckling) on the spindle due to MT-cell cortex interaction. The sum of all these forces accounts for the total force on the spindle. The net force balance on the spindle (rigid rod constituting *SPB*(*L*) and *SPB*(*R*)) determines the spindle position.

Here, the strength of dynein-mediated pulling is regulated by 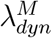 (Eq. S7-S8 in the Supporting Material) and 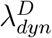 (Eq. S10-S11 in the Supporting Material), cortical pushing by *A*_*M*_ and *A*_*D*_ (Eq. S5 and Eq. S6 in the Supporting Material), average MT length by 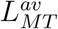 (Eq. S5-S8, Eq. S10-S11, Eq. S13-S14 in the Supporting Material). In the numerical simulation, the corresponding model parameters are adjusted according to the values listed in Table S2 in the Supporting Material. In most cases, the choice of the parameter values is based on the earlier published reports. However, due to the paucity of exact numbers in the literature, a few of the chosen parameter values are optimized for the current study through sensitivity analysis.

##### Utility

In wild-type cells, after SPB separation close to the septin ring, we observe stable spindle localization inside the daughter bud, parallel to the axis joining the center of the mother and daughter bud. This stable localization naturally results from a mechanical force balance within the cellular confinement governed majorly by MTs and molecular motors. Rather than presuming a force balance landscape that supports this localization, we begin by utilizing the simplistic analytical model to screen various forces and determine which combinations of forces are responsible for replicating the qualitative features of the observed phenotypes. Furthermore, closed-form force expressions elucidate the subtle force characteristics of spindle positioning from a largely mechanistic perspective. Taking a cue from this analytical screening of forces, we ‘feed’ the learned force combinations into a detailed agent-based model with explicit simulations of stochastic MT dynamics and MT interactions with the cell cortex to validate the lessons from the analytical model screen. It is natural to ask why we consider the 1-dimensional model when the 3-dimensional model presumably tells us more? The reasons are: (a) to have a first-hand understanding of steady-state characteristics due to force balance when the spindle is placed near the septin ring from a minimal model and (b) to quickly screen the combinatory effect of various forces on spindle positioning. Therefore, we reiterate that it is beyond the scope of the 1-dimensional model to investigate the dynamics of the spindle positioning; we have a 3-dimensional agent-based model to serve that purpose. The mathematical model is described in detail in the Supporting Material. In the following, we present the model investigations exploring MTOC clustering and spindle positioning in *C. neoformans*.

#### Analytical model assumptions and comparisons with the agent-based model

The mathematical model construction is based upon several simplifications described in the following: 1. The model does not account for the process of bi-orientation on the NE. We assume a bi-oriented configuration of two SPBs throughout (Fig. 2B-2C). In the agent-based model, after SPB duplication, the process of SPB biorientation has been taken into account. 2. In the analytical model, the spindle length is fixed independent of its position along the axis of symmetry. In the agent-based simulations, spindle length is determined by the forces acting on nMTs and cMTs. 3. In the 1-dimensional analytical model, the motion of the nucleus/spindle is strictly confined along the X-axis. We only emphasize the translational motion of the spindle governed by a set of MT-mediated forces. The rotational degrees of freedom responsible for the spindle orientation is not considered in the model for simplicity. The spindle is always laid upon the X-axis, with its orientation being parallel to the symmetry axis (X-axis). Allowing the spindle to move only along the X-axis accounts for the net cancellation of forces along the transverse direction since MT nucleation is considered isotropic in all directions. In the agent-based model, simulated on a 3-dimensional geometry, no such restrictions have been imposed (Fig. 2A-2C). 4. The analytical model does not include stochastic fluctuations originating from the randomness in MT dynamics or its interaction with the cell cortex. The closed-form expressions of the governing forces are evaluated in a time-independent quasi-equilibrium configuration. In the agent-based model, a significant source of stochastic fluctuations is the MT dynamic instability [1, 63–65] and attachment/detachment of molecular motors. 5. For simplicity, we have not considered a load-dependent variation of average MT length in the analytical model. However, in the agent-based simulation, load-dependent modulation of the MT dynamic instability parameters, growth velocity (*v*_*g*_), and catastrophe frequency (*f*_*c*_) have been taken into account.

### Clustering of MTOCs progresses via redundant pathways

We have shown previously that all KTs cluster to form a punctum in *∼* 25 mins [16]. We further demonstrated that KTs colocalize with MTOCs [20]. Now, we validated the time required for MTOCs to coalesce into an SPB by calibrating the clustering time since bud initiation (time set to zero), in terms of the budding index (Fig. 1E-1F). Since the bud initiation, the daughter bud grows with time in a reasonably uniform manner over a large population of cells. As the bud growth rate is roughly consistent, the budding index can be thought of as an intrinsic ‘clock’ of a budded cell (Fig. 1E). The number of MTOCs in cells having similar budding indices (implying that the cells are at the same cell cycle stage) was counted and plotted with ‘time’ (calibrated from the budding index) to estimate the MTOC clustering time (Fig. 1F). Note that, while measuring the absolute time, rather than calculating it from the bud-size, would be more appropriate for studying the course of MTOC clustering, the present manuscript aimed to understand the plausible mechanisms of MTOC clustering from the estimate of average clustering time. Since unclustered and clustered MTOCs occur at two endpoints of the measurements, the time difference between these events estimated from several bud-size measurements was consistent.

We observed that the timescale for KT clustering [16] and MTOC clustering (Fig. 1F) are similar (*∼* 25 min) and in good synchrony with each other (Fig. 1A-1D). Although electron microscopy suggested as many as 14-16 MTOCs [26], with a limited resolution, we could identify a maximum of 8-10 MTOCs in small budded cells. Next, we address how all the MTOCs cluster into a single group within this specific timescale using an agent-based mechanistic model (Fig. 2A, Fig. 3A, Fig. S2). First, we investigated the MTOC clustering process via MT-driven ‘search and capture’ on the NE without the cortical interactions (Fig. 3A, Fig. S2D-S2E). First, we determined the time required for all the MTOCs to cluster if cMTs growing out of MTOCs slide along the NE and directly capture the MTOCs (Fig. S2D). We considered that in the ‘direct search and capture’ mechanism (Fig. 3A, Fig. S2D), the MTs contributing to the clustering process remain confined to the NE while growing in a random direction. As the ‘searcher’ MT tip nucleated from an MTOC grazes along the NE and ‘captures’ another MTOC (‘target’), the MTOCs move toward each other along the MT until they fuse (Fig. 3A, Fig. S2D). The movement of the MTOCs is strictly restricted to the NE. In this context, we assumed a fixed number of grazing cMTs per MTOC (*∼* 3-4 cMTs per MTOC). We found that, in the explored parameter regime, the timescale for complete clustering achieved by this ‘direct search and capture’ pathway is significantly large, surpassing the relevant mitotic timescales. The MTOCs take *∼* 1.5 h to cluster entirely, that too, in *∼* 10 % of the cell population only. The MTOCs fail to cluster into a single focus in the rest of the cell population even after *∼* 4-5 hr. Note that, in the current *in silico* study, the term ‘cell population’ alludes to the total number of independent simulation runs started with uncorrelated initial configurations. In this particular context, we performed many separate simulation runs, and in 10 % of these simulations, clustering less than 1.5 h was observed. Thus, it is unlikely to be an ‘efficient’ pathway for MTOC clustering in the currently explored parameter regime. However, the possibility that the same pathway functioning efficiently in combination with other pathways cannot be ruled out.

**FIG. 3.**
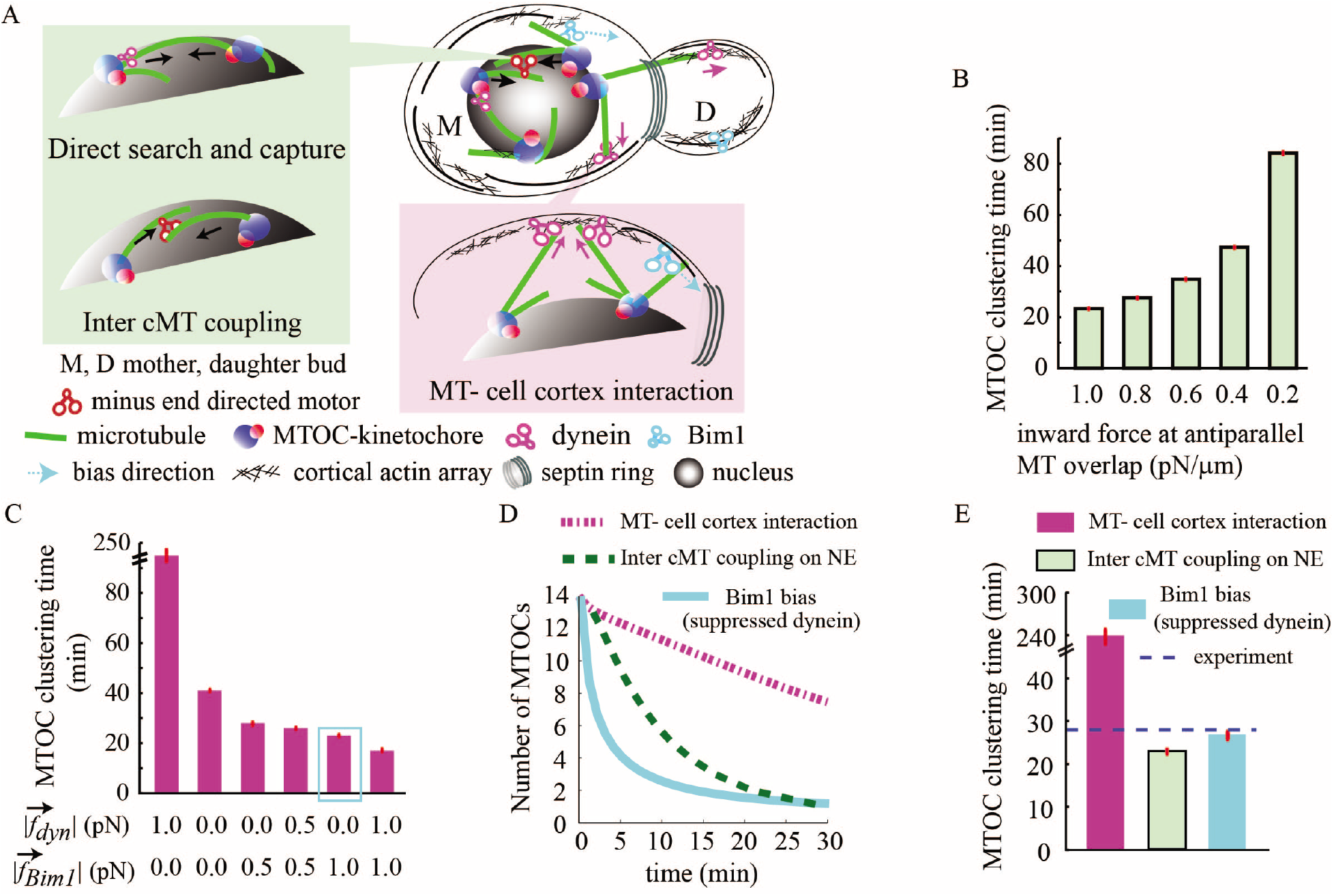
Cooperative interactions among MTs and motors govern MTOC clustering. (A) Model schematic depicting MT-mediated ‘search and capture’ processes and forces required for the clustering of MTOCs. Arrows indicate the direction of forces. Plausible clustering mechanisms illustrated are: a growing cMT directly capturing an MTOC (‘direct search and capture’), sliding of antiparallel cMTs via crosslinking minus end-directed motors on the NE (inter cMT coupling at the NE), and cMT plus-ends interacting with cell cortex (MT-cortex interaction). (B) Clustering time in the sole presence of inter cMT coupling at the NE increases as the minus-ended motor-generated inward force at MT-MT overlap decreases. The MT number is fixed at 4 per MTOC. (C) Clustering solely via MT-cell cortex interaction. The clustering is faster when Bim1 bias is enhanced. 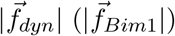 represents the magnitude of the force generated by a single dynein (Bim1) on each MT at mother cortex. (D) Time progression of MTOC clustering in different mechanisms. The first/fifth bar in Fig. 3C denotes the parameter values corresponding to the magenta/cyan curve in Fig. 3D, respectively. (E) Comparison between MTOC clustering timescales via MT-cell cortex interaction, inter cMT coupling on NE and Bim1 bias with suppressed dynein activity at cell cortex 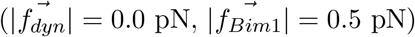. In all figures, sample size n*>*2000 for simulation and red bars indicate SEM (wherever shown).

Next, we introduced antiparallel sliding amongst the grazing cMTs on the NE (inter cMT coupling, Fig. 3A, Fig. S2E). Inward sliding of the overlapping cMTs from two different MTOCs can be steered by crosslinking minus end-directed motors [66]. The crosslinking activity of minus end-directed motors acting at the MT-MT overlap brings the engaged MTOCs towards each other. In the earlier scenario, a ‘searcher’ MT has to grow and capture another MTOC. However, in the present context, a ‘searcher’ MT can grow and capture another MT segment so that the minus end-directed motors can crosslink them and initiate antiparallel sliding. Adding the minus end-directed motor crosslinking and sliding on the NE facilitate the timely clustering; clustering of all the MTOCs (*∼* 100 %) into a single object happens within *∼* 23 min (Fig. 3B, Movie M2). As expected, when the inward force between the MTOCs due to minus-ended motors diminishes, the net clustering time increases (Fig. 3B). Additional parameter sensitivity analysis for this clustering mechanism is described in the Supporting Material, Fig. S3.

In essence, the modeling results propose the ‘minus end directed motor mediated antiparallel cMT sliding on NE’ as a plausible mechanism of MTOC clustering within experimentally observed time window. A possible candidate for this process is minus-ended dynein. Hence, we experimentally tested whether dyneins are present on NE during MTOC clustering. To this note, the dual-color imaging of Dyn1 and MTOCs will strengthen the case of theoretically proposed minus-end directed motor-mediated coupling and sliding of antiparallel cMTs. However, it is technically challenging to address this where we have to construct a strain where either Dyn1 and MTOCs or Dyn1 and MTs are tagged simultaneously due to the limited availability of auxotrophic markers in this organism. Instead, we expressed GFP-tagged Dyn1 and studied its relative localization with respect to the nuclear periphery protein Sad1-mCherry. We have shown previously that Sad1 localizes to the nuclear periphery and the dot-like signal of Sad1 is only restricted to the NE [20]. Sad1 also connects kinetochores to the MTs/MTOCs and facilitates clustering. We observed multiple dynein puncta surrounding the nuclear periphery (Fig. S12). Occasionally, dynein partially colocalizes with Sad1 (Fig. S12), indicating that Dyn1 might be involved in the sliding of cMTs. Since some signals from Dyn1 were weakly detected, we could not quantify the exact number of dynein molecules in this strain. Note that, at this stage, the presence of any other minus-end directed motor candidate (besides dynein) cannot be ruled out (therefore, testing the role of other minus-end directed motor candidates e.g kinesin-14 in MTOC clustering is a worthy future venture).

Next, in the model, we explore the role of cMT-cell cortex interaction leading to the clustering of the MTOCs. Two major cortex-based forces on the cMTs are: (a) net pull on the MT segments sliding inside the cortex via the cortical dyneins towards the cortex; (b) Bim1 mediated force on the cMTs (sliding +ve end of the cMTs along the cortex) having a directional preference towards the septin ring (MT-cell cortex interaction, Fig. 3A, Fig. S2A-S2B). We found that the exclusive activity of dynein-mediated MT-cell cortex interaction leads to nearly complete clustering of the MTOCs, but the time estimated (*∼* 240 min) is way too high (Fig. 3C, Movie M3). However, a decrease in the cortical pull and/or subsequent increase in the Bim1 mediated bias reduces the clustering time scale to ≲ 50 min (Fig. 3C, 3D, Movie M4). We find that the dynein mediated pull on the MTOCs via cMTs acts in random directions, whereas the Bim1 mediated bias is directed towards the septin ring. Thus dynein dominated cortical pull suppresses the effective Bim1-bias and delays the clustering. When dyneins are suppressed, due to Bim1-bias all the MTOCs are drifted towards the septin ring along the NE and cluster rapidly. Note that if the clustering time via a mechanism largely exceeds the physiological time limit estimated from the live-cell imaging, the sole presence of the mechanism under consideration may be discarded. In reality, in the context of a dividing cell, significantly delayed clustering by any ‘inefficient’ mechanism is unlikely to happen.

The time scales for complete MTOC clustering appears to be similar and close to the physiological time limit when (a) MTOCs aggregate due to antiparallel sliding of the grazing MTs on the NE via minus ended motors and (b) MTOCs aggregate due to diminished cortical dynein pull concomitantly with enhanced Bim1-bias at the cortex (Fig. 3D-3E, Table S3 in the Supporting Material). This highlights a crucial physical possibility that the clustering mechanisms may be redundant, i.e., the mechanisms either act in liaison or independently

The complete clustering of MTOCs into a single body marks the formation of the mature SPB followed by SPB duplication. Subsequently, the duplicated SPBs separate into a bioriented configuration that initiates spindle formation. In our earlier and present studies, it has been shown that the mitotic spindle stabilizes inside the daughter bud close to the septin ring in *C. neoformans* [16] (Fig. 1A, Movie M1, Fig. 2A).

Therefore, an obvious precursor to the spindle positioning inside the daughter bud is nuclear migration from the mother bud to the daughter. From a mechanistic standpoint, this is facilitated by cortical interaction of cMTs (Fig. 2A, Fig. S2A-S2C) [14, 16]. Interestingly, cortical interaction is also an important mode of MTOC clustering, as we have already shown. Therefore, MTOC clustering, nuclear migration, and subsequent positioning of the spindle are the sequence of crucial mitotic events which are unlikely to be mutually exclusive. The key elements necessary for orchestrating the pre-anaphase MTOC clustering and nuclear migration are (a) MT integrity, (b) spatiotemporal localization and activity of Bim1 and dynein [16]. Bim1 and dynein majorly generate the forces which are transduced via cMTs to the MTOCs (later SPBs) and the nucleus.

Previously, we showed that Aurora B kinase Ipl1 in *C. neoformans* plays a pivotal role in (a) maintaining MT stability and (b) regulating Bim1/dynein activity or localization [16]. The depletion of Ipl1 interrupts both MT stability and Bim1/dynein activity. We observed that delayed KT clustering correlates with delayed nuclear migration among heterogeneous phenotypes of defective KT clustering and impaired nuclear migration in Ipl1 depleted cells. Since the time scales of MTOC clustering and KT clustering are similar and occur concomitantly [16] (Fig. 1B-1F), it is likely that delay in MTOC clustering also leads to a delay in nuclear migration.

Taking cues from these experimental observations, we asked the following questions in the *in silico* model: (a) What happens to the nuclear migration and concomitantly progressing MTOC clustering when cortical Bim1 and dynein densities are varied? Note that, ‘density’ of molecular motors and ‘force’ applied by them are proportional in our model. Thus, variations in the Bim1 and dynein densities allude to equivalent variations in the respective forces. (b) Does impaired/delayed clustering lead to faulty migration? (c) What happens to the spindle positioning and spindle orientation when MTOC clustering is delayed?

To address these questions, we carried out sensitivity analysis over the following parameters: Bim1 bias force per MT, the number of MT per MTOC, and average cMT length (Fig. 4A-4C, Fig. S5A-S5B). We find that the time required for MTOC clustering and nuclear migration are reasonably correlated. If the MTOC clustering is delayed/accelerated, nuclear migration also follows a proportional timescale (Fig. 4A-4C) [16].

**FIG. 4.**
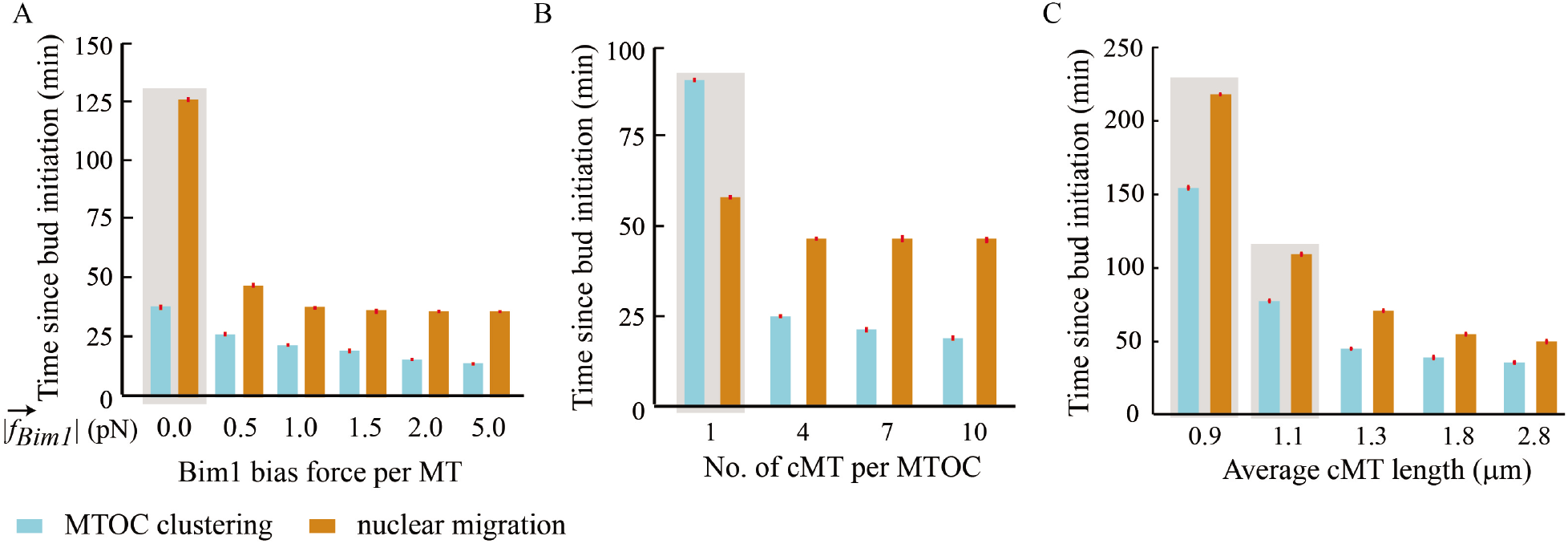
MTOC clustering and nuclear migration: sensitivity to model parameters. (A-C) Time required for complete MTOC clustering and proper nuclear migration to daughter bud when Bim1 bias force per MT (A), number of MT per MTOC (B) and average cMT length (C) are varied. For MTOC clustering, solely the mechanism of “MT-cell cortex interaction with diminished cortical pull and enhanced Bim1 bias” is considered. In all figures, sample size n*>*2000 for simulation and red bars indicate SEM (wherever shown).

Furthermore, a close inspection of Fig. 1A indicates that the mother bud also appears to grow over time, albeit at a much slower speed. We numerically assessed whether this small change in the mother bud size with time affects the time scales of MTOC clustering and nuclear migration. To this note, we have performed several simulations where the mother bud also grows with time in harmony with the daughter bud. In our simulations, we begin with a particular radius of the mother bud. The radius increases with time until it reaches a fixed, predetermined value. Quite evidently, in the simulations, the mother bud’s growth rate is considered to be much slower than that of the daughter bud. In the currently explored parameter regime, we do not observe any significant change in the time scales of MTOC clustering and nuclear migration when the feature of mother bud growth over time is introduced in the model (Fig. S11). Therefore, it is reasonable to keep the mother bud at a fixed size throughout the simulations (unless mentioned otherwise).

Interestingly, the *in silico* analysis further suggests that the delay in nuclear migration and MTOC clustering do not affect the spindle positioning or the spindle orientation in the currently explored parameter regime (Fig. S5A-S5B). Variations in Bim1 bias force per MT and the number of MT per MTOC incur variations in the nuclear migration time (and MTOC clustering time) within a reasonable range (Fig. 4A-4C). But the spindle positioning within the daughter bud (and spindle orientation) turns out to be independent of the variations in Bim1 bias force per MT and number of MT per MTOC (Fig. S5A-S5B). In other words, irrespective of whether the nuclear migration is delayed or accelerated, the stable spindle position (and spindle orientation) remain unchanged within this specific cellular geometry (Fig. S5A-S5B). Experimental quantification of various spindle attributes in wild-type and upon molecular perturbations (Fig. S1, Fig. S4A-S4C, Fig. S5C-S5E, Fig. S6) are discussed in the following sections. Additionally, the quantification protocols for the experimental data on nuclear migration are discussed in the Supporting Material in detail.

To understand the stable spindle localization, we propose a simple analytical model that estimates net force balance on the spindle (See the Supporting Material and Fig. 2B-2C). We considered closed-form expressions for various MT-based forces and examined possibilities of the spindle localization under the combinatory effect of these forces (Fig. 2, Fig. 5, Fig. S7).

**FIG. 5.**
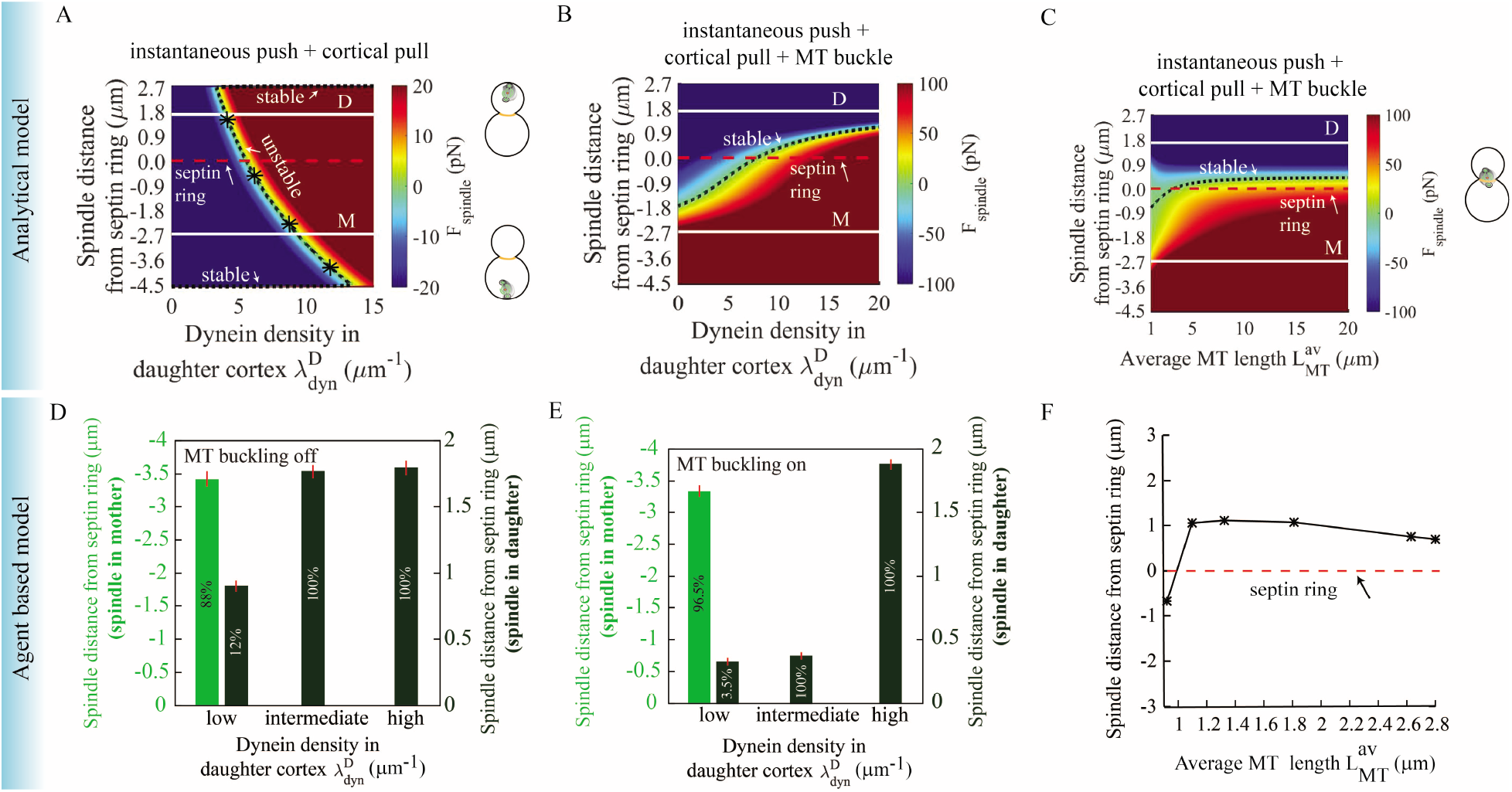
The mechanistic models of force balance explain spindle positioning in *C. neoformans*. (A) Spindle stably collapses onto the cell cortex as dynein density in the daughter cortex 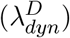 is varied. Dynein density in the mother cortex 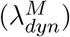 is fixed and instantaneous cortical push is present in both mother and daughter cortex. Color bars represent the net force on the spindle and white solid lines denote the center of the daughter and mother buds. (B-C) MT buckling is present combined with instantaneous push and cortical pull. Increasing dynein density in the daughter cortex 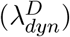 steadily brings the spindle into the daughter, localizing near the septin ring (B). Spindle position varies depending on the average MT length 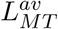 (D-E) Spindle distance from septin ring in the absence (D) or presence (E) of MT buckling at the cortex as observed in the agent-based simulation. The values within the bars indicate the percentage of spindles in mother or daughter across the cell population. (F) Spindle position depending on the variation in average MT length 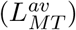 as observed in the agent-based simulation when instantaneous cortical push, dynein pull, and MT buckling are acting in tandem. In plots A-F, - ve/+ve distance refers to the spindle in mother (M)/daughter (D) bud.

### Balance of cellular mechanical forces guides the spindle position

A spindle is constantly acted upon by molecular forces arising from the interactions of its components with the surrounding [14, 16, 36, 43, 48, 67–69], e.g., cMTs interacting with the cell cortex via dynein generates a pull on the nucleus toward the cell periphery; similarly, cMTs buckling in contact with the cell membrane pushes the nucleus away from the membrane. These interactions are plausible in both mother and daughter bud cell cortices. Besides, interactions of cMTs with the septin ring and the cortical actin cables generate an effective bias translating the plus ends of the cMTs toward the daughter bud. Since these forces are spatially inhomogeneous and rapidly varying with time, it is logical to ask how the nucleus and the spindle attain steady positions before the chromosomal segregation.

We formulate an analytical template in one-dimension that accommodates key MT-mediated forces originating from the mother and daughter cortices acting on the SPBs (Fig. 2B-2C). The mechanistic force balance landscape emerging from the mathematical model allows us to carry out sensitivity analysis across a broad parameter regime constituting a myriad of forces with different characteristics (e.g., instantaneous pushing and buckling of the cMTs, dynein mediated pulling, etc.). From experiments and agent-based simulation, we observe that the average spindle length is *∼* 1.5 *µ*m in wild-type (Fig. S5C) and the neck to spindle distance is *∼* 1 *µ*m while the spindle is in the daughter bud (Fig. S5D). Also, the spindle lies almost parallel to the axis of symmetry (orientation angle *∼* 15 degree), joining the centers of the mother-daughter bud (Fig. S5E).

To test how sensitive is the spindle positioning to the cMT-cortex based forces, we introduced instantaneous push and dynein pull from both the mother and daughter cortex. Then, we varied the daughter cortex’s dynein density, keeping the rest of the parameters fixed at base values denoted in Table S2. If the daughter cortex’s dynein pull is too strong, the spindle collapses onto the daughter cortex. In the other limit, the spindle collapses onto the mother cortex, when the mother cortex’s cortical pull overpowers the pull from the daughter (Fig. 5A). Thus, in the absence of MT buckling, the spindle localizes at the cortex. Note that the term ‘collapse’ means the spindle is dragged too close to the cell cortex.

Adding the force due to MT buckling (with two other forces: cortical pull and instantaneous push) restores the stable spindle position near the septin ring (Fig. 5B-5C). The spindle robustly maintains its position inside the daughter bud, close to the septin ring upon considerable variation in the average cMT length (Fig. 5C, Table S4 in the Supporting Material).

We also tested the spindle positioning in the absence and presence of MT buckling, exploiting the three-dimensional agent-based computational model (Fig. 5D-5E). We find that a moderate dynein pull is crucial for the spindle to migrate into the daughter. In the absence of MT buckling, the spindle localizes deep inside the daughter cortex for moderate and robust dynein pull from the daughter (Fig. 5D). For small dynein pull from the daughter cortex, the nucleus is retained in the mother bud (Fig. 5D). In the experiment, we observe that upon dynein depletion, most of the cells have nuclei retained in the mother bud (Fig. S6A), which agrees with the model outcome (Fig. 5D). Deletion of Bim1 also leads to faulty nuclear migration (Fig. 4A, Fig. S6B). While acquiring the statistics in Fig. S1, Fig. S6A-S6B, we have considered only large budded cells having budding indices 0.65 and above. Since the budding index is an indicator of ‘time’ (Fig. 1E), higher budding indices refer to the cells in which a considerable amount of time has elapsed after bud initiation. In most wild-type cells with budding indices above 0.65, the nuclei are settled in the daughter bud (Fig. S1). Upon dynein depletion/Bim1 deletion, we observe more cells with budding indices above 0.65 have nuclei retained in the mother bud (Fig. S6A-S6B) because in those cells, the nuclear migration is possibly delayed/impaired. The same conclusion can be gleaned from the statistics of the budding indices of unsegregated cells in Fig. S4A. Next, to understand if dynein and Bim1 both are required for nuclear migration, we generated a strain where dynein is depleted in the absence of Bim1 (GAL7-DYN1 bim1Δ) in the histone GFP-H4 tagged background. We quantified the percentages of cells exhibiting abnormal nuclear segregation or no segregation of nuclear masses in bim1Δ, GAL7-DYN1 and/or GAL7-DYN1 bim1Δ strains and compared them with the control wild-type H99 strain (Fig. S4B). We observed a significant increase in the number of unbudded and large budded bi-nucleated cells upon simultaneous depletion of dynein and Bim1 compared to the control (Fig. S4C). This result indicates that both Bim1 and dynein are required for nuclear migration where simultaneous depletion of both proteins results in delayed nuclear migration due to which the nucleus divides inside the mother bud itself in the majority of GAL7-DYN1 bim1Δ cells. A note on the quantification of the experimental data on nuclear migration is included in the Supporting Material.

With MT buckling turned ‘on’, the spindle settles inside the daughter bud and stabilizes close to the septin ring in the currently explored parameter regime (Fig. 5E). An exciting feature of the spindle noticed via 3-dimensional agent-based simulations is its proper orientation parallel to the mother-daughter axis, similar to the microscopic images in Fig. 1A. While, in the absence of MT buckling (Fig. 5D), the spindle inside the daughter appears to be randomly oriented relative to the axis. Due to these differences in spindle orientation, the absolute position of the spindles in the absence and presence of MT buckling (Fig. 5D-5E) may not be compared explicitly. Nevertheless, based on Fig. 5D-5E we can say that the daughter cortex’s high dynein density pulls the spindle deep inside the daughter bud. Unlike the 1-dimensional analytical model, we do not observe any spindle collapse in the 3-dimensional agent-based model. The plausible reasons are (a) difference in the dimensionality between the two models, (b) stochastic fluctuations in the agent-based model compared to no fluctuation in the 1-dimensional analytical model, (c) dynamic changes in spindle orientation in the absence of MT buckling as discussed above in 3-dimensional simulation, etc. as opposed to parallel orientation in 1-dimensional model.

As the mechanics of spindle positioning is largely MT mediated [16, 36, 39], we tested the dependence of spindle positioning on the average MT length. From the analytical and the agent-based model analysis, we found that when the average MT length is ‘too short’ (*∼* 1 *µ*m), the spindle is retained in the mother (Fig. 5C, Fig. 5F). In other words, the nucleus is unable to migrate to the daughter bud. This result corroborates with our experimental observations (Fig. S6C-S6D). We observe spindle localization inside the daughter bud near the septin ring in wild-type cells (Fig. 1A, Fig. S6C, Movie M1). But in cells treated with nocodazole, the MT integrity is severely compromised (Fig. S6D). In that scenario, we do not observe any spindle localization in the daughter bud (Fig. S6D).

We further observe that in the presence of MT buckling, the variation in average MT length above a threshold does not affect the spindle localization inside the daughter bud (Fig. 5C, Fig. 5F). As a plausible explanation, we argue that when the average cMT length is above a threshold, a few cMTs always reach the cortex. At the cortex, the interaction of MT and molecular motors transduce pN order forces, which is sufficient to stabilize the spindle at the proper location. Nevertheless, the force due to a MT’s buckling is considerably large compared to other relevant forces (Table S2). Therefore, the buckling transition of only a few cMTs from the cortex produces sufficient forces dominating the spindle localization. The position of the spindle remains unaffected because (a) the buckling mediated forces pointing away from the mother and daughter bud cell cortex are oppositely directed (Fig. 2C), (b) number of MTs undergoing buckling at mother and daughter bud cell cortex are similar on average. Therefore, in the buckling-dominated landscape, the forces directed away from the mother and daughter cell cortex nullify each other near the septin ring.

To conclude, the spindle distance from the septin ring measured in the experiment (Fig. S5E) and estimated from the analytical (Fig. 5C) and computational models (Fig. 5F) reasonably agree with each other. Additional spindle characteristics predicted by the model are summarized in the Supporting Material, Fig. S7, and Table S4. Furthermore, we have also performed additional agent-based simulations to explore how varying dynein density profiles in mother and daughter cell cortex impact the timescales of MTOC clustering and nuclear migration (Fig. S13). The results are discussed in the Supporting Material in detail.

In the agent-based computational model described above, the stochastic effects come into the picture through the MT dynamic instability and MT-cell cortex interaction. While this is a crucial source of stochasticity, other sources, including attachment-detachment of molecular motors (occurring at much shorter time scales) could significantly influence the earlier model outcomes. In view of that, we introduced the feature of stochastic MT-dynein attachment-detachment in the existing model. A similar stochastic character is introduced in the context of Bim1 as well. The description of the model framework with stochastic attachment-detachment and detailed results are reported in the Supporting Material (Fig. S8-S10).

## DISCUSSION

Our theoretical approach combines (a) an agent-based computational model of MT ‘search and capture’, and (b) an analytical model to explore the mitotic events in *C. neoformans*. With the help of the computational model, we examined the role of MT ‘search and capture’ during MTOC clustering, nuclear migration and spindle localization. Using the analytical model, we screened various combinations of the MT-based mechanistic forces and found plausible mechanical force balance conditions that may orchestrate proper spindle positioning. We reached the following conclusions (Fig. 6).

**FIG. 6.**
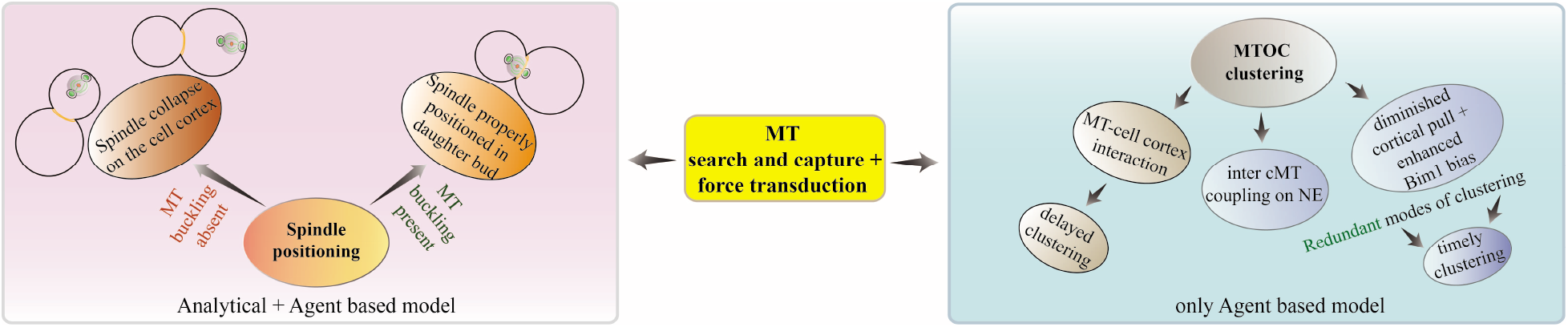
Summary of model outcomes describing mechanistic aspects of MTOC clustering and spindle positioning in *C. neoformans*. The agent-based model depicts redundant mechanisms for the timely clustering of MTOCs. Inter cMT coupling at NE and/or cortical interaction of MTs with suppressed dynein pull and enhanced Bim1 bias may facilitate timely clustering, either independently or in unison. Furthermore, the analytical model supported by agent-based simulations suggests that proper spindle positioning in the daughter bud near the septin ring requires MT buckling from the cell cortex.

The emergent time scales for MTOC clustering are similar in two scenarios; the design principles of which are based on a) effective inter-MTOC attraction due to minus end-directed motor crosslinking between grazing antiparallel MTs on the NE and b) effective drift of all the MTOCs towards the septin ring due to Bim1 mediated cortical bias with diminished dynein pull from the cortex. In the first scenario, the clustering time scales down with the minus-end motor mediated inward force at the MT-MT overlap. In the latter one, the clustering time scales up with the net cortical dynein pull on the MTs. From a model perspective, the comparable timescales of these two independent clustering mechanisms indicate a possible redundancy between the two mechanisms (Fig. 3D-3E).

During mitosis, cells self-organize the spindle to facilitate nuclear division, and for a faithful division, it is essential to have a bipolar spindle. The reason is that the sister chromatids require to partition into two opposite poles and settle in two daughter progeny. In mammalian cells, the prevalence of more than two centrosomes at the beginning of mitosis can generally be linked to multipolar spindle formation [36]. Multipolar spindles and the presence of more than two centrosomes/MTOCs often lead to chromosome instability, aneuploidy, erroneous attachments of chromosomes, and various other defects of chromosome segregation hindering faithful division. Recent studies highlight an interesting pathway of bypassing these defects where multiple centrosomes/MTOCs assemble into two groups/poles, making way for bipolar spindle formation in multi-centrosome/multi-MTOC cells. The process of converging multiple centrosomes into two poles during spindle formation is often termed ‘centrosome clustering’ [36, 70–72].

However, in the context of MTOC clustering in *C. neoformans*, the pathway for grouping the MTOCs into two poles turns out to be different. In *C. neoformans* we observe that MTOCs dynamically move toward each other along the outer NE’s surface and eventually fuse, forming the SPB. During this process, KTs are found to colocalize with the MTOCs (Fig. 1B-1D) [20]. Unlike centrosome clustering resulting in two independent poles, MTOCs cluster into a single pole. The clustered pole, namely the SPB, subsequently duplicates into two SPBs paving the way for bipolar spindle formation.

Our modeling results indicate that Bim1 mediated cortical bias plays a crucial role in MTOC clustering and nuclear migration. But, how Bim1 (EB1) participates in nuclear migration process in the absence of Kar9/Myo2 is still unclear. A previous study [61] reported that *C. neoformans* lacks a Kar9 homolog. However, we have now identified the putative homolog of Kar9 in *C. neoformans* having a conserved LXXPTPh motif required for its interaction with Bim1. Our preliminary analysis of the localization of the putative Kar9 protein suggests that upon over-expression, it exhibits MT-like localization, indicating it could possibly be a true homolog of Kar9. However, the localization of Kar9 expressed under the native promoter is yet to be analyzed. At present, our knowledge regarding its function in the movement of the spindle towards the septin ring is also limited, and this is currently being investigated as an independent study. In addition to this, it would be interesting to experimentally probe the predicted Bim1 biased (suppressed dynein) regime (for MTOC clustering time) by adding dynein inhibitors at different concentrations in a time-lapse experiment. Telomere-localized SUN and KASH proteins in association with MT-associated motor proteins (dynein and kinesins) have been shown to facilitate telomere clustering in fission yeast [73]. In light of this and our results on role dynein and SUN-domain protein Sad1 in KT clustering independently, we propose that a similar model could also apply to MTOC/KT clustering in this organism, and we are interested in exploring this in detail in the future.

Previous studies outline that antiparallel MT bundler Ase1 influences the spindle assembly process [74] and chromosome alignment [75, 76]. The role of Ase1/PRC1 in directed force generation at the MT-MT overlaps was also studied previously [77, 78]. One of our proposed MTOC clustering mechanisms entails minus-end directed motor mediated sliding at the antiparallel MT-MT overlaps. Therefore, it would be interesting to investigate Ase1/PRC1 in the antiparallel MT-MT overlaps. In our agent-based model, we have ignored this for simplicity. Besides, in the current experimental setup, it is also difficult to explore the role of Ase1/PRC1 in MTOC clustering. An experimental investigation in this direction, supported by theoretical modeling, would be a worthy future endeavor.

By screening various MT-based forces, our analytical model qualitatively highlights the mechanical requirements for proper spindle positioning. It has been suggested that the plane of mitotic division is defined by this stable positioning of the spindle inside the daughter bud with spindle orientation almost parallel to the axis joining the centers of the mother and daughter buds[14, 16, 79]. Our model shows that in the presence of two opposing forces, instantaneous cortical push and dynein pull (no MT buckling), the spindle collapses onto the cortex. Additional force due to MT buckling at the mother and daughter cortex is essential to restore the spindle’s stable positioning near the septin ring. A key reason is the force due to buckling that scales with the inverse square of the MT length. Thus, when the spindle is close to the cortex, buckling MTs strongly push it away, preventing the collapse. The analytical model also shows several interesting positions of the spindle which are unstable. However, the 1-dimension model can be extended to higher dimensions, which would include several important characteristic degrees of freedom (e.g., the orientation of the spindle, the angular displacement of the SPBs, etc.) in the analysis as well.

To examine the physical basis and the consistency of the spindle positioning attributes, the analytical model results are compared with the agent-based computational model. One of the primary differences between the analytical and computational models is the stochastic effects. The agent-based model entails stochastic fluctuations stemming from the MT dynamic instability, a finite number of MTs, motor activity, etc. In contrast, the analytical model does not contain stochastic fluctuations and temporal degrees of freedom. Possibly, due to intrinsic fluctuations in the computational model, we do not observe any unstable spindle position.

In the agent-based model, a major source of fluctuations is the stochastic MT dynamics manifesting the random switch between the states of polymerization, and depolymerization [1, 63–65]. In the mathematical model, the stochastic MT dynamics is not explicitly considered; the MT length distribution is chosen to be exponential. In the agent-based model, stochasticity also comes into the picture through MT interaction with the cortex/cell membrane. When an MT elongates to the cortical region, the consequence is a probabilistic choice of the MT a) undergoing catastrophe, b) sliding along the cortex, and c) buckling with the tip hinged/pivoted at the cell membrane. However, stochasticity stemming from the explicit binding or unbinding of the motors on MTs, collective ‘walking’ of various groups of motors on the filaments, and associated modulations in force transmission via the MTs are not explicitly incorporated in the coarse-grained model. The stochastic effects arising from the thermal fluctuation at the molecular level and interactions of the ‘objects’ with embedding viscous medium (e.g., the cytoplasm, the nucleoplasm) are not accounted for in the model.

Previously, the mechanics of MT aster positioning has been studied with considerable details in various contexts [80–82]. It has been suggested that both pushing and pulling forces (primarily relative to the cell boundary) govern the force balance landscape necessary for MT aster positioning. The mechanical conditions for a stable centering position for an MT aster within a confined geometry are: (a) pushing forces in tandem with MT catastrophe in the cytoplasm before the MTs hit the cell boundary; (b) pulling forces in tandem with MT sliding/slippage at cell cortex [82]. In the context of *C. neoformans*, we observe that the stable spindle positioning near the septin ring is strongly supported by the pushing forces (including buckling), which conforms with the condition (a). However, in the currently explored parameter regime, we do not observe stable spindle positioning near the septin ring when the cortical pull is acting alone (Fig. S7B) or in combination with the instantaneous push (Fig. 5A, Fig. S7B, Fig. S7D-S7E). In special cases when (a) the average MT length is significantly large compared to the cell size, (b) there are many cMTs or (c) cMT segments that keep sliding inside the cortex, the cortical pull may facilitate stable spindle positioning near the septin ring. Nevertheless, experiments indicate that the cMT number per SPB is limited and the average MT length is of the order of the cell size in *C. neoformans* [14, 16] restricting the currently explored *in silico* parameter regime.

It is important to mention that the analytical model is one-dimensional with simplified force expressions. Exploring the possibility of pulling mediated ‘stable centering’ in the current context is possibly beyond the scope of the analytical model presented here. Taking cues from other mechanistic models of MT aster positioning [80–83], a detailed mathematical model can be designed to elucidate the mechanics of spindle positioning further.

In summary, in the current study, we constructed a phenomenology-based model that consistently explores spindle assembly aspects during mitotic cell division. Using the model, we have elucidated several important features of MTOC clustering, nuclear migration, and spindle positioning, largely from a mechanistic point of view (Fig. 6). While attempting to understand these processes in the light of dominant microtubule-based forces (e.g., buckling or pulling from the cortex), we ignored weaker forces that may arise from sources *viz*, actin turnover, other molecular motors, thermal fluctuations, etc. Nevertheless, all these forces are important and a more realistic model should incorporate them in our future projects. Note that the attributes of the dynamic processes involved in the mitotic cell cycle differ widely across different cell types. Different organisms evolved (with increasing complexity at the molecular level) to self-engineer the process of cell division. Thus, investigating the mechanistic principles of the cell division across different organisms using a systems biology-based approach in collaboration with molecular biology experiments stands as the subject of future research.

## MATERIALS AND METHODS

### Experimental protocol

#### Media and growth conditions

The wild-type and deletion mutant strains were grown in YPD (1 % yeast extract, 2 % peptone, 2 % dextrose) at 30° C. The conditional mutant strains were grown in YPG (1 % yeast extract, 2 % peptone, 2 % galactose) as a permissive medium and YPD as a non-permissive medium at 30° C. The strains and primers used in this study are listed in Table S1 in the Supporting Material.

#### Construction of mCherry Sad1 overexpression strain in DYN1-3XGFP background

Dynein was tagged with triple GFP under the native promoter in H99 background as mentioned previously [16] to generate CNNV121. For the N-terminal tagging of Sad1 protein, the cassette was amplified by PCR using terminal primers (VYP202 and VYP203) from the strain harboring SAD1 gene placed under GAL7-mCherry sequence (CNVY182). The resulting PCR product (approximately 6.4 kb) was transformed into the strain where DYN1 is tagged with 3X GFP (CNNV121) using biolistic method [84].

#### Construction of conditional mutant of DYN1 in bim1Δ GFP-H4 and GFP-H4 backgrounds

The conditional mutants of DYN1 were constructed by amplifying the cassette from plasmid pUSG7DYN1 [16] using terminal primers (NV404 and NV356). The resulting PCR product (approximately 5.3 kb) was transformed into H99 and bim1Δ strains expressing GFP-H4 (CNVY108 and CNNV107 respectively) using biolistic method [84]. The transformants obtained were confirmed using primers (VYP131 and NV405).

#### Microscopic image acquisition, live-cell imaging and analysis

The wild-type and deletion mutant strains were grown in YPD overnight. The cells were washed with 1x phosphate-buffered saline (PBS) and the cell suspension was placed on agarose (2 %) patch present on the slide which was covered by a coverslip. The images were acquired at room temperature using laser scanning inverted confocal microscope LSM 880-Airyscan (ZEISS, Plan Apochromat 63x, NA oil 1.4) equipped with highly sensitive photo-detectors or Axio Observer Calibri (ZEISS). The Z-stack images were taken at every *µ*m. Live-cell imaging was performed at 30°C on an inverted confocal microscope (ZEISS, LSM-880) equipped with a temperature-control chamber (Pecon incubator, XL multi SL), a Plan Apochromat 100x NA oil 1.4 objective and GaAsp photodetectors. Two-color images were taken by the sequential switching between RFP and GFP filter lines (GFP/FITC 488, mCherry 561 for excitation and GFP/FITC 500/550 band-pass, mCherry 565/650 longpass for emission). For time-lapse microscopy of mCherry-CENP-A and GFP-Tub1, images were collected at a 2-min interval with 1.5 % intensity exposure and a 60-s interval with 1 % intensity exposure, respectively, with 0.5 *µ*m Z-steps. To quantitate the defects in the conditional mutants CNSD155 and CNSD156, the overnight culture grown in permissive condition was reinoculated into permissive and non-permissive media and grown at 30°C, 180 RPM for 12 hrs respectively. The cells were pelleted and washed with 1X PBS (pH 7.4) and were imaged using Axio Observer Calibri (ZEISS) to score for defects. GFP/FITC 488 filter was used for excitation and GFP/FITC 500/550 band pass for emission. Z-stack images were taken at every 0.6 *µ*m. All the images were displayed after the maximum intensity projection of images at each time using ImageJ and processed using ZEISS Zen software/ImageJ/Adobe photoshop.

#### Post-acquisition analysis

The budding indices of cells were calculated as the ratio of the daughter bud’s diameter to the mother bud’s diameter. The diameters were measured by drawing a straight line using a line tool in ImageJ.

## AUTHOR CONTRIBUTIONS

R.P. and K.S. conceived and directed the study; S.C. and R.P. wrote the manuscript with K.S.; S.C. and S.S. performed the theoretical modeling, N.V. and P.S. performed experiments and analyzed data along with other coauthors. All coauthors contributed in editing of the revised manuscript and approved the content.

## ACKNOWLEDGMENTS

This work was supported by the fellowship from SERB (Science and Engineering Research Board), Department of Science and Technology (DST), India (EMR/2017/001346) to RP and Tata Innovation Fellowship (BT/HRD/35/01/03/2017) to KS. KS acknowledges the J.C. Bose National Fellowship from Science and Engineering Research Board (SERB) (JCB/2020/000021), Government of India and a research grant from SERB (CRG/2019/005549). SC was supported by a fellowship from the University Grants Commission (UGC), India, SS was supported by the fellowship from INSPIRE (IF131156) program DST, India, NV was supported by the fellowships (09/733 (0253)/219-EMR-I) and 9/733 (0161)/2011-EMR-I from Council of Scientific & Industrial Research (CSIR), India. PS is a senior research fellow supported by JNCASR, India.

## References

[1] M. Dogterom and S. Leibler, Physical Review Letters 70(9), 1347 (1993).

[2] J. Wu and A. Akhmanova, Annu. Rev. Cell Dev. Biol. 33, 4.1–4.25 (2017).

[3] M. Bornens, Science 335, 422–26 (2012).

[4] P. T. Conduit, A. Wainman, and J. W. Raff, Nat. Rev. Mol. Cell Biol. 16, 611 (2015).

[5] V. Rodionov, E. Nadezhdina, and G. Borisy, PNAS 96, 115–20 (1999).

[6] K. Chabin-Brion, J. Marceiller, F. Perez, C. Settegrana, A. Drechou, G. Durand, and C. Poüs, Mol. Biol. Cell 12, 2047–60 (2001).

[7] L. Hurtado, C. Caballero, M. P. Gavilan, J. Cardenas, M. Bornens, and R. M. Rios, Journal of Cell Biology 193, 917–33 (2011).

[8] T. Vinogradova, R. Paul, A. D. Grimaldi, J. Loncarek, P. M. Miller, D. Yampolsky, V. Magidson, A. Khodjakov, A. Mogilner, and I. Kaverina, Mol. Biol. Cell 23, 820–33 (2012).

[9] T. Vinogradova, P. M. Miller, and I. Kaverina, Cell Cycle 8, 2168 (2009).

[10] J. Paz and J. Lüders, Trends in Cell Biology 28(3), 176 (2018).

[11] J. V. Kilmartin, Philos Trans R Soc Lond B Biol Sci. 369, 20130456 (2014).

[12] T. chen Lin, A. Neuner, and E. Schiebel, Trends Cell Biol. 25, 296–307 (2014).

[13] T. chen Lin, A. Neuner, Y. T. Schlosser, A. N. Scharf, L. Weber, and E. Schiebel, Elife 3, e02208 (2014).

[14] S. Sutradhar, V. Yadav, S. Sridhar, L. Sreekumar, D. Bhattacharyya, S. K. Ghosh, R. Paul, and K. Sanyal, Molecular Biology of the Cell 26, 3954 (2015).

[15] L. Kozubowski, V. Yadav, G. Chatterjee, S. Sridhar, M. Yamaguchi, S. Kawamoto, I. Bose, J. Heitman, and K. Sanyal, mBio 4(5), 00614 (2013).

[16] N. Varshney, S. Som, S. Chatterjee, S. Sridhar, D. Bhattacharyya, R. Paul, and K. Sanyal, PLoS Genet 15(2), e1007959 (2019).

[17] J. Heitman, Fungal Biol Rev. 25(1), 48 (2011).

[18] G. Janbon, K. Ormerod, D. Paulet, E. B. 3rd, V. Yadav, G. Chatterjee, N. Mullapudi, C. Hon, R. Billmyre, F. Brunel, Y. Bahn, W. Chen, Y. Chen, E. Chow, J. Coppee, A. Floyd-Averette, C. Gaillardin, K. Gerik, J. Goldberg, S. Gonzalez-Hilarion, S. Gujja, J. Hamlin, Y. Hsueh, G. Ianiri, S. Jones, C. Kodira, L. Kozubowski, W. Lam, M. Marra, L. Mesner, P. Mieczkowski, F. Moyrand, K. Nielsen, C. Proux, T. Rossignol, J. Schein, S. Sun, C. Wollschlaeger, I. Wood, Q. Zeng, C. Neuveglise, C. Newlon, J. Perfect, J. Lodge, A. Idnurm, J. Stajich, J. Kronstad, K. Sanyal, J. Heitman, J. Fraser, C. Cuomo, and F. Dietrich., PLoS Genet. 10, e1004261 (2014).

[19] D. Srikanta, F. Santiago-Tirado, and T. Doering, Yeast 31(2), 47 (2014).

[20] V. Yadav and K. Sanyal, mSphere 3(4), e00190 (2018).

[21] N. Varshney and K. Sanyal, Current Genetics 65, 1341 (2019).

[22] C. G. Rasmussen, J. A. Humphries, and L. G. Smith, Annual Review of Plant Biology 62, 387 (2011).

[23] B. Byers and L. Goetsch, J Bacteriol. 124(1), 511–23 (1975).

[24] M. Winey and E. O’Toole, Nat Cell Biol. 3(1), E23–7 (2001).

[25] C. D. Souza and S. Osmani, Eukaryot Cell 6(9), 1521–27 (2007).

[26] M. Yamaguchi, S. K. Biswas, Y. Kuwabara, M. Ohkusu, M. Shimizu, and K. Takeo, Journal of Electron Microscopy 59(2), 165 (2010).

[27] M. Yamaguchi, M. Ohkusu, S. Biswas, and S. Kawamoto, Nihon Ishinkin Gakkai Zasshi. 48(4), 147–52 (2007).

[28] T. Tanaka, M. Stark, and K. Tanaka, Nat Rev Mol Cell Biol 6(12), 929–42 (2005).

[29] J. Thakur and K. Sanyal, Eukaryot Cell. 10, 1295 (2011).

[30] J. Thakur and K. Sanyal, PLoS Genet. 8, e1002661 (2012).

[31] A. Gladfelter and J. Berman, Int Rev Cytol. 7(12), 875 (2009).

[32] I. Heath, Int Rev Cytol. 64, 1–80 (1980).

[33] A. Straube, M. Brill, B. Oakley, T. Horio, and G. Steinberg, Mol Biol Cell 14, 642 (2003).

[34] M. Piel and P. T. Tran, Current Biology 19 (17), R823 (2009).

[35] K. E. Sawin, P. C. Lourenco, and H. A. Snaith, Current Biology 14, 763 (2004).

[36] S. Chatterjee, A. Sarkar, J. Zhu, A. Khodjakov, A. Mogilner, and R. Paul, Biophysical Journal 119(2), 434 (2020).

[37] T. Kiyomitsu and I. M. Cheeseman, Nat. Cell Biol. 14, 311 (2012).

[38] D. H. Park and L. S. Rose, Dev. Biol. 315, 42 (2008).

[39] F. J. McNally, J. Cell Biol. 200(2), 131 (2013).

[40] M. Théry, A. Jiménez-Dalmaroni, V. Racine, M. Bornens, and F. Julicher, Nature 447(7143), 493 (2007).

[41] N. Minc, D. Burgess, and F. Chang, Cell 144, 414 (2011).

[42] G. Gay, T. Courtheoux, C. Reyes, S. Tournier, and Y. Gachet, J Cell Biol 196, 757 (2012).

[43] N. Pavin and I. M. Tolic, Annu Rev Biophys 45, 279 (2016).

[44] A. V. Zaytsev, D. Segura-Pena, M. Godzi, A. Calderon, E. R. Ballister, R. Stamatov, A. M. Mayo, L. Peterson, B. E. Black, F. I. Ataullakhanov, M. A. Lampson, and E. L. Grishchuk, Elife 5, e10644 (2016).

[45] M. W. Elting, M. Prakash, D. B. Udy, and S. Dumont, Curr Biol 27, 2112 (2017).

[46] J. Li, L. Cheng, and H. Jiang, Mol Biol Cell 30, 2458 (2019).

[47] G. Letort, I. Bennabia, S. Dmitrieffb, F. Nedelec, M.-H. Verlhaca, and M.-E. Terreta, Mol Biol Cell. 30(7), 863 (2019).

[48] A. R. Lamson, C. J. Edelmaier, M. A. Glaser, and M. D. Betterton, Biophysical Journal 116, 1719 (2019).

[49] N. P. Ferenz, R. Paul, C. Fagerstrom, A. Mogilner, and P. Wadsworth, Curr Biol. 19(21), 1833–1838 (2009).

[50] F. Nedelec, J Cell Biol 158, 1005 (2002).

[51] R. Wollman, G. Civelekoglu-Scholey, J. M. Scholey, and A. Mogilner, Mol Syst Biol 4, 195 (2008).

[52] B. Rubinstein, K. Larripa, P. Sommi, and A. Mogilner, Phys Biol 6, 016005 (2009).

[53] I. Kalinina, A. Nandi, P. Delivani, M. R. Chacón, A. H. Klemm, D. Ramunno-Johnson, Krull, B. Lindner, N. Pavin, and I. M. Tolić-Nørrelykke, Nat Cell Biol. 15(1), 82 (2013).

[54] H. Maiato, A. M. Gomes, F. Sousa, and M. Barisic, Biology (Basel) 6 (2017), 10.3390/biology6010013.

[55] S. Som, S. Chatterjee, and R. Paul, Phys. Rev. E 99, 012409 (2019).

[56] W. Korinek, M. Copeland, A. Chaudhuri, and J. Chant., Science 287, 2257 (2000).

[57] L. Lee, J. Tirnauer, J. Li, S. Schuyler, J. Liu, and D. Pellman, Science 287, 2260 (2000).

[58] R. Miller, S. Cheng, and M. Rose, Mol. Biol. Cell. 11, 2949 (2000).

[59] C. Cepeda-García, N. Delgehyr, M. A. J. Ortiz, R. ten Hoopen, A. Zhiteneva, and M. Segal, Molecular Biology of the Cell 21(15), 2685–2695 (2010).

[60] K. Shulist, E. Yen, S. Kaitna, A. Leary, A. Decterov, D. Gupta, and J. Vogel, Scientific Reports 7, 11398 (2017).

[61] S. C. Lee and J. Heitman, Eukaryot Cell. 11(6), 783–94 (2012).

[62] M. W. Staudt, E. K. Kruzel, K. Shimizu, and C. M. Hull, Fungal Genet Biol. 47(4), 310 (2010).

[63] T. E. Holy and S. Leibler, PNAS 91 (12), 5682 (1994).

[64] T. Mitchison and M. Kirschner, Nature 312, 237 (1984).

[65] M. Kirschner and T. Mitchison, Cell 45, 329 (1986).

[66] L. Winters, I. Ban, M. Prelogovic, I. Kalinina, N. Pavin, and I. M. Tolic, BMC Biology, 10.1186/s12915 (2019).

[67] S. Baumgärtner and I.M. Tolić, PLoS ONE 9(4), e93781 (2014).

[68] G. Civelekoglu-Scholey, D.J. Sharp, A. Mogilner, and J. Scholey, Biophysical Journal 19(11), 3966 (2006).

[69] R. Farhadifar, C. F. Baer, A. C. Valfort, E. C. Andersen, T. Muller-Reichert, M. Delattre, and D. J. Needleman, Curr Biol 25, 732 (2015).

[70] M. Kwon, S. A. Godinho, N. S. Chandhok, N. J. Ganem, A. Azioune, M. Thery, and D. Pellman, Genes & Development 22, 2189 (2008).

[71] F. Gergely and R. Basto, Genes & Development 22, 2291 (2008).

[72] V. Marthiens, M. Piel, and R. Basto, Journal of Cell Science 125, 3281 (2012).

[73] M. Yoshida, S. Katsuyama, K. Tateho, H. Nakamura, J. Miyoshi, T. Ohba, H. Matsuhara, F. Miki, K. Okazaki, T. Haraguchi, O. Niwa, Y. Hiraoka, and A. Yamamoto, J Cell Biol 200(4), 385 (2013).

[74] S. A. Rincon, A. Lamson, R. Blackwell, V. Syrovatkina, V. Fraisier, A. Paoletti, M. D. Betterton, and P. T. Tran, Nature Communications 8, 15286 (2017).

[75] M. Jagrić, P. Risteski, J. Martinčić, A. Milas, and I.M. Tolić, eLife 10:e61170 (2021).

[76] B. Polak, P. Risteski, S. Lesjak, and I.M. Tolić, EMBO Rep 18, 217 (2017).

[77] Z. Lansky, M. Braun, A. Lüdecke, P. R. ten Wolde, M. E. Janson, and S. Diez, Cell 160(6), 1159 (2015).

[78] I. Gaska, M. E. Armstrong, A. Alfieri, and S. Forth, Develepmental Cell 54(3), 367 (2020).

[79] M. K. Balasubramanian, E. Bi, and M. Glotzer, Current Biology 14, R806–R818 (2004).

[80] L. Laan, N. Pavin, J.H.JG. Romet-Lemonne, M. van Duijn, M. P. Lopez, R. D. Vale, F. Jülicher, S. L. Reck-Peterson, and M. Dogterom, Cell. 148(3), 502 (2012).

[81] N. Pavin, L. Laan, R. Ma, M. Dogterom, and F. Jülicher, New Journal of Physics 14, 105025 (2012).

[82] R. Ma, L. Laan, M. Dogterom, N. Pavin, and F. Jülicher, New Journal of Physics 16, 013018 (2014).

[83] S. W. Grill, K. Kruse, and F. Jülicher, Phys. Rev. Lett. 94, 108104 (2005).

[84] R. Davidson, M. Cruz, R. Sia, B. Allen, J. Alspaugh, and J. Heitman, Fungal Genet Biol. 29(1), 38 (2000).

