## Supporting Material for "Mechanics of MTOC clustering and spindle positioning in budding yeast *Cryptococcus neoformans*"

### Quantification of the experimental data on nuclear migration

The binary classification of nuclear migration timescales is correlated with the budding indices of individual cells. Hence, the data representation in terms of the budding indices is more informative than the simplified binary classification. Some experimental data sets for the budding indices of the cells with the nucleus in the mother bud is already reported in our earlier study (1). In the present analysis, we have now significantly increased our data set to quantify budding indices in large budded cells with an unsegregated nuclear mass from the wild-type, Bim1 deletion and/or Dynein depleted conditions. The quantification of budding indices (B.I) was done in triplicates, the cells with B.I (> 0.5) were analyzed and the data is represented in Fig. S4. However, for the sake of simplicity (and enhanced readability), taking cue from these budding indices-based data sets, we classified cells in various cell cycle stages (i.e various budding index values) into two categories: (a) cells in which the nuclei are retained in the mother bud and (b) cells in which the nuclei are migrated to the daughter bud. The simplified data representation in terms of this binary classification helps to summarize the overall fate of nuclear migration in (a) wild-type cells, (b) Dyn1 depleted cells, (c) Bim1 deleted cells.

### Simulation Procedure

#### Modeling the cellular confinement

The geometric construction of the whole cell is considered to be the intersection of two unequal spheres representing the mother bud (radius $R_M$) and the daughter bud (radius $r_D$). The septin ring is marked by the circle spanned at the intersection of the two spheres. The mother bud volume remains constant while the daughter bud's growth rate is tuned, taking a cue from the experimental observations. The nucleus is taken as a sphere (radius $r_{nuc}$). Underneath the cell boundary, a cortical layer of finite width $l_c$ accounts for the MT interaction at the cortex. The characteristic parameters (e.g. dynein density etc.) of the cortical layer at the mother bud and the daughter bud are controlled independently due to possible functional differences at the molecular level (1, 2).

#### Modeling the MTs

MTs are modelled as semiflexible polymers characterized by four innate dynamic instability parameters, viz, growth velocity ($v_g$), shrinkage velocity ($v_s$), catastrophe frequency ($f_c$) and rescue frequency ($f_r$). While probing for an alternative mechanism, $f_c$ of the MTs searching for the KT is modulated by the local concentration/gradient of the RanGTP like chemical. MTs hitting the cortex experience dynein pull, undergo buckling transition when pivoted and apply an instantaneous pushing force on the nucleus. A subpopulation of MTs, graze along the cortex and the NE.

#### Modeling MTOCs and kinetochores

The KTs and MTOCs are modeled similarly as in the previous studies (1, 2). MTOCs are rigid spheres on the outer NE and the movement of the MTOCs is strictly confined to the NE. The SPBs are considered as rigid spheres (radius $r_{SPB}$) embedded on the NE.

Spherical KTs with a hardcore repulsion ($\vec{f}_{intersection}$) prevent any finite overlap amongst the KTs. The repulsive force is proportional to the instantaneous overlap between two overlapping KTs (1). Sister KTs remain paired with each other by the cohesin spring until metaphase. A Hookean force, proportional to the inter-KT separation, is introduced amongst the sister KTs to account for the effect of the cohesin spring holding them together (1, 2). In the computational model, the Hookean force is given by $\vec{f}_{cohesin} = K_{cohesin}\vec{d}_{KT-KT}$, where $K_{cohesin}$ is the spring constant and $|\vec{d}_{KT-KT}|$ is the instantaneous separation between two sister KTs. Relevant model parameters are listed in Table S2. The description of various MT mediated force interactions is discussed in the following sections and summarized in Table S5.

#### Modeling cMT-cell cortex interaction and the inter cMT coupling at the NE

From the MTOCs and later from the SPBs, the cMTs nucleate in any random direction within the cell, excluding the nucleus. Various force transductions due to cMT-cell cortex interaction in the *in silico* model construction are categorically listed in the following.

1. When a growing cMT reaches to the cortex and hits the cell boundary, the steric cell boundary impedes the MT tip against its growth. We characterize the instantaneous force exerted on the MT tip upon each encounter with the cell boundary as an instantaneous push ($\vec{f}^{cell-mem}_{push-inst}$). The magnitude of the instantaneous push is taken to be 1 pN per MT on each encounter. The net instantaneous push is $\vec{F}^{cell-mem}_{push-inst}= \sum_{cMTs} \vec{f}^{cell-mem}_{push-inst}$.
2. Apart from the instantaneous push, a MT growing inside the cell cortex undergoes sliding. The 'bent' sliding segment inside the cell cortex provides anchorage points for the cortical dyneins. For computational simplicity, we introduce the parameter, mean number of dyneins per unit length in mother (daughter) cortex ($\lambda^{M(D)}_{dyn}$) on the MT segments inside the cortex. The force due to the dynein pull on a cMT ($\vec{f}^{cor}_{dyn}$) is proportional to $\lambda^{M(D)}_{dyn} l_{cor}$, where $l_{cor}$ is the combined length of the penetrated MT segment inside the cell cortex. $M$, $D$ in the suffix denotes mother and daughter bud cell cortex respectively. The net cortical dynein pull is $\vec{F}^{cor}_{dyn}=\sum_{cMTs} \vec{f}^{cor}_{dyn}=\sum_{cMTs} \lambda^{M(D)}_{dyn} l_{cor}\ \vec{f}_{dyn}$, where $\vec{f}_{dyn}$ is the force generated by a single dynein motor.
3. Similarly, the force due to Bim1 bias on a cMT within mother cortex is $\vec{F}^{cor}_{Bim1}=\sum_{cMTs} \vec{f}^{cor}_{Bim1}=\sum_{cMTs} \lambda^{M}_{Bim1} l_{cor}\ \vec{f}_{Bim1}$ where $\vec{f}_{Bim1}$ is the force generated by a single Bim1. The force due to Bim1 is directed toward the septin ring (Table S5). In the daughter cortex, there is no Bim1 bias. For simplicity, we chose $\lambda^{M}_{Bim1}=\lambda^{M}_{dyn}$.
4. The MTs impinged at the cell membrane also undergo buckling transition. MT buckling is a type of deformation of the MT under compressive loading force. Owing to the deformation, the MT imparts a restoring force which depends on the buckling length of the MT (estimated in (3)). We considered first-order Euler buckling in the present context. Whether a MT hitting the cell membrane, deep inside the cortex, will transduce dynein mediated pull or a net push away from the cortex due to Euler buckling or undergo catastrophe is dictated by predefined probabilities $P^{cell-mem}_{buckle}$, $P^{cell-mem}_{dyn-pull}$ and $P^{cell-mem}_{cat}$, where $P^{cell-mem}_{buckle}$ = 1- $P^{cell-mem}_{dyn-pull}$-$P^{cell-mem}_{cat}$. The buckling force ($\vec{f}^{cor}_{buckle}$) is proportional to $l^{-2}$ where $l$ is the concerned cMT length. The net buckling force is $\vec{F}^{cor}_{buckle}= \sum_{cMTs} \vec{f}^{cor}_{buckle}$ ($|\vec{f}^{cor}_{buckle}|=D_{buckle} l^{-2}$, where $D_{buckle}$ refers to the flexural rigidity of the MT ~ 200 pN $\mu$m$^2$) (4). In mother cortex, $P^{cell-mem}_{dyn-pull}$=0.0, $P^{cell-mem}_{buckle}$=0.5, $P^{cell-mem}_{cat}$=0.5; in daughter cortex, $P^{cell-mem}_{dyn-pull}$=0.8, $P^{cell-mem}_{buckle}$=0.1, $P^{cell-mem}_{cat}$=0.1.
5. Moreover, a polymerizing MT at the cortex experiences resistance from the cortex on the MT tip in the vicinity of the cell membrane. The force exerted on the MT tip due to this resistance is chosen to be Hookean spring-like. We estimate the force on a polymerizing MT ($f^{MT-poly}_{cor}$) deep inside the cortex as $k_{cor} l_{cor}$. Here, $k_{cor}$ is the equivalent spring constant characterizing $f^{MT-poly}_{cor}$ that pushes the MT tip away from the cortex. We evaluate the net push on the MTOCs/SPBs due to MT polymerization inside the cortex ($F^{MT-poly}_{cor}$) by summing the force contributions ($f^{MT-poly}_{cor}$) overall growing MTs within the cortical region ($F^{MT-poly}_{cor}=\sum^{cortex}_{MTs} f^{MT-poly}_{cor}$).

The overall load on a cMT at the cortex due to the resistance from the cortical substance modulates the dynamic instability parameters of the cMT. The intrinsic parameters $v_g$ and $f_c$ are tuned as described in the following: (a) $v_g = v_{g0} \exp(f_{load}/f_{stall})$, where $v_{g0}$ represents the unimpeded growth velocity with no load exerted on the cMT, $f_{load}$ and $f_{stall}$ denote load force and stall force per cMT respectively; (b) $f_c$=$f^{stall}_c$/[1+ ($f^{stall}_c/f_{c0}$ -1) exp ($f_{load}/f_{stall}$)]. The load force $f_{load} \sim -k_{cor} l_{cor}$ (1, 2).

To estimate the force due to the inter cMT coupling at the NE $\vec{f}_{overlap}$, we evaluate the net overlap length ($l_{ovl}$) between a pair of overlapping antiparallel MTs. The force exerted on the MTOCs is proportional to $\lambda_{ovl} l_{ovl}$, where $\lambda_{ovl}$ is the mean number of minus end-directed motors per unit length at the MT-MT overlap on the NE.

#### Modeling nMT-kinetochore interaction

The nMTs are categorized into two distinct subsets, interdigitated MTs within the nuclear volume facilitating biorientation of SPBs via molecular motors i.e. interpolar MTs (ipMTs) and MTs establishing contact between SPBs and KTs i.e. kinetochore MTs (kMTs). The collective activity of crossbridging kinesin-5 motors at the ipMT-ipMT overlaps generates a force $f_{ipMT} = l^{ipMT}_{ovl} \lambda_{ipMT} f_{kinesin-5}$ where $l^{ipMT}_{ovl}$ is the total overlapping length summed over all ipMTs emanating from two SPBs. $\lambda_{ipMT}$ is the average number of kinesin-5 motors per unit length at the ipMT-ipMT overlap. $f_{kinesin-5}$ is the force exerted by a single kinesin-5 motor.

Stochastic growth and shrinkage of the kMTs due to dynamic instability leads to a tug of war between several oppositely directed competing forces. During the growth phase of the kMT, the kMT tip imparts a net push $f_{push}^{growth} = l_{pen} K_{fibril}$ on the KT attached to it. Here, $l_{pen}$ is the penetration length of the kMT into the KT, $K_{fibril}$ is the stiffness constant of the connecting spring that mimics the KT fibril in the model. A kMT in shrinkage phase, pulls the KT with a force $f_{pull}^{shrinkage} = l_{separation} K_c$, where $l_{separation}$ is the distance between the kMT tip and the corresponding KT. $K_c$ denotes the stiffness constant of the Hookean spring linking the depolymerizing kMT tip and the KT.

To maintain a constant gap between the SPBs and the KT cluster, a length-dependent catastrophe of the kMTs has been incorporated in the model. The catastrophe frequency of a kMT ($f_c$) is regulated in the following manner, $f_c = h l_{kMT}$, where $l_{kMT}$ is the kMT length (1, 2).

#### Equations of motions governing the spatiotemporal dynamics of the nucleus, MTOCs, kinetochores, SPBs

Due to single cMT interacting with the cortex, forces are applied on the nucleus and the MTOCs/SPBs, simultaneously. Additionally, the force stemming from the ipMT interaction is also exerted on the SPBs. If $\vec{F}_{nucleus}$, $\vec{F}_{MTOC}$ and $\vec{F}_{SPB}$ are the net resultant forces exerted on the nucleus, MTOC and SPB, respectively, the corresponding equations of motion can be gleaned as,

$$\frac{d\vec{R}_{nucleus}}{dt} = \frac{\vec{F}_{nucleus}}{\zeta_{nucleus}} \quad \text{(S1)}$$

$$\frac{d\vec{R}_{MTOC}}{dt} = \frac{\vec{F}_{MTOC}}{\zeta_{MTOC}} \quad \text{(S2)}$$

$$\frac{d\vec{R}_{SPB}}{dt} = \frac{\vec{F}_{SPB}}{\zeta_{SPB}} \quad \text{(S3)}$$

where $\vec{R}_{nucleus}$, $\vec{R}_{MTOC}$ and $\vec{R}_{SPB}$ denote the instantaneous positions of the nucleus, MTOCs and SPBs respectively, at a certain time step. $\zeta_{nucleus}$, $\zeta_{MTOC}$, $\zeta_{SPB}$ represent viscous drag on the corresponding objects. $\vec{F}_{nucleus}$, $\vec{F}_{MTOC}$ and $\vec{F}_{SPB}$ contain vectorial contributions from $\vec{F}_{cor}^{MT-poly}$, $\vec{F}_{push-inst}^{cell-mem}$, $\vec{F}_{dyn}^{cor}$ and $\vec{F}_{buckle}^{cor}$. The net force on an MTOC $\vec{F}_{MTOC}$ is divided into two components: (a) tangential to the NE and (b) normal to the NE. As the movement of MTOC is considered constrained on NE, only the tangential force component is responsible for moving the MTOC on NE, the normal component is not allowed to contribute to the motion of MTOCs confined at the NE. The same procedure applies to the motion of the SPBs as well, as the SPBs are taken to be embedded on the NE throughout the mitotic period considered in the simulation. Due to the attachment of the MTOCs with the surface of the NE, the resultant force on the nucleus can be computed as the vectorial sum over the net cortical forces on individual MTOCs.

In a similar fashion, the motion of a KT is dictated by the following equation of motion,

$$\frac{d\vec{R}_{kinetochore}}{dt} = \frac{\vec{F}_{kinetochore}}{\zeta_{kinetochore}} \quad \text{(S4)}$$

Here, $\vec{R}_{kinetochore}$, $\vec{F}_{kinetochore}$ and $\vec{\zeta}_{kinetochore}$ represent instantaneous position, net force on the KT and the viscous drag experienced by the KT respectively. $\vec{F}_{kinetochore}$ entails a vectorial summation over $\vec{f}_{push}^{growth}$, $\vec{f}_{pull}^{shrinkage}$, $\vec{f}_{ipMT}$, $\vec{f}_{cohesin}$ and $\vec{f}_{intersection}$. For simplicity, we have not considered contributions of thermal diffusion (Brownian motion) while computing the positional update of any of the objects considered in the model.

The equations of motion are discretized and solved using Euler's method at every time step.

### Mathematical model for spindle positioning in budding yeast

In 2-dimensions, we consider that the mother and the daughter bud constitute two intersecting circles of radii $R_M$ and $r_D$ respectively. The axis of symmetry of the cellular confinement (e.g. the mother-daughter conglomerate) is taken to be the X-axis with the origin '$O$' at the center of the mother bud. The center of the daughter bud is placed at a distance '$d$' from the center of the mother (Fig. 2B-2C). We compute various MT-based forces on the spindle upon MT interaction with the cell cortex. The mathematical formulation of different force balance terms (e.g., forces originating from MT pushing, dynein mediated pull on the MTs at the cell cortex, MT buckling, etc.) is characterized by exponential length distribution of MTs (4–6). The direction of the forces is depicted in Fig. 2C.

#### Pushing forces

MTs are nucleated from the two SPBs separated by a distance of $2a$ (spindle length) with spindle-center at $x$. For convenience, we mark the SPB located at the left of the spindle ($SPB(L)$) and the SPB located at the right of the spindle ($SPB(R)$). When an MT nucleates out of the $SPB(L)$, it grows a distance $R_M + x - a$ to establish contact with the cell membrane (Fig. 2B). Hence, the instantaneous pushing force on the $SPB(L)$ due to the MTs hitting the mother bud cell membrane reads,

$$F_{push-inst}^{cell-mem(M)} = A_M e^{-(R_M+x-a)/L_{MT}^{av}} \tag{S5}$$

The exponential weight factor enters the expression due to the length of dynamic MTs governed by the following distribution: $N(l) \propto e^{-l/L_{MT}^{av}}$ where $l$ is the length of the MT under consideration. The instantaneous pushing force from the MTs nucleating out of the $SPB(R)$ has to elongate up to a distance $d + r_D - x - a$ along the axis of symmetry to make contact with the cell membrane in the daughter bud. Hence, the instantaneous pushing force contribution from the daughter bud cell membrane is,

$$F_{push-inst}^{cell-mem(D)} = -A_D e^{-(d+r_D-x-a)/L_{MT}^{av}} \tag{S6}$$

Here $A_M$ and $A_D$ define the amplitude of the instantaneous pushing force contribution from the mother and the daughter bud cell membrane. The prefactors $A_M$ and $A_D$ are proportional to the number of MTs interacting with the cell membrane where instantaneous pushing force per MT is taken to be $\sim$ 1 pN.

#### Pulling forces

When an MT emanating from the $SPB(L)$ grows all the way to penetrate the mother cortex, the dyneins localized at the cortex anchor to the MT segment orchestrating a net pull ($F_{dyn}^{cor(M)}$) toward the mother cortex. Since an elongating MT segment inside the cortex can bend and undergoes sliding as illustrated in the schematic diagram, the pulling force is categorized into two parts: 1. the pull on the uncurled/straight MT segment in the cortex and 2. the pull on the MT segment engaged in 'lateral sliding' along the cortex. Similarly, MTs nucleating out of the $SPB(R)$ upon elongating up to the daughter cortex can penetrate and experience dynein mediated pull. For an MT nucleating from the $SPB(L)$ within the one-dimensional confinement, it has to extend at least up to a distance of $R_M - l_c + x - a$ to establish physical contact with the mother cortex. It is also evident from the schematic diagram that after reaching the cortex, the 'uncurled' MT tip can advance up to $R_M + x - a$ without bending (Fig. 2B). Henceforth the net pull on the $SPB(L)$ from the 'uncurled' MT segments in the mother cortex can be evaluated in the following manner.

$$\begin{aligned} F_{dyn(uncurled)}^{cor(M)} &= -B_M \lambda_{dyn}^M \int_{R_M-l_c+x-a}^{R_M+x-a} e^{-s/L_{MT}^{av}} ds \\ &= -B_M \lambda_{dyn}^M L_{MT}^{av} (e^{l_c/L_{MT}^{av}} - 1) e^{-(R_M+x-a)/L_{MT}^{av}} \end{aligned} \tag{S7}$$

Similarly, we can also evaluate the net dynein pull contribution from the 'arc' like MT segment sliding within the mother cortex. Therefore, the net force due to the dynein pull on the sliding 'arc' $F_{dyn(sliding)}^{cor(M)}$ reads

$$\begin{aligned} F_{dyn(sliding)}^{cor(M)} &= -B_M \lambda_{dyn}^M \int_0^{l_{arc}^M} ds\ e^{-(R_M+x-a+s)/L_{MT}^{av}} \\ &= -B_M \lambda_{dyn}^M L_{MT}^{av} e^{-(R_M+x-a)/L_{MT}^{av}} (1 - e^{-l_{arc}^M/L_{MT}^{av}}) \end{aligned} \tag{S8}$$

Here $l_{arc}^M$ is the 'arc' like segment traced from the intersection of the axis of symmetry and the mother bud cell membrane to the peripheral contact of the septin ring with the mother bud cell membrane. Here for the sake of simplicity, we assume that the sliding MTs passing through the mother cortex extend up to the septin ring. The value of $l_{arc}^M$ is estimated to be

$$l_{arc}^M = \frac{\pi R_M}{2} + R_M \tan^{-1}(x_{sp}/y_{sp}) \tag{S9}$$

From $SPB(R)$ we assume that the MTs nucleate in the direction of the daughter bud and interact with the daughter cortex and orchestrate a pull ($F_{dyn}^{cor(D)}$) in a similar fashion with which MTs from $SPB(L)$ interact with the mother cortex. Thus, the net pull due to the anchored dyneins on the 'uncurled' MT segments inside the daughter cortex reads,

$$\begin{aligned} F_{dyn(uncurled)}^{cor(D)} &= B_D \lambda_{dyn}^D \int_{d+r_D-l_c-(x+a)}^{d+r_D-(x+a)} ds\ e^{-s/L_{MT}^{av}} \\ &= B_D \lambda_{dyn}^D L_{MT}^{av} (e^{l_c/L_{MT}^{av}} - 1) e^{-(d-x-a+r_D)/L_{MT}^{av}} \end{aligned} \tag{S10}$$

Similarly, we can compute the contribution in the net dynein pull on $SPB(R)$ from the 'sliding arc' passing through the daughter cortex in the following manner.

$$
\begin{aligned}
F^{cor(D)}_{dyn(sliding)} &= B_D \lambda^D_{dyn} \int_0^{l^D_{arc}} ds\, e^{-(d+r_D-x-a+s)/L^{av}_{MT}} \\
&= B_D \lambda^D_{dyn} L^{av}_{MT} e^{-(d+r_D-x-a)/L^{av}_{MT}} (1 - e^{-l^D_{arc}/L^{av}_{MT}})
\end{aligned}
\tag{S11}
$$

Here, $l^D_{arc}$ is estimated to be the arc length traced from the intersection of the axis of symmetry with the daughter cortex and the peripheral contact of the septin ring with the cell membrane. From the schematic diagram we obtain the $l^D_{arc}$ to be

$$
l^D_{arc} = \frac{\pi r_D}{2} + r_D \tan^{-1}\left(\frac{d - x_{sp}}{y_{sp}}\right) \tag{S12}
$$

Note here that the prefactors $B_M$ and $B_D$ are proportional to the number of MTs experiencing dynein mediated pulling in the mother and the daughter cortex, respectively.

We have assumed that the effective dynein density is smeared uniformly across the whole mother and daughter cortex in the above analysis. However, experimental observations point toward a non-uniform, differential spatial arrangement of dyneins in the mother and the daughter cortex. These observations prompted us to accommodate the differential spatial profiling of dynein in the mother and the daughter cortex into the current mathematical model. Force transduction due to the differential spatial organization of dyneins in the mother and daughter cortex is described in detail in the following section.

#### Buckling forces

MTs with tips passed into the mother(daughter) cortex can buckle where the buckling probability is a tunable parameter in the current model. MTs impinged and pivoting at the mother(daughter) cortex generate a length-dependent 'pushing' force that increases with the squared inverse length of the MT while undergoing buckling. It is evident from the geometric construction that the forces stemming from MT buckling at the mother and daughter cortex are oppositely directed (Fig. 2C). We have taken the following form for the force produced by cortical buckling at the mother cortex ($F^{cor(M)}_{buckle}$)

$$
F^{cor(M)}_{buckle} = \frac{D_M}{(R_M + x - a)^2}\, e^{-(R_M+x-a)/L^{av}_{MT}} \tag{S13}
$$

Similarly, force generated due to the buckling at the daughter cortex $F^{cor(D)}_{buckle}$ is taken as

$$
F^{cor(D)}_{buckle} = -\frac{D_D}{(d + r_D - x - a)^2}\, e^{-(d+r_D-x-a)/L^{av}_{MT}} \tag{S14}
$$

$D_M$ and $D_D$ are proportional to the buckling amplitude times the average number of MTs undergoing buckling transition in the mother and daughter cortices, respectively.

### One-dimensional mathematical model describing the forces due to localized dynein organization in the daughter bud cell cortex:

In the mathematical template, we assume that the mother cortex's dynein density is uniform, as considered in the earlier scenario. However, in the daughter cortex, the dyneins are spatially localized as discrete patches of length $l_1$ with the gap between two consecutive patches being $l_2$. We further assume that the effective dynein density within a patch falls exponentially with distance. Hence,

$$
\lambda^D_{dyn}(s) = \lambda^D_{dyn}(0) e^{-s/\xi^D_{dyn}} \tag{S15}
$$

Here, $\lambda^D_{dyn}(0)$ denotes the effective dynein density at the intersection of the axis of symmetry and the daughter cortex. $\xi^D_{dyn}$ is an external tunable parameter in the current model which dictates how the effective dynein density drops off within the patch as one moves away from the intersection of the axis of symmetry and the daughter cortex in the transverse direction along the daughter cortex. It is evident from the present model construction that the net contribution to $F^{cor(D)}_{dyn(sliding)}$ will come from the discrete patches only. Hence the integration $\int_0^{l^D_{arc}}$ will have nonzero finite contributions from $\int_0^{l_1} + \int_{l_1+l_2}^{l_2+2l_1} + \ldots$ only. Now the expression for $F^{cor(D)}_{dyn(sliding)}$ reads,

$$\begin{aligned}
F^{cor(D)}_{dyn(sliding)} &= B_D \int_0^{l^D_{arc}} ds\ e^{-(d+r_D-x-a+s)/L^{av}_{MT}} \lambda^D_{dyn}(s) \\
&= B_D \lambda^D_{dyn}(0) \int_0^{l^D_{arc}} ds\ e^{-(d+r_D-x-a+(1+\gamma)s)/L^{av}_{MT}} \\
&= B_D \lambda^D_{dyn}(0) \Bigg[ \int_0^{l_1} ds\ e^{-(d+r_D-x-a+(1+\gamma)s)/L^{av}_{MT}} + \\
& \int_{l_1+l_2}^{l_2+2l_1} ds\ e^{-(d+r_D-x-a+(1+\gamma)s)/L^{av}_{MT}} + \int_{2l_2+2l_1}^{2l_2+3l_1} ds\ e^{-(d+r_D-x-a+(1+\gamma)s)/L^{av}_{MT}} + ... \Bigg] \\
&= \frac{B_D \lambda^D_{dyn}(0) L^{av}_{MT}}{(1+\gamma)} e^{-(d+r_D-x-a)/L^{av}_{MT}} \Bigg[ 1 - e^{-(1+\gamma)l_1/L^{av}_{MT}} + e^{-(1+\gamma)(l_1+l_2)/L^{av}_{MT}} \\
& - e^{-(1+\gamma)(l_2+2l_1)/L^{av}_{MT}} + e^{-(1+\gamma)(2l_2+2l_1)/L^{av}_{MT}} - e^{-(1+\gamma)(2l_2+3l_1)/L^{av}_{MT}} + ... \Bigg]
\end{aligned} \tag{S16}$$

In the above expressions we take, $L^{av}_{MT}/\xi^D_{dyn} = \gamma \implies 1/\xi^D_{dyn} = \gamma/L^{av}_{MT}$. Let us reconstruct the above expression for $F^{cor(D)}_{dyn(sliding)}$ by taking,

$$e^{-(1+\gamma)l_1/L^{av}_{MT}} = \mathcal{P} \tag{S17}$$

$$e^{-(1+\gamma)l_2/L^{av}_{MT}} = \mathcal{Q} \tag{S18}$$

Plugging $\mathcal{P}$ and $\mathcal{Q}$ into the expression of $F^{cor(D)}_{dyn(sliding)}$ we obtain,

$$\begin{aligned}
F^{cor(D)}_{dyn(sliding)} &= \frac{B_D \lambda^D_{dyn}(0) L^{av}_{MT}}{1+\gamma} e^{-(d+r_D-x-a)/L^{av}_{MT}} \\
& \Big[ 1 - \mathcal{P} + \mathcal{P}\mathcal{Q} - \mathcal{P}^2\mathcal{Q} + \mathcal{P}^2\mathcal{Q}^2 - \mathcal{P}^3\mathcal{Q}^2 + ... \Big] \\
&= \frac{B_D \lambda^D_{dyn}(0) L^{av}_{MT}}{1+\gamma} e^{-(d+r_D-x-a)/L^{av}_{MT}} \left[ 1 - \mathcal{P}(1-\mathcal{Q}) \frac{1-(\mathcal{P}\mathcal{Q})^n}{1-\mathcal{P}\mathcal{Q}} \right]
\end{aligned} \tag{S19}$$

where $nl_1 + nl_2 \simeq nl_1 + (n-1)l_2 \simeq l^D_{arc}$. Furthermore, only retaining terms up to the first order we obtain,

$$F^{cor(D)}_{dyn(sliding)} = \frac{B_D \lambda^D_{dyn}(0) L^{av}_{MT}}{1+\gamma} e^{-(d+r_D-x-a)/L^{av}_{MT}} \left[ 1 - e^{-(1+\gamma)l_1/L^{av}_{MT}} \right] \tag{S20}$$

It is evident from the above equation that by putting $\gamma = 0$ and $l_1 = l^D_{arc}$, we get back the earlier expression with uniform dynein density.

#### Steric Forces

We have considered steric hindrance between the cell membrane and the spindle. The steric clash between the cell membrane and the spindle is devised via the hardcore repulsion between cell membrane in the mother bud and $SPB(L)$, and between the cell membrane in the daughter bud and $SPB(R)$. The steric repulsion in the mother bud $F^{steric(M)}_{cell-mem}$ is taken as

$$F^{steric(M)}_{cell-mem} = C_M e^{-\zeta_M(R_M+x-a)/L^{av}_{MT}} \tag{S21}$$

Similarly, the steric repulsion in the daughter bud $F^{steric(D)}_{cell-mem}$ is

$$F^{steric(D)}_{cell-mem} = -C_D e^{-\zeta_D(d+r_D-x-a)/L^{av}_{MT}} \tag{S22}$$

We take $C_M = C_D$ and $\zeta_M = \zeta_D$ since the repulsive forces stemmed at both the mother and daughter cell-membrane upon steric interaction are identical in nature. From the expression, it is also clear that the magnitude of the force solely depends upon the instantaneous overlap between the objects under consideration. Here $C_M$ and $C_D$ refer to the amplitudes of the forces generated due to the steric clashes between cell membrane (mother or daughter) and $SPB(L \text{ or } R)$. The length scale, more precisely, the short-range limit of the steric forces is governed by $\zeta_M$ and $\zeta_D$. As $\zeta_M$ (or $\zeta_D$) with nonzero positive value is levelled up, $F^{steric(M \text{ or } D)}_{cell-mem}$ drops sharply with increasing distance.

To include the effect of the finite size of the $SPB(L \text{ or } R)$ ($r_{SPB}$), in the steric force expressions $R_M$ is replaced by $R_M \to R_M - r_{SPB}$ and $r_D$ is replaced by $r_D \to r_D - r_{SPB}$ where $r_{SPB}$ is the radius of the SPB.

### ADDITIONAL NUMERICAL ANALYSIS OF MTOC CLUSTERING (AGENT-BASED MODEL)

In several models, it has been shown that cellular processes involving MT mediated 'search and capture' mechanisms are often optimized by the regulation of the number of MTs (7, 8). In a similar 'search and capture' process, as the MT number increases, the capture time gradually decreases to a threshold timescale followed by a saturation (9); the scenario implies that upon a step-wise increase in the number of MTs per MTOC, the clustering time is expected to decrease. We further investigated the sensitivity to varying cMT numbers in a case by case fashion; a. the MT number per MTOC is kept constant upon clustering of two MTOCs, b. adding the MT number upon a fusion of the MTOCs up to a maximum limit to 4 MTs per MTOC and then, c. removing the maximum limit respectively (Fig. S3A). We observe that within the currently explored parameter regime, irrespective of the microscopic mechanisms (a fixed number of MT per MTOC or an increasing number of MTs per MTOC) the clustering of the MTOCs can be achieved within the physiological time scale of  25-30 min. The exact mechanism can only be ascertained by future experiments.

It is demonstrated that MTOCs colocalize with KTs (10, 11) (Fig. 1B-1D). In our earlier *in silico* analysis (1), we assumed that the KTs are connected to MTOCs from the beginning of the simulation. We further assumed that each KT is connected to a single MTOC. In the model, at the onset of the simulation, the objects that cluster to form an SPB are taken to be MTOC-KT pairs as depicted in the cartoons of Fig. 1D, Fig. 2A, Fig. S2A-S2E. In this setting, as the MTOCs cluster, KTs also cluster in unison. Therefore, the MTOC clustering time and KT clustering time remain synonymous when the number of KTs per MTOC is chosen to be unity. Note that the total number of MTOCs is fixed at 14 (as the number of KTs/chromosomes in each cell) at the onset of simulation. Next, we relaxed the assumption of single KT per MTOC and examined what happens to the KT clustering if one or more KTs are attached to the MTOCs at the beginning (Fig. S3B). However, the total number of MTOCs and KTs are kept fixed at the onset of the simulation, as before. Our model predicted that if more than one KT is attached to an MTOC, the KT clustering process is marginally faster than the MTOCs (Fig. S3B). Consider the situation where more than one KT is attached to an MTOC, but the total number of KTs and the total number of MTOCs are equal and fixed at 14. Therefore, few MTOCs would be there which are not attached to any KT. Complete KT clustering requires the clustering of only those MTOCs which are associated with KTs. However, complete MTOC clustering involves the fusion of both the following set of MTOCs: (a) MTOCs attached to KTs, (b) MTOCs not associated with any KT. In that scenario, we expect that time taken for all the KTs to cluster would be less than that of MTOC clustering. Therefore, in the outcomes of the present model construction, the marginal time difference in the clustering times of KTs and MTOCs arises, which is evident from Fig. S3B.

Next, we estimated the time it would take when both the following mechanisms: (a) inter cMT coupling at NE and (b) MT-cell cortex interaction with suppressed dynein and enhanced Bim1 bias are acting concomitantly. Note that, in the currently explored model parameter regime, both mechanisms can render complete MTOC clustering within the physiological time limit of about 25-30 min (Fig. 1B-1F, Fig. 3B-3E, Table S3) (1). Now, if the mechanism of MT-cell cortex interaction (with suppressed dynein and enhanced Bim1 bias) is turned on alongside the mechanism of inter cMT coupling at NE, we expect that the clustering time would further reduce. In the model analysis, we tuned the level of Bim1 bias when both the mechanisms are at play. As expected, increased Bim1 bias alongside inter cMT coupling further reduces the clustering time up to a specific time limit in the currently explored parameter regime (Fig. S3C).

### ADDITIONAL NUMERICAL ANALYSIS OF SPINDLE POSITIONING (ANALYTICAL MODEL)

#### Instantaneous push from the mother and the daughter cortex forces the spindle to localize in mother bud

From the analytical model geometry (Fig. 2B-2C) and Fig. S7A, it is clear that the instantaneous push from the mother and the daughter cortex are oppositely directed. In the expressions for the instantaneous pushing forces $F^{cell-mem(M)}_{push-inst}$ and $F^{cell-mem(D)}_{push-inst}$, the percentage of MTs experiencing instantaneous cortical push is determined by $A_M$ and $A_D$ respectively. We varied this percentage as a tunable parameter of the system with the pre-imposed constraint $A_M = A_D$ for this particular case. Since the daughter bud size is smaller than the mother bud, the balancing act between these two competing forces leads to a stable spindle position inside the mother bud (Fig. S7A). A configuration with equal mother and daughter bud size would lead to a stable spindle position at the center of the septin ring. Importantly, due to the very nature of the force which pushes the

$SPB(L$ and/or $R)$ away from the cell membrane mother and/or daughter, in the resultant force balance landscape, only stable fixed points of the spindle position appear (Fig. S7A).

##### In the sole presence of cortical pull from mother and daughter, spindle collapses onto the cortex

The sole presence of cortical pull from the mother and daughter cortex gives rise to unstable fixed points of the spindle position in the force balance contour for various spindle positions. In a parameter space mapped by $\lambda^{M}_{dyn}$= 4.0 $\mu$m $^{-1}$, $\lambda^{D}_{dyn}$ = 5.0 $\mu$m$^{-1}$, we varied $B_M (= B_D)$ and observed that stable fixed points of the spindle position in the mother and daughter cortex with a slew of unstable fixed points sandwiched in between (Fig. S7B). The unstable fixed points of the spindle close to the septin ring indicate that either the spindle collapses onto the mother cortex or the daughter cortex while slightly displaced from the unstable fixed point of the spindle position depending upon the direction of perturbation.

##### Tug of war between cortical push and pull leads to a collapse of the spindle onto the cortex

We first considered a theoretical mechanistic framework, where force transduction only from the mother cortex or the daughter cortex is allowed. In that construction, we observe a sharp spatial transition of the spindle from the daughter (mother) cortex to mother (daughter) cortex upon step-wise increase in dynein density in mother (daughter) cortex as depicted in Fig. S7C (relevant parameters: $A_M$ = 5 pN, $A_D$ = 5 pN, $B_M$ = 5 pN, $B_D$ = 5 pN, wherever applicable). Next, we carried out a sensitivity analysis in a scenario where the instantaneous cortical push and dynein mediated cortical pull stemming from both the mother and daughter cortex are present (Fig. 5A, Fig. S7D). We varied the dynein density in the mother cortex $\lambda^{M}_{dyn}$ keeping the dynein density in the daughter cortex $\lambda^{D}_{dyn}$ fixed at 8.0 $\mu$m$^{-1}$ and vice versa ( $D_M$=$D_D$=0; other parameters fixed at base values as shown in Table S2; $\lambda^{M}_{dyn}$ fixed at 5.0 $\mu$m$^{-1}$, when $\lambda^{D}_{dyn}$ is varied). The force balance showcases that for lower values of dynein density in the mother cortex, cortical pull from the daughter ($\lambda^{D}_{dyn}$ fixed at 8.0 $\mu$m$^{-1}$) paired with the instantaneous push from the mother bud dominate leading to a spatial collapse of the spindle onto the daughter cortex marked by the stable fixed points of the spindle position in the daughter cortex. As the dynein density in the mother cortex ($\lambda^{M}_{dyn}$) is gradually increased, cortical pull from the mother cortex takes the driver's seat. Subsequently, the dominant cortical pull from the mother cortex initiates a spatial transition of the spindle to the mother cortex via a string of unstable fixed points of the spindle position marked by the oblique segment of the 'Z' contour (Fig. S7D). Similarly, when dynein density in the mother cortex ($\lambda^{M}_{dyn}$) is kept fixed at 5 $\mu$m$^{-1}$ and dynein density in the daughter cortex ($\lambda^{D}_{dyn}$) is varied, we observe a spatial collapse of the spindle onto the daughter cortex at higher values of $\lambda^{D}_{dyn}$ with an intermediate regime having three fixed points of the spindle position (stable fixed points at the mother and the daughter cortex with unstable fixed points wedged in between) as discernible in the inverted 'Z' contour (Fig. 5A). We also obtain fixed point contour of the spindle position having stable fixed points at the mother and daughter cortex with unstable fixed points located in between when average MT length ($L^{av}_{MT}$) is varied (Fig. S7E) keeping other relevant parameters fixed (Table S2 and $\lambda^{M}_{dyn}$ = 5 $\mu$m$^{-1}$, $\lambda^{D}_{dyn}$ = 10 $\mu$m$^{-1}$).

##### Combination of MT buckling with cortical push and pull prevents collapse, maintains stable spindle position near septin ring

Allowing force transduction solely either from the mother cortex or the daughter cortex in the presence of MT buckling, the stable fixed points of the spindle position make a spatial transition toward the mother (daughter) cortex from the daughter (mother) cortex upon a gradual increase in cortical dynein density $\lambda^{M}_{dyn}$ or $\lambda^{D}_{dyn}$ (Fig. S7F). It is imperative to note that unlike previous scenarios, in the presence of MT buckling, the spindle does not collapse onto the cortical region; instead, it localizes $\simeq$ 0.5 $\mu$m away from the cortex (Fig. S7F) in the currently explored parameter regime. It is evident from the expression of the buckling forces $F^{cor(M)}_{buckle}$ and $F^{cor(D)}_{buckle}$ that in the vicinity of the cortex, the force generated due to MT buckling shoots up to a very high value. Since the force due to MT buckling is of 'pushing' nature, it pushes the spindle away from the cortex, thus preventing the spindle from collapsing onto the cortex. In presence of MT buckling transition in both the mother and daughter cortex, the balancing act between the governing forces ($F^{cell-mem(M \text{ and } D)}_{push-inst}$, $F^{cor(M \text{ and } D)}_{dyn}$, $F^{cor(M \text{ and } D)}_{buckle}$), leads to a stable positioning of the spindle within the daughter bud in the vicinity of the septin ring (Fig. 5B-5C). We observed that upon step-wise increment in the dynein density within the daughter cortex ($\lambda^{D}_{dyn}$), the stable fixed point contour of the spindle position tend to shift deep inside the daughter cortex owing to an enhancement in the net force directed toward the daughter cortex as shown in Fig. 5B (relevant parameters: $\lambda^{M}_{dyn}$ = 3.0 $\mu$m$^{-1}$, other parameters as in Table S2). We also carried out sensitivity analysis on the robustness of the stable spindle positioning close to the septin ring within the daughter bud upon variation in the average MT length ($L^{av}_{MT}$). We find that across a reasonably significant range of $L^{av}_{MT}$ values, the location of the stable fixed point for the spindle remains unperturbed within the daughter bud (Fig. 5C, relevant parameters: $\lambda^{M}_{dyn}$ = 3.0 $\mu$m$^{-1}$,

$\lambda^{D}_{dyn}$ = 10.0 $\mu$m$^{-1}$, other parameters as in Table S2).

#### Differential spatial arrangement of dyneins in mother and daughter cortex results in force balance landscape with stable as well as unstable positions of the spindle

Our previous study (1) indicate that the spatial arrangement of dyneins in the mother and daughter cortex is different. The dynein puncta in the mother cortex are uniformly distributed, whereas in the daughter cortex a large localized dynein punctum near the axis of symmetry is observed. This differential spatial arrangement alludes to the fact that for proper nuclear migration (e.g., proper spindle positioning) a directed pull from the daughter cortex on the nucleus is crucial to navigating the nucleus inside the daughter bud. Further, we also observed that irrespective of whether the cortical milieu is enriched with uniform dynein distribution at both the mother and the daughter cortex or the differential distribution mentioned above, the innate mechanistic perspective relies upon the net pulling force generation from the daughter cortex. This leads to a question of how the force balance landscape governing the spindle positioning behaves when one makes a gradual transition from localized puncta in the daughter cortex to a full-fledged uniform distribution. To address that, we assign a characteristic length scale $\xi^{D}_{dyn}$ with the effective localized dynein puncta within the daughter cortex. As $\xi^{D}_{dyn}$ is increased, the effective spatial profile of dyneins within the daughter cortex makes a gradual changeover to a uniform distribution. With this mathematical template at hand, we carried out sensitivity analysis on the force balance landscape governed by various MT mediated forces (e.g., instantaneous push, dynein pull, MT buckling transition) upon the variation in the parameter $\xi^{D}_{dyn}$. In the presence of instantaneous push and dynein pull from the daughter (with no force transduction from the mother cortex and no MT buckling), the gradual increment in $\xi^{D}_{dyn}$ leads to an enhancement in the dynein pull from the daughter invoking a spatial changeover of the spindle from the mother cortex to the daughter cortex (Fig. S7G). For lower values of $\xi^{D}_{dyn}$, instantaneous push dominates the force balance landscape shoving the spindle onto the mother cortex (marked by the stable fixed-point contour of the spindle position in the mother cortex). Furthermore, in the sole presence of dynein pull from both the mother and the daughter cortex, the variation in $\xi^{D}_{dyn}$ yields null force contours (the inverted 'Z' contour) having stable fixed points of the spindle position at the mother and the daughter cortex and a string of unstable fixed points of the spindle location sandwiched in between (Fig. S7H). The gradual increase in $\xi^{D}_{dyn}$ leads to stable fixed points of the spindle position inside the daughter cortex, around $\sim$ 0.5 $\mu$m away from the septin ring when instantaneous push, dynein pull and MT buckling at both the mother and daughter cortex act alongside each other (Fig. S7I) in the currently explored parameter regime. Stable spindle positioning at higher values of $\xi^{D}_{dyn}$ (referring to the uniformity within the spatial organization of dyneins) further corroborates with the 'agent-based' simulation, which includes uniform distribution of dyneins in both mother and daughter cortex as depicted in Fig. S5A, Fig. S5B, Fig. S5D and Fig. 5E (third bar), Fig. 5F. The spindle localization is observed to be at a distance $\sim$ 1 $\mu$m from the septin ring inside the daughter bud (Fig. S5A, Fig. S5B, Fig. S5D, Fig. 5E, third bar, and Fig. 5F).

#### Agent-based simulations with differential spatial arrangement of dyneins in mother and daughter cell cortex

We have observed that the spatial arrangement of dyneins in mother and daughter bud is different (1). In view of this, we have carried out a few simulations with the daughter cortex containing dynein punctum as opposed to the uniform dynein density. Note that the dynein/Bim1 distribution in the mother cell cortex is kept uniform as before for the sake of simplicity. In the model setup, the daughter cell cortex contains a distinct 'high-density' dynein patch in the region close to the mother-daughter cell axis (Fig. S13A). We termed this 'dynein-rich' region as the 'axial patch' in the text. Besides this axial patch, we further considered a uniform distribution of dyneins 'smeared' everywhere else inside the daughter cell cortex (Fig. S13A). Note that the dynein density at the axial patch is characterized by $\lambda^{D\ (axial\ patch)}_{dyn}$, whereas the dynein density elsewhere in the daughter cortex is characterized by $\lambda^{D\ (rest)}_{dyn}$. With this model setup at hand, we explored the following scenarios (Fig. S13A-S13B). (a) We considered dynein pull only at the axial patch and no dynein pull from elsewhere in the daughter cortex. In this scenario, we observe that the MTOC clustering remains unperturbed. But the nuclear migration is severely impaired. Our simulation outcomes reveal that only in ~26% of total cell population nuclear migration into the daughter bud occurs, and that too is delayed by ~10 min on average (Fig. S13B). In other ~74% of cell population, the nucleus fails to migrate the daughter bud during the total time covered in the simulation run (Fig. S13B). Now, why does the nuclear migration get severely compromised while the MTOC clustering remains unaffected? The reason is, the mobilization of MTOCs on the NE is largely governed by the MT mediated cortical force transduction from the mother cell cortex. The forces stemming from the daughter cortex do not play a significant role in the clustering process. Contrary to that, the migration of the nucleus toward the daughter bud requires a directed pull on the nucleus from the daughter cortex. In this scenario, the cMTs inside the daughter cortex experience no dynein pull except while hitting the axial patch. Therefore, the effective/overall pull from the daughter cortex significantly diminishes, leading to an impaired nuclear migration. (b) Next, we introduced a 'background' uniform dynein distribution in the daughter cell cortex in addition to the axial patch (Fig. S13B, third and fourth set of bars). In the currently explored parameter regime, we found that the ‘wild-type’ time scale of nuclear migration is rescued when the pull from the axial patch is supplemented by relatively ‘small’ pull stemming from uniform dynein distribution elsewhere inside the daughter cortex (Fig. S13B). (c) In the currently explored parameter range, our simulation outcomes further predict that in a limit where the dynein density is uniform throughout the cell cortex (e.g. density at axial patch $\lambda_{dyn}^{D\ (axial\ patch)}$ = density elsewhere $\lambda_{dyn}^{D\ (rest)}$), the experimentally measured wild-type timescales for MTOC clustering and nuclear migration are also attainable (Fig. S13B, first set of bars).

In summary, our in silico analysis identifies two distinct theoretical possibilities when both the MTOC clustering and nuclear migration timescales obtained from the simulations reasonably concur with the experimentally measured timescales (Fig. S13B): (a) ‘high density’ axial patch supplemented with background ‘low density’ uniform dynein distribution and (b) uniform dynein density throughout. However, as the experiments reveal the prevalence of bright dynein spots in the daughter cortex, the possibility (a) is more likely to be a ‘closer-to-experiment’ approximation/interpretation from the modeling point of view. The modeling analysis described above is further complimented by our earlier study reported in (1).

### Effects of stochastic attachment-detachment of dynein, Bim1 on MTOC clustering and nuclear migration

The agent-based computational model described in the preceding sections discounts all stochasticity apart from MT dynamic instability and MT interaction with the cell cortex. While these are important sources of stochasticity, other sources, including motor binding/unbinding, which occur at much shorter time scales, could significantly change the overall picture we gleaned from the earlier model outcomes. In view of that, we extended our earlier agent-based model and incorporated the feature of stochastic MT-motor attachment-detachment in the existing model. Note that we did not simulate the motors’ explicit walking on the MT tracks; instead, within the model, the stochasticity is introduced in the number of dyneins binding to and unbinding from an MT at each time point. The same feature of stochasticity is applied to Bim1 as well. In the model setup, the stochastic binding/unbinding of dynein, Bim1, only concerns the subpopulation of cMTs that interacts with the cell cortex. The cMTs whose plus ends are coursing through the cytoplasm but yet to reach the cell cortex remain unaffected by the feature of stochastic binding-unbinding.

In what follows, we describe how this additional component of stochastic binding/unbinding of dynein, Bim1 is integrated into the protocols of the existing agent based model setup. Note that, in the model, we mainly considered two principal force-generating candidates inside the cell cortex, dynein and Bim1, which influence the mechanics of MTOC clustering and nuclear migration. Moreover, the earlier model outcomes reveal that the MTOC clustering within the experimentally observed wild-type timescale requires ‘enhanced Bim1 + suppressed dynein activity’ in the mother cell cortex. To that note, upon implementing the stochastic motor dynamics, the MTOC clustering simulations are carried out in a parameter regime characterized by ‘enhanced Bim1 + suppressed dynein activity’. Below we present how the number of dyneins (and Bim1) on an MT at the cortex is computed at each time point of the simulation.

Let us consider $\mathcal{N}_{dyn}$ to be the maximum number of dyneins per $\mu$m that can attach to a cMT segment inside the cell cortex (Fig. S8). The number of dyneins per unit length (per $\mu$m) bound to an MT at any time $t$ is taken as $n_{dyn}(t)$ where $n_{dyn}(t) \leq \mathcal{N}_{dyn}$. Furthermore, the rate at which a ‘free’ unattached dynein binds to a cMT is chosen to be $\bar{K}_{dyn}^{on}$ (Fig. S8). Now the question is at time $t$, how many more dyneins can attach per unit length of a cMT segment inside the cell cortex? This number would depend upon the difference between the two following quantities: (a) $\mathcal{N}_{dyn}$, the maximum number of dyneins that can be accommodated per unit length of an cMT segment and (b) $n_{dyn}(t)$, the number of dyneins already attached to the cMT segment. Effectively the rate at which dyneins bind to a cMT ($k_{dyn}^{on}$) at time $t$ can be written as (12, 13),

$$k_{dyn}^{on}(t) = [\mathcal{N}_{dyn} - n_{dyn}(t)]\bar{K}_{dyn}^{on} \tag{S23}$$

Next we introduced the load dependent unbinding of dynein from the cMTs inside the cell cortex. Let $\bar{K}_{dyn}^{u0}$ be the load independent unbinding rate of dyneins from a cMT (Fig. S8). Furthermore, we consider $\vec{F}_{cMT}^{res}(t)$ to be the net ‘pushing’ force experienced by a cMT (Fig. S8). The net pushing force is the resistance applied on a growing cMT inside the cell cortex against the polymerization directed away from the cell cortex. Therefore, $\vec{F}_{cMT}^{res}$ constitutes the following forces: (a) force due to cMT buckling $\vec{f}_{buckle}^{cor}$, (b) instantaneous push $\vec{f}_{push-inst}^{cell-mem}$ and (c) cortical resistance on growing cMTs $\vec{f}_{cor}^{MT-poly}$. The constituting forces are already described in detail in the preceding sections. For simplicity, we assumed this $\vec{F}_{cMT}^{res}(t)$ as the the load force shared by the attached dyneins at time $t$. Given the load $\vec{F}_{cMT}^{res}(t)$, it is required to estimate how many dyneins can detach per unit length of a cMT segment inside the cell cortex. This number would depend on (a) $n_{dyn}(t)$, the number of dyneins already attached to the unit length of a cMT segment and (b) $\vec{F}_{cMT}^{res}(t)$, the net ‘pushing’ force/load experienced by a cMT against its polymerization. The instantaneous rate at which dyneins detach/unbind from a MT ($k_{dyn}^{off}(t)$) can be written as (12–15),

$$k_{dyn}^{off}(t) = n_{dyn}(t)\bar{K}_{dyn}^{u0} \exp\left[\frac{|\vec{F}_{cMT}^{res}|}{F_{dyn}^{detach}}\right] \tag{S24}$$

Here, $F_{dyn}^{detach}$ is the characteristic detachment force of dynein, which is, in our case, a predetermined parameter input to the model system.

Similarly, we can also implement the stochastic binding/unbinding protocol for Bim1 in the mother cell cortex. The effective rate at which Bim1 binds to a cMT ($k_{Bim1}^{on}$) at time $t$ can be written as (12, 13),

$$k_{Bim1}^{on}(t) = [\mathcal{N}_{Bim1} - n_{Bim1}(t)]\bar{K}_{Bim1}^{on} \tag{S25}$$

Here, $\mathcal{N}_{Bim1}$ is the maximum number of Bim1 that can attach per unit length of a cMT segment (Fig. S8) and $n_{Bim1}(t)$ is the number of Bim1 already attached to the cMT segment at time $t$. $\bar{K}_{Bim1}^{on}$ denotes the rate at which an unattached Bim1 attaches to a cMT (Fig. S8).

The load dependent unbinding rate of Bim1 ($k_{Bim1}^{off}(t)$) (12, 13, 15, 16),

$$k_{Bim1}^{off}(t) = n_{Bim1}(t)\bar{K}_{Bim1}^{u0} \exp\left[\frac{|\vec{F}_{cMT}^{res}|}{F_{Bim1}^{detach}}\right] \tag{S26}$$

Here, $\bar{K}_{Bim1}^{u0}$ denotes the load independent unbinding rate of Bim1 from a cMT (Fig. S8) and $F_{Bim1}^{detach}$ is the characteristic detachment force of Bim1. The quantity $F_{Bim1}^{detach}$ is a predetermined parameter input to the model system. The base parameter values considered in the simulation are acquired from previous studies (12, 13, 17–20) and numerical experimentation.

#### Simulations and sensitivity to model parameters

As explained in the previous section, there are four sets of parameters that govern the stochastic attachment-detachment of dynein and Bim1: (a) maximum number of dyneins $\mathcal{N}_{dyn}$ (and Bim1 $\mathcal{N}_{Bim1}$) that can attach to a cMT segment inside the cell cortex per $\mu$m, (b) attachment rates of dynein $\bar{K}_{dyn}^{on}$ and Bim1 $\bar{K}_{Bim1}^{on}$, (c) detachment rates of dynein $\bar{K}_{dyn}^{u0}$ and Bim1 $\bar{K}_{Bim1}^{u0}$, (d) characteristic detachment force of dynein $F_{dyn}^{detach}$ (and Bim1 $F_{Bim1}^{detach}$). We carried out sensitivity analysis across these above mentioned parameters. The outcomes from this *in silico* analysis are reported in Fig. S9-S10. In a nutshell, our simulations identify certain parameter regimes where the time scales of MTOC clustering and nuclear migration remain insensitive to parameter variations and stays within a reasonable range close to the ‘realistic’/‘proper’ time scales.

### SUPPORTING FIGURES

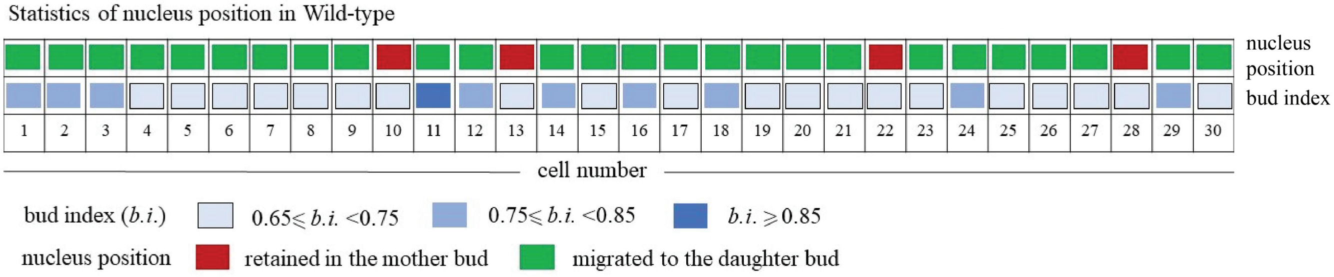


FIGURE S1 Statistics of whether the nucleus is localized in mother bud or daughter bud in wild-type cells. Each column represents a cell indicating the color-coded status of its budding index and the localization of the nucleus within the cell.

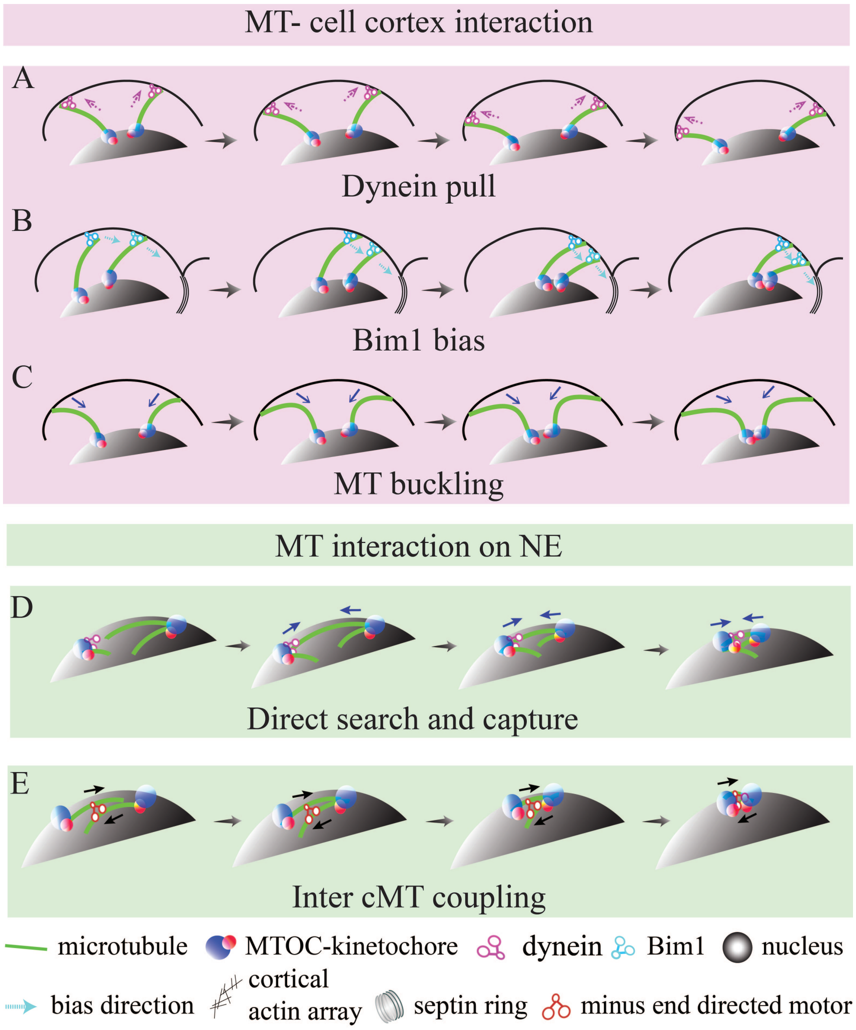


FIGURE S2 Schematic depicting the movement of MTOCs under the effect of various MT mediated forces accounted in the model. (A-C) Representative cartoons illustrate the trajectory of MTOCs due to various forces originating from MT-cell cortex interaction: dynein pull (A), Bim1 bias (B), MT buckling (C). (D-E) Representative cartoons illustrate the trajectory of MTOCs due to various forces originating from MT interaction on NE: MTs directly 'search and capture' MTOCs (D) and inter cMT coupling on NE via minus end-directed motors (E).

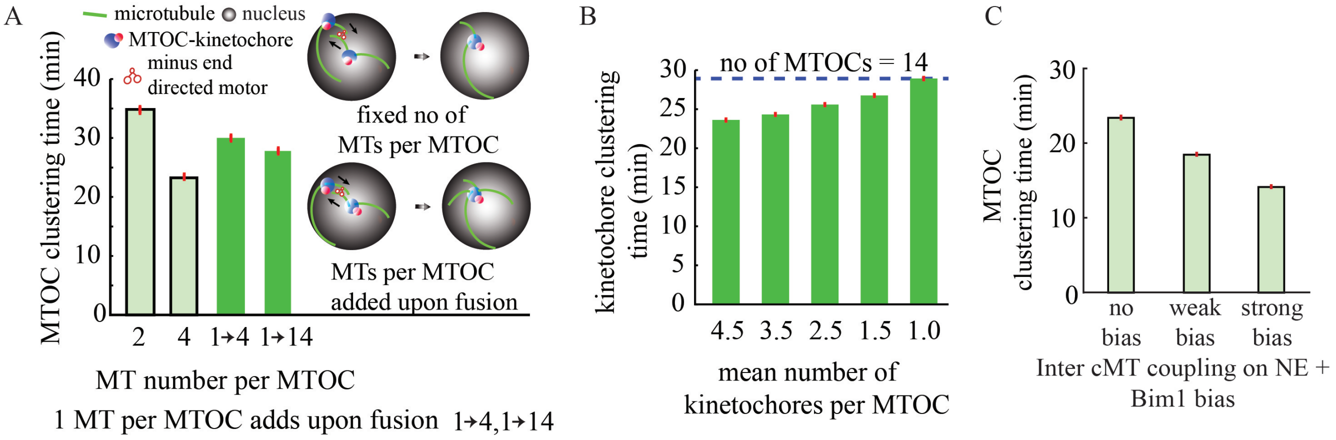


FIGURE S3 Additional numerical tests for MTOC clustering. (A) (First and second bars from left) MTOC clustering time with a fixed number of cMTs per MTOC. Clustering solely via 'inter cMT coupling at the NE' is considered here. (Third bar, 1 ⟶ 4) Clustering time when every MTOC nucleates a single cMT initially and upon the fusion of the MTOCs the MTs aggregate up to a maximum of 4 cMTs per MTOC. (Fourth bar, 1 ⟶ 14) Similar to the "Third bar" but no constraint on the number of cMTs that can add up to the number of MTOCs. (B) Asynchronous assembly of KTs/MTOCs, if more than one KT is associated with a single MTOC. (C) MTOC clustering time when two proposed modes of MTOC clustering namely 1. the inter cMT coupling on NE and 2. MT-cell cortex interaction having enhanced Bim1 bias and diminished cortical pull are considered to be acting together.

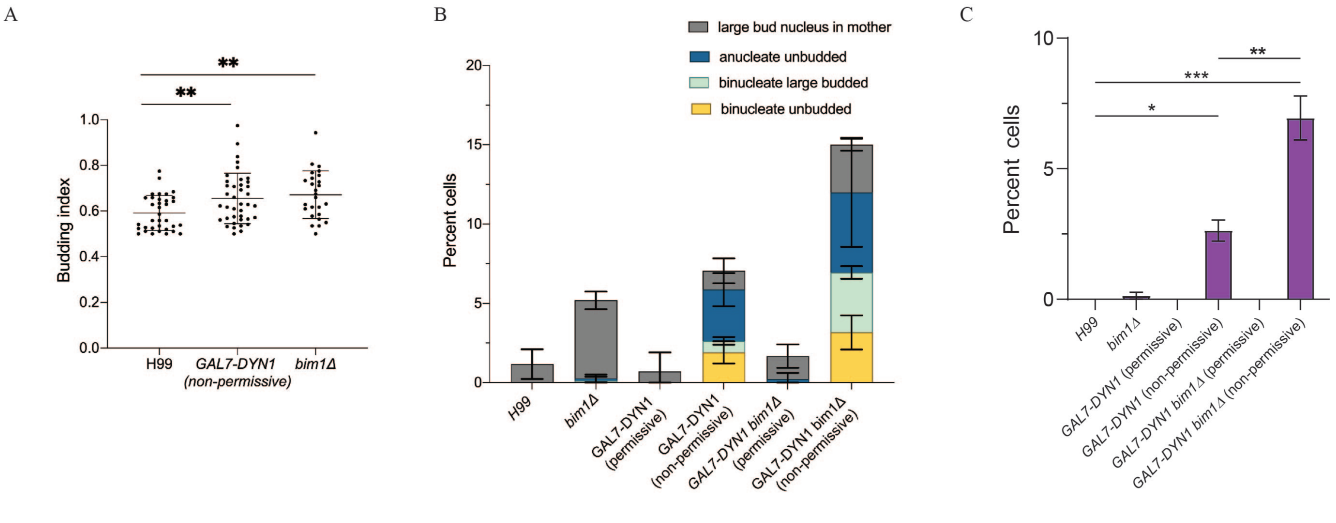


FIGURE S4 Nuclear segregation defects in dynein and Bim1 mutants. (A) Quantification of budding indices (B.I) in the unsegregated large budded cells from the wild-type, Bim1 deletion and Dynein depleted conditions. Analysis of large-budded cells with an unsegregated nucleus from the wild-type CNVY108, GAL7-DYN1 CNSD155 and bim1△ CNNV107 strains expressing histone GFP-H4. The cells with an unsegregated nucleus in the mother bud with the B.I. ≥ 0.5 are plotted (n ≥ 40). Mean and SD values are marked. **p-value<0.01, unpaired t-test. (B) The extent of abnormal nuclear division in the wild-type CNVY108, GAL7-DYN1 CNSD155, bim1△ CNNV107 and GAL7-DYN1 bim1△ CNSD156 cells expressing histone GFP-H4 was measured. Indicated abnormal nuclear phenotypes were quantified from different cell populations 12 h post-incubation in permissive (YPDextrose) and non-permissive (YPGalactose) conditions at 30°C. Wild-type and bim1△ strains were grown in YPDextrose. Mean and SEM values are marked. (C) Percentages of binucleated large-budded and unbudded cells were quantified using a nuclear segregation marker, histone GFP-H4, and plotted for wild-type CNVY108, GAL7-DYN1 CNSD155, bim1△ CNNV107 and GAL7-DYN1 bim1△ CNSD156 strains. Mean and SEM values are indicated. One-way ANOVA was performed to test the significance (n>100). ***p-value<0.001, **p-value<0.01 and *p-value<0.05.

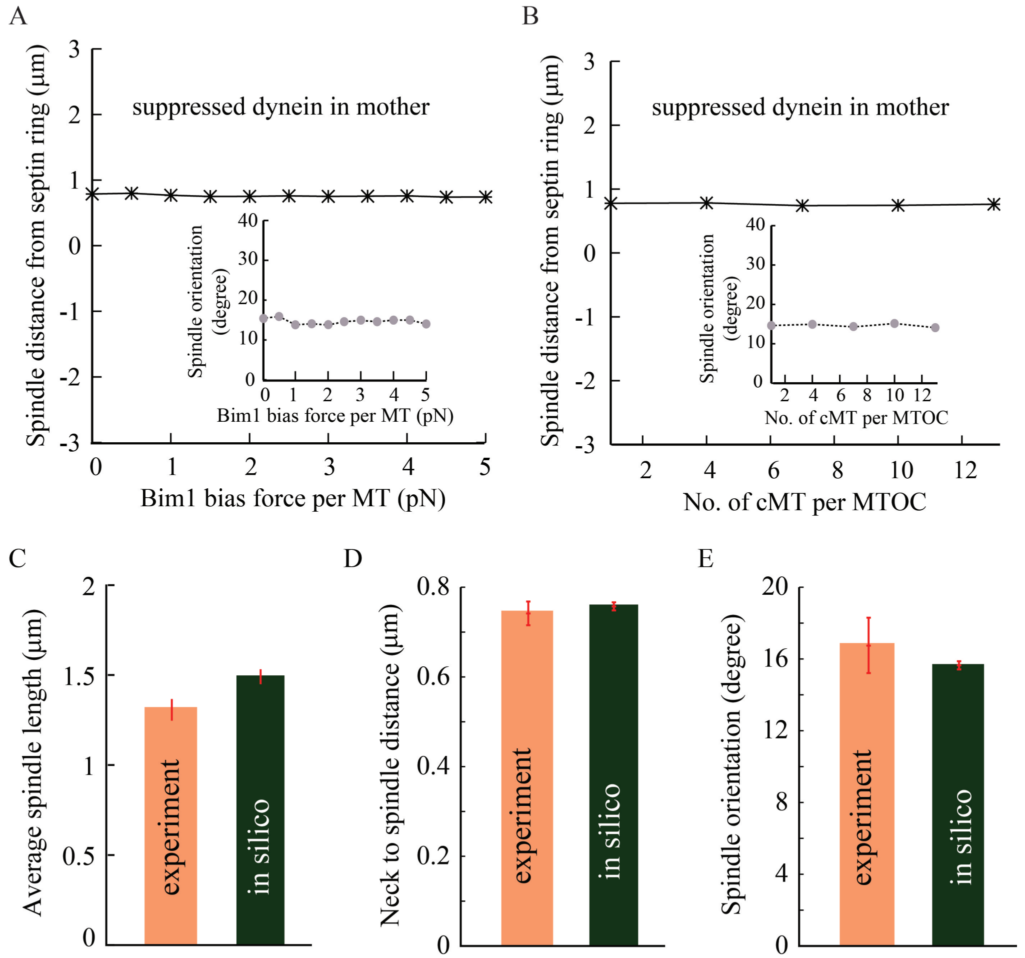


FIGURE S5 Multiple characteristics of spindle architecture and localization. (A-B) Spindle distance from septin ring and spindle orientation (inset) relative to the axis joining the center of the mother and daughter bud when the magnitude of force per MT due to Bim1 bias (A) and the number of cMT per MTOC (B) are varied. The spindle position and orientation turn out to be insensitive to the variations in Bim1 bias force per MT and the number of cMT per MTOC in the explored parameter regime. (C) The length of the spindle (n=68, experiment) in metaphase cells is shown. (D) The spindle to neck (junction of mother-daughter bud) distance in the wild-type (n=70, experiment). (E) The spindle orientation with respect to the mother-daughter axis (n=52, experiment). The +ve/-ve sign in the distance indicates the spindle position inside the daughter/mother bud. In all figures, n > 2000 for simulation and red bars indicate SEM (wherever shown).

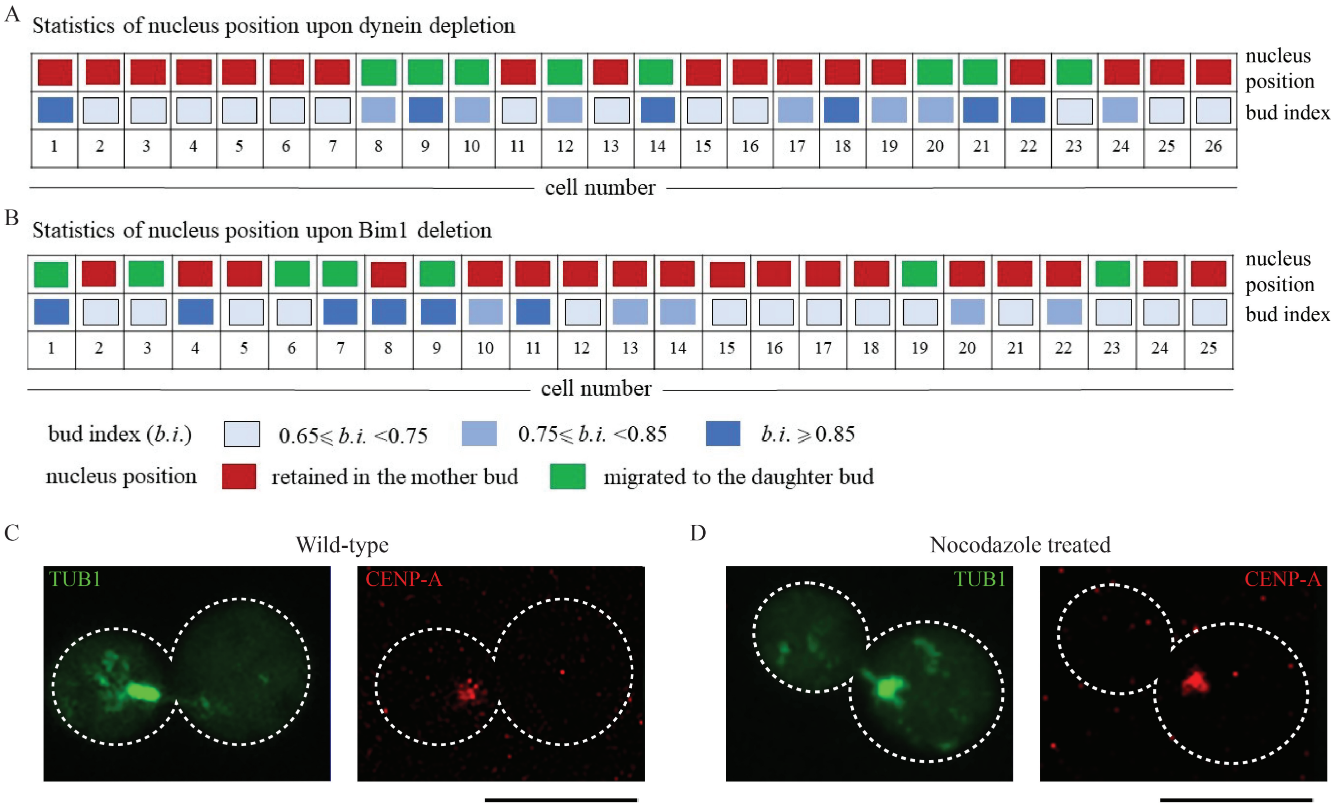


FIGURE S6 Nucleus position within cells upon various molecular perturbations. (A-B) Each column represents a cell indicating the color-coded status of its budding index and the localization of the nucleus within the cell. The statistics/distribution shows the nucleus position (whether the nucleus is localized in the mother bud or the daughter bud) upon dynein depletion (A), Bim1 deletion (B). (C-D) Images of cells showing localization of MTs (GFP-TUB1) and KTs (mCherry-CENP-A) in large budded wild-type (C) and nocodazole treated cells (D). Bar, 5 $\mu$m.

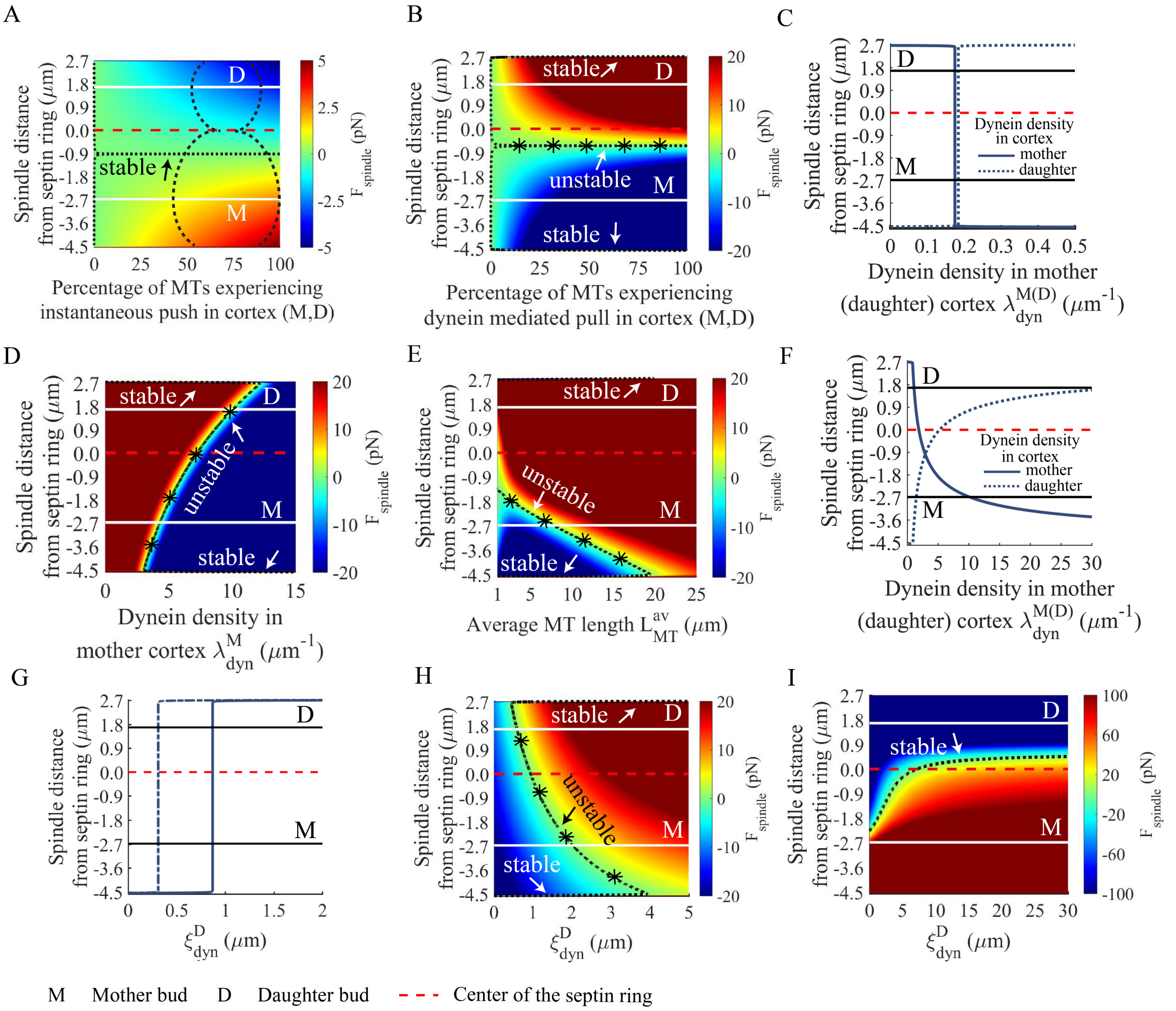


FIGURE S7 Effect of various force combinations on spindle positioning. In all figures, -ve/+ve distance refers to the spindle in mother (M)/daughter (D) bud. In the color maps, white solid lines denote the center of the daughter and mother bud; color bars represent net force on the spindle. (A) In the presence of instantaneous cortical push from the mother and daughter cortex, the spindle stabilizes (dotted black line) close to the septin ring inside the mother bud. (B) In the presence of dynein mediated cortical pull from the mother and daughter cortex, the spindle collapses onto the cell cortex. An unstable spindle position is found in between. (C) Variation in dynein pull from mother (daughter) cortex ($\lambda_{dyn}^{M(D)}$) with instantaneous cortical push from mother (daughter) cortex acting alongside. The spindle collapses either onto the mother cortex or onto the daughter cortex. Forces are exerted solely from the mother (daughter) cortex. (D) Spindle position upon variation in dynein density in the mother cortex ($\lambda_{dyn}^{M}$) with fixed dynein density in the daughter cortex. Instantaneous cortical push is present in both mother and daughter. (E) Spindle position upon variation in average MT length ($L_{MT}^{av}$) when instantaneous cortical push and dynein pull are combined. (F) Spindle position upon variation in dynein density in mother (daughter) cortex ($\lambda_{dyn}^{M(D)}$) when instantaneous push, dynein pull, and MT buckling in mother (daughter) cortex act together. MT buckling force (directed away from cortex) dominates close to the mother (daughter) cortex. Forces are exerted solely from the mother (daughter) cortex. (G) Variation of the characteristic length scale spanning the localized dynein patch in the daughter cortex ($\xi_{dyn}^{D}$) yields unstable spindle positions slotted in between stable fixed positions of the spindle position in the mother and the daughter cortex while mutually opposing MT mediated instantaneous push and dynein pull solely from the daughter cortex compete against each other. For this particular scenario, force transduction from the mother cortex is switched 'off'. Relevant parameters: $\lambda_{dyn}^{D}$ = 1 $\mu m^{-1}$ (solid line) and 2 $\mu m^{-1}$ (dashed line), $A_D$ = 5 pN. (H) Variation in the characteristic length scale accounting for the size of localized dynein patches in the daughter cortex ($\xi_{dyn}^{D}$) results in unstable fixed positions as the spindle undergoes a gradual spatial transition from the stable fixed positions at the mother cortex to the stable fixed positions at the daughter cortex. The force balance landscape is governed by cortical pull stemming from the mother and daughter cortex only (relevant parameters: $\lambda_{dyn}^{M}$ =1 $\mu m^{-1}$; $\lambda_{dyn}^{D}$ = 4.5 $\mu m^{-1}$). (I) In the presence of MT buckling transition together with instantaneous push and dynein pull from the mother and the daughter cortex, step-wise increment in $\xi_{dyn}^{D}$ enhances net pull toward the daughter resulting in a stable spindle positioning inside the daughter bud.

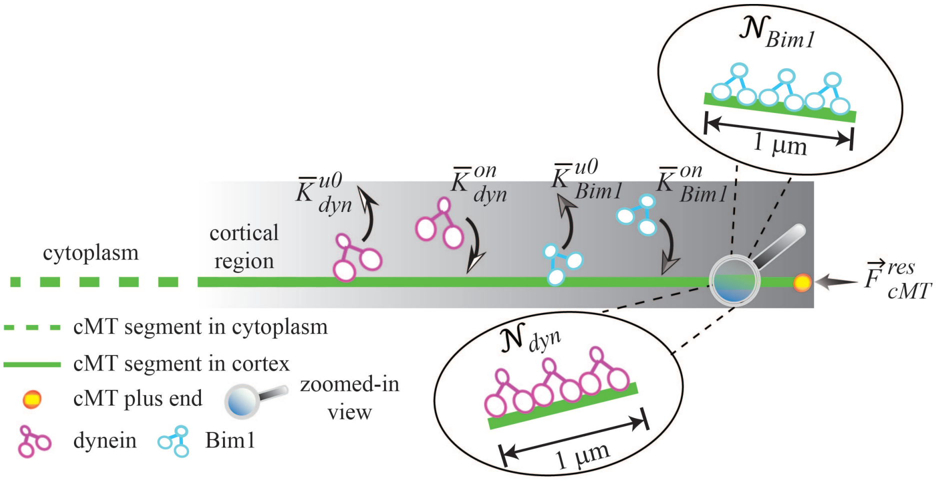


FIGURE S8 Model schematic showing the stochastic attachment-detachment of dynein and Bim1 inside the cell cortex. $\mathcal{N}_{dyn}$ ($\mathcal{N}_{Bim1}$) denotes the maximum number of dyneins (Bim1) that can attach to a cMT segment inside the cell cortex per $\mu$m. $\bar{K}^{on}_{dyn}$ ($\bar{K}^{on}_{Bim1}$) and $\bar{K}^{u0}_{dyn}$ ($\bar{K}^{u0}_{Bim1}$) represent the attachment rate of dynein (Bim1) on cMT segment inside the cell cortex and detachment rate of dynein (Bim1) on cMT segment inside the cell cortex respectively. $\vec{F}^{res}_{cMT}$ denotes the net 'pushing' force/load on a cMT acting against its growth.

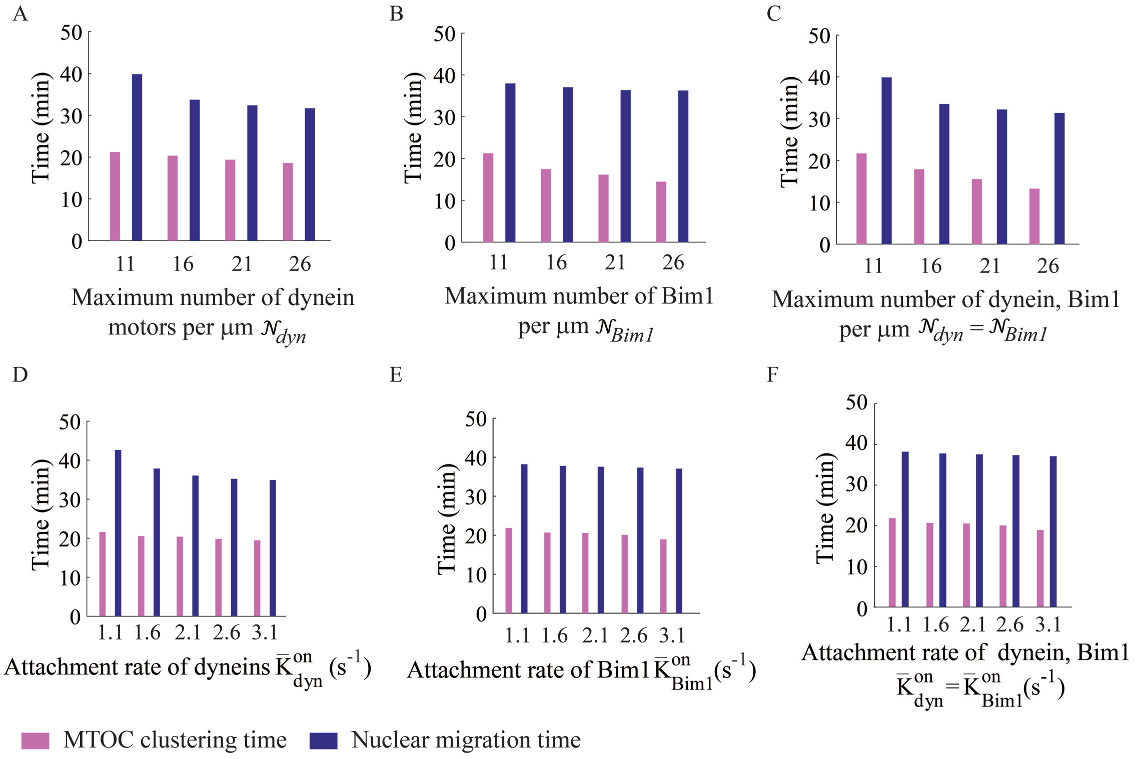


FIGURE S9 Sensitivity of MTOC clustering and nuclear migration time scales to the model parameters governing the stochastic attachment of motors on the MTs. (A-B) Dependence of the time scales of MTOC clustering and nuclear migration on the maximum number of dynein $\mathcal{N}_{dyn}$ (A) and Bim1 $\mathcal{N}_{Bim1}$ (B) that can attach to a cMT segment inside the cell cortex per $\mu$m. When $\mathcal{N}_{dyn}$ is varied, $\mathcal{N}_{Bim1}$ is kept fixed at 12 per $\mu$m and vice versa. Other parameters: $\bar{K}^{on}_{dyn} = \bar{K}^{on}_{Bim1} = 1.6\ s^{-1}$; $\bar{K}^{u0}_{dyn} = \bar{K}^{u0}_{Bim1} = 0.27\ s^{-1}$; $F^{detach}_{dyn} = F^{detach}_{Bim1} = 0.67$ pN. (C) Simultaneous variation in the maximum number of dynein and Bim1 per $\mu$m ($\mathcal{N}_{dyn}$ and $\mathcal{N}_{Bim1}$) does not significantly alter the timescales of MTOC clustering and nuclear migration. For simplicity, we took $\mathcal{N}_{dyn} = \mathcal{N}_{Bim1}$ in this variation. Other parameters: $\bar{K}^{on}_{dyn} = \bar{K}^{on}_{Bim1} = 1.6\ s^{-1}$; $\bar{K}^{u0}_{dyn} = \bar{K}^{u0}_{Bim1} = 0.27\ s^{-1}$; $F^{detach}_{dyn} = F^{detach}_{Bim1} = 0.67$ pN. (D-E) Dependence of MTOC clustering and nuclear migration time scales on the attachment rates of dynein $\bar{K}^{on}_{dyn}$ (D) and Bim1 $\bar{K}^{on}_{Bim1}$ (E). When the attachment rate of dynein is varied, the attachment rate of Bim1 is kept fixed at 1.6 $s^{-1}$ and vice versa. Other parameters: $\mathcal{N}_{dyn} = \mathcal{N}_{Bim1} = 12$ per $\mu$m; $\bar{K}^{u0}_{dyn} = \bar{K}^{u0}_{Bim1} = 0.27\ s^{-1}$; $F^{detach}_{dyn} = F^{detach}_{Bim1} = 0.67$ pN. (F) Simultaneous variation in the attachment rates of dynein and Bim1 does not significantly alter the timescales of MTOC clustering and nuclear migration. For the sake of simplicity, we considered $\bar{K}^{on}_{dyn} = \bar{K}^{on}_{Bim1}$ in this variation. Other parameters: $\mathcal{N}_{dyn} = \mathcal{N}_{Bim1} = 12$ per $\mu$m; $\bar{K}^{u0}_{dyn} = \bar{K}^{u0}_{Bim1} = 0.27\ s^{-1}$; $F^{detach}_{dyn} = F^{detach}_{Bim1} = 0.67$ pN.

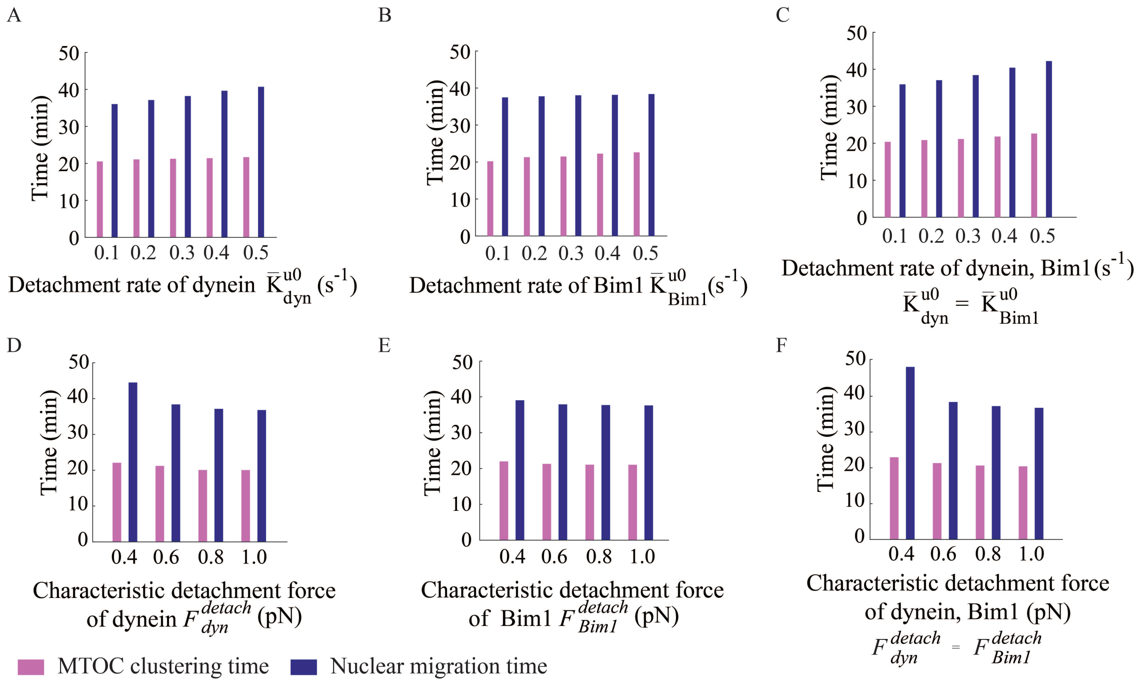


FIGURE S10 Sensitivity of MTOC clustering and nuclear migration time scales to the model parameters governing the stochastic detachment of motors. (A-B) Dependence of the time scales of MTOC clustering and nuclear migration on the detachment rates of dynein $\bar{K}^{u0}_{dyn}$ (A) and Bim1 $\bar{K}^{u0}_{Bim1}$ (B). When $\bar{K}^{u0}_{dyn}$ is varied, $\bar{K}^{u0}_{Bim1}$ is kept fixed at 0.27 $s^{-1}$ and vice versa. Other parameters: $\mathcal{N}_{dyn} = \mathcal{N}_{Bim1}$ = 12 per $\mu$m; $\bar{K}^{on}_{dyn} = \bar{K}^{on}_{Bim1}$ = 1.6 $s^{-1}$; $F^{detach}_{dyn} = F^{detach}_{Bim1}$ = 0.67 pN. (C) Simultaneous variation in the detachment rates of dynein and Bim1 ($\bar{K}^{u0}_{dyn}$ and $\bar{K}^{u0}_{Bim1}$) does not significantly alter the timescales of MTOC clustering and nuclear migration. For the sake of simplicity, we considered $\bar{K}^{u0}_{dyn} = \bar{K}^{u0}_{Bim1}$ in this variation. Other parameters: $\mathcal{N}_{dyn} = \mathcal{N}_{Bim1}$ = 12 per $\mu$m; $\bar{K}^{on}_{dyn} = \bar{K}^{on}_{Bim1}$ = 1.6 $s^{-1}$; $F^{detach}_{dyn} = F^{detach}_{Bim1}$ = 0.67 pN. (D-E) Dependence of MTOC clustering and nuclear migration time scales on the characteristic detachment force of dynein $F^{detach}_{dyn}$ (D) and Bim1 $F^{detach}_{Bim1}$ (E). Other parameters:$\mathcal{N}_{dyn} = \mathcal{N}_{Bim1}$ = 12 per $\mu$m, $\bar{K}^{on}_{dyn} = \bar{K}^{on}_{Bim1}$ = 1.6 $s^{-1}$; $\bar{K}^{u0}_{dyn} = \bar{K}^{u0}_{Bim1}$ = 0.27 $s^{-1}$. When the characteristic detachment force of dynein $F^{detach}_{dyn}$ is varied, the characteristic detachment force of Bim1 $F^{detach}_{Bim1}$ is kept fixed at 0.67 pN and vice versa. (F) Simultaneous variation in the characteristic detachment forces of dynein and Bim1 ($F^{detach}_{dyn}$ and $F^{detach}_{Bim1}$) does not significantly alter the timescales of MTOC clustering and nuclear migration. For simplicity, we chose $F^{detach}_{dyn} = F^{detach}_{Bim1}$ in this variation. Other parameters:$\mathcal{N}_{dyn} = \mathcal{N}_{Bim1}$ = 12 per $\mu$m, $\bar{K}^{on}_{dyn} = \bar{K}^{on}_{Bim1}$ = 1.6 $s^{-1}$, $\bar{K}^{u0}_{dyn} = \bar{K}^{u0}_{Bim1}$ = 0.27 $s^{-1}$.

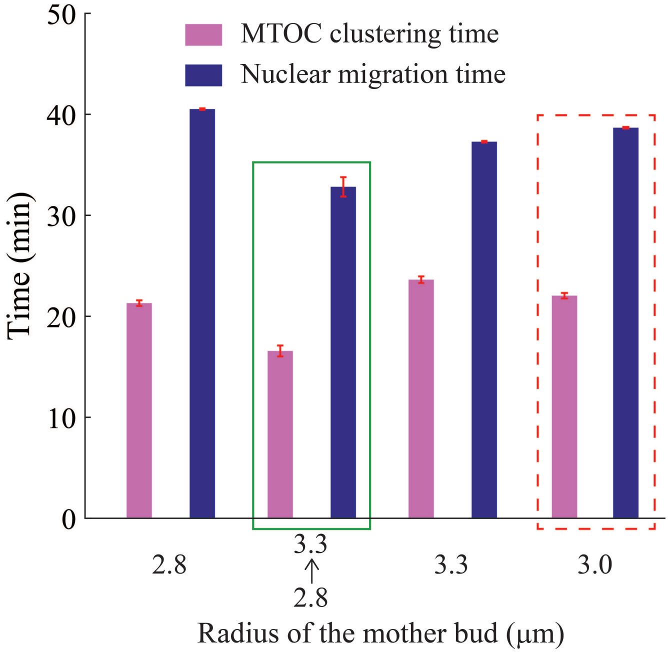


FIGURE S11 Dependence of the time scales of MTOC clustering and nuclear migration on the mother bud size. The set of bars enclosed by solid green rectangle represents the case in which the mother bud radius slowly grows from 2.8 $\mu$m to 3.3 $\mu$m, in concurrence with the much faster growth of daughter bud during the simulation. The set of bars enclosed by red dashed rectangle refers to the specific simulations in which the parameter values listed in Table S2 are used. In the currently explored parameter range, the mother bud size variations do not change the time scales of MTOC clustering and nuclear migration significantly. Note that in all simulations (unless mentioned otherwise), the mother bud radius is kept fixed at 3.0 $\mu$m (Table S2).

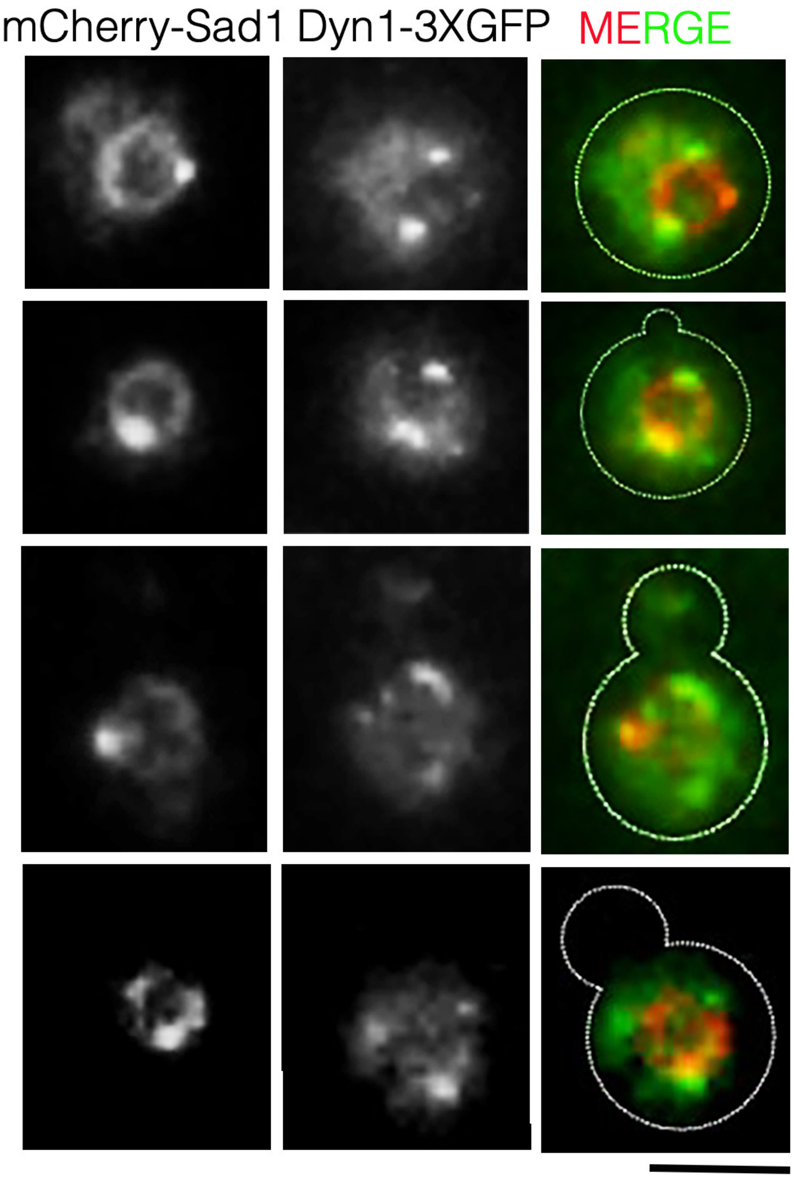


FIGURE S12 Snapshots depicting the localization of the nuclear envelope protein, Sad1 (mCherry-Sad1, depicted in red in the MERGE panel) and Dynein (Dyn1-3xGFP, depicted in green in the MERGE panel) at different stages of the cell cycle in C. neoformans. mCherry-Sad1 was expressed using the GAL7 promoter and localized along the nuclear periphery. Bar, 5 $\mu$m.

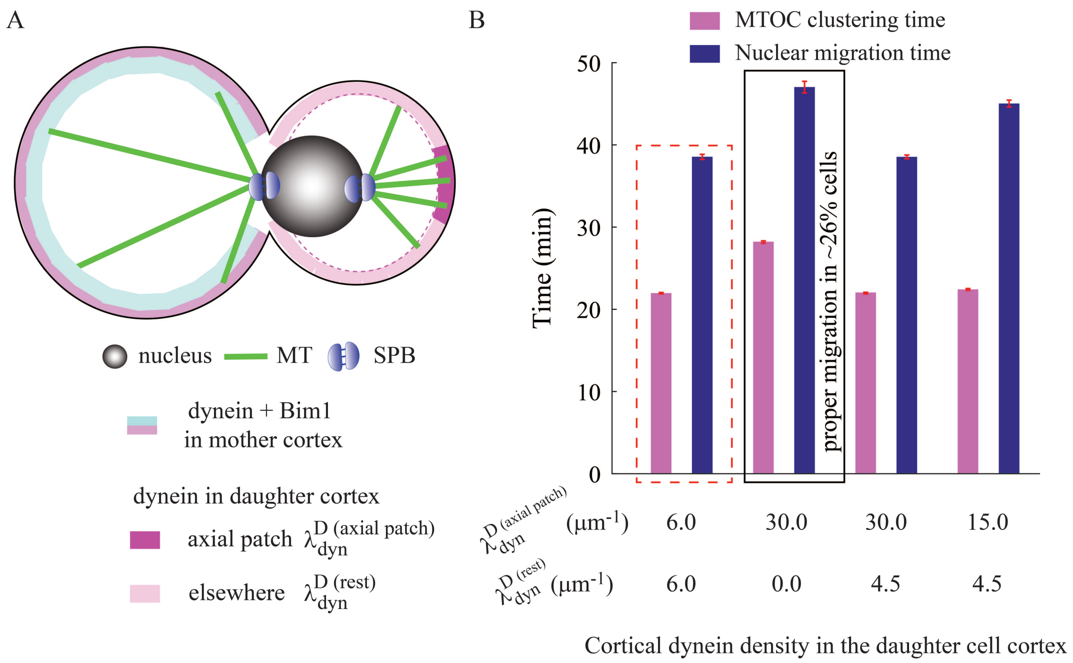


FIGURE S13 Additional numerical experiments with differential dynein density profile in mother and daughter bud cell cortex (agent based model). (A) Schematic diagram illustrates the the cortical dynein density profile considered in the agent based simulations of Fig. S13B. (B) Dependence of MTOC clustering and nuclear migration time scales on various spatial profiles of cortical dyneins regulated by model parameters $\lambda_{dyn}^{D\ (axial\ patch)}$ and $\lambda_{dyn}^{D\ (rest)}$. Quite evidently, the parameter choices corresponding to the set of bars enclosed by red dashed rectangle represent a configuration with uniform dynein density (Table S2).

### SUPPORTING MOVIES

**M1.** Cell cycle stages in *C. neoformans* wild-type cell. Wild-type cells expressing GFP-Tub1 were imaged after every 2 min by confocal microscopy. KTs (CENP-A) were visualized by mCherry-tagged CENP-A. The video corresponds to the still images from Fig. 1A.

**M2.** MTOC clustering facilitated via inter-cMT coupling at the nuclear envelope (NE). MTOCs are blue; KTs are magenta; MTs are green.

**M3.** MTOC clustering facilitated via MT-cell cortex interaction. Force produced by single dynein on each MT at mother cortex ($|\vec{f}_{dyn}|$) is chosen to be 1.0 pN. Bias force towards septin ring produced by single Bim1 on each MT at mother cortex ($|\vec{f}_{Bim1}|$) is chosen to be 0.0 pN. MTOCs are blue; KTs are magenta; MTs are green.

**M4.** MTOC clustering facilitated via MT-cell cortex interaction (enhanced Bim1 bias + suppressed dynein activity). Force produced by single dynein on each MT at mother cortex ($|\vec{f}_{dyn}|$) is chosen to be 0.0 pN. Bias force towards septin ring produced by single Bim1 on each MT at mother cortex ($|\vec{f}_{Bim1}|$) is chosen to be 1.0 pN. MTOCs are blue; KTs are magenta; MTs are green.

| Strains used in this study | | |
|---|---|---|
| Strain name | Genotype | Reference |
| CNVY108 | MAT$\alpha$ H99::GFP-H4-NAT (pVY3) | (11) |
| CNVY107 | MATa H99::GFP-$\alpha$TUB1-NAT (pLKB35), H99::mCherry-CSE4-NEO (pLKB74) | (1) |
| CNNV105 | MATa H99::GFP-$\alpha$TUB1-NAT (pLKB35), H99::mCherry-CENP-A-NEO (pLKB74), IPL1:: GAL7p-IPL1-HygB | (1) |
| CNNV107 | MAT$\alpha$ H99::GFP-H4-NAT (pLKB35) + BIM1△::NEO | (1) |
| CNNV110 | MAT$\alpha$ H99::GFP-H4-NAT (pVY3), DYN1:: GAL7p-DYN1-HygB | (1) |
| CNNV120 | MAT$\alpha$ SPC98::SPC98p-SPC98-3xGFP-NEO | This study |
| CNSD152 | MAT$\alpha$ H99 SAD1p::GAL7p-mCherry-SAD1-HYG DYN1:: DYN1p-DYN1-3xGFP-NEO | This study |
| CNSD155 | MAT$\alpha$ H99::GFP-H4-NAT (pVY3) DYN1::GAL7p-DYN1-HygB | This study |
| CNSD156 | MAT$\alpha$ H99::GFP-H4-NAT (pVY3) BIM1△:: NEO DYN1::GAL7p-DYN1-HygB | This study |
| CNVY182 | MAT$\alpha$ H99 SAD1p::GAL7p-mCherry-SAD1-HYG, GFP-NDC1-NAT (pVY4) | (10) |
| CNNV119 | MAT$\alpha$ BIM1::BIM1p-BIM1-3xGFP-NEO | (1) |
| CNNV116 | MAT$\alpha$ DYN1:: DYN1p-DYN1-3xGFP-NEO, IPL1::GAL7p-IPL1-HygB | (1) |
| CNNV121 | MAT$\alpha$ DYN1:: DYN1p-DYN1-3xGFP-NEO | This study |

| Primers used in this study | | |
|---|---|---|
| Primer name | Sequence | Description |
| VYP202 | GCCAACATGGACATCCGTCTCC | Forward primer for GAL7-mCherry SAD1 cassette |
| VYP203 | CGACTGCCGGAGTCCATGCTC | Reverse primer for GAL7-mCherry SAD1 cassette |
| VYP131 | ATACGCGAGTCCAACTTCGC | Confirmatory forward primer for GAL7 DYN1 cassette |
| NV356 | ACGCGTCGACAGTCCTTCATCAAGATGTTATGC | Reverse cloning primer for upstream of GAL7p-DYN1 cassette |
| NV404 | ATTTGCGGCCGCTGAGATCATGGTCGCTTACATC | Forward cloning primer for downstream of GAL7p-DYN1 cassette |
| NV405 | CTAGACACTCTTATCAGCACCATCC | Confirmatory reverse primer for GAL7 DYN1 cassette |

Table S1: Strains and primers used in this study

| Abbreviation | Meaning | Value | Reference |
|---|---|---|---|
| $R_M$ | Radius of the mother bud | 3 $\mu$m | (1) |
| $r_D$ | Radius of the daughter bud | 0.0-2.15 $\mu$m | (1) |
| $r_{nuc}$ | Radius of the nucleus | 1.0 $\mu$m | (1) |
| $r_{SPB}$ | Radius of single SPB | 0.125 $\mu$m | (21, 22) |
| $k_{cor}$ | Spring constant of the cortex | 5.0 pN/$\mu$m | (2, 23) |
| $\eta_{cyt}$ | Viscosity of cytoplasm | 5.0 pN s/$\mu$m$^2$ | (2, 23) |
| $\eta_{nu}$ | Viscosity of nucleoplasm | 10.0 pN s/$\mu$m$^2$ | (2, 23) |
| $\eta_{NE}$ | Effective viscosity of NE | 10-15 pN s/$\mu$m$^2$ | (2, 23) |
| $v_g$ | MT growth velocity | 10.4 $\mu$m min$^{-1}$ | (1, 24, 25) |
| $v_s$ | MT shrinkage velocity | 28.6 $\mu$m min$^{-1}$ | (1, 24, 25) |
| $f_c$ | Catastrophe frequency of MT | 1.0 min$^{-1}$ | (1, 24, 25) |
| $f_r$ | Rescue frequency of MT | 0.02 min$^{-1}$ | (1, 24, 25) |
| $f_c^{stall}$ | Catastrophe rate of stalled MT | 0.04 s$^{-1}$ | (26) |
| $f_{stall}$ | MT stall force | 1.7 pN | (27) |
| $f_{dyn}$ | Force produced by single dynein | 1.0 pN | (12, 28) |
| $f_{Bim1}$ | Force produced by single Bim1 | 1.0 pN | (1) |
| $\lambda_{dyn}^{M(D)}$ | Number of dyneins per unit length per MT | 6.0 /$\mu$m | (29) |
| $\lambda_{ipMT}$ | Number of ipMT motors per unit length per MT | 1.0 /$\mu$m | (23) |
| $\lambda_{dyn}^{ovl}$ | Number of minus ended motors per unit length at NE | 1.0 /$\mu$m | This study |
| $f_{kinesin-5}$ | Force produced by single kinesin-5 motor | 1.0 pN | (2) |
| $K_{cohesin}$ | Spring constant of the cohesion springs | 0.1 pN/$\mu$m | (30) |
| $K_c$ | Spring constant of KT-kMT connection springs | 10.0 pN/$\mu$m | (23, 31) |
| $K_{fibril}$ | Spring constant of the KT fibril | 5.0 pN/$\mu$m | (23, 31) |
| $D_{buckle}$ | Buckling amplitude per MT | 200.0 pN $\mu$m$^2$ | (4) |
| | Additional parameters for analytical model | | |
| $A_M$ ($A_D$) | Instantaneous cortical push amplitude at mother(daughter) | 5 pN | This study |
| $B_M$ ($B_D$) | Cortical pull amplitude at mother(daughter) | 5 pN | This study |
| $D_M$ ($D_D$) | Buckling amplitude at mother(daughter) | 1000 pN $\mu$m$^2$ | (4), this study |
| $C_M$ ($C_D$) | Steric repulsion amplitude at mother(daughter) | 10 pN | (4), this study |
| $\zeta_M$ ($\zeta_D$) | Range of steric repulsion at mother(daughter) | 1000 | (4), this study |
| $L_{MT}^{av}$ | Average cMT length | 10 $\mu$m | This study |
| $a$ | Spindle half length | 1.0 $\mu$m | This study |
| $r_D$ | Radius of the daughter bud | 2.15 $\mu$m | This study |
| $l_c$ | Cortex width | 0.2 $\mu$m | (2, 32) |

Table S2: List of parameters chosen for the model analysis

| Interaction mechanism | Clustering time (min) | % of MTOCs clustered within 25 min |
|---|---|---|
| cMT-cell cortex interaction | ~ 240 | ~ 40 |
| Inter cMT coupling at the NE | ~ 23 | ~ 100 |
| cMT-cell cortex interaction with diminished cortical pull and enhanced Bim1 bias | ~ 25 (1) | ~ 99 |

Table S3: Various mechanisms of MTOC clustering as examined by the *in silico* model

| Instantaneous push | Cortical pull | MT buckle | Variation | Magnitude | Spindle position | Plot |
|---|---|---|---|---|---|---|
| | | | | low | M* (stable) | |
| ✓(M,D) | ✓(M,D) | ✗(M,D) | $\lambda^{D}_{dyn}$ | intermediate | M* (stable), D* (stable) & unstable points in between | Fig. 5A |
| | | | | high | D* (stable) | |
| | | | | low | D* (stable) | |
| ✓(M), ✗(D) | ✓(M), ✗(D) | ✗(M,D) | $\lambda^{M}_{dyn}$ | intermediate | 'sharp' transition† | Fig. S7C |
| | | | | high | M* (stable) | |
| | | | | low | M* (stable) | |
| ✗(M), ✓(D) | ✗(M), ✓(D) | ✗(M,D) | $\lambda^{D}_{dyn}$ | intermediate | 'sharp' transition† | Fig. S7C |
| | | | | high | D* (stable) | |
| | | | | low | D* (stable), M* (stable) & unstable points in between | |
| ✓(M,D) | ✓(M,D) | ✗(M,D) | $L^{av}_{MT}$ | intermediate | " | Fig. S7E |
| | | | | high | D* (stable) | |
| | | | | low | D* (stable) | |
| ✓(M), ✗(D) | ✓(M), ✗(D) | ✓(M), ✗(D) | $\lambda^{M}_{dyn}$ | intermediate | stable transition from D to M | Fig. S7F |
| | | | | high | M (stable, slightly away from the cell cortex) | |
| | | | | low | M* (stable) | |
| ✗(M), ✓(D) | ✗(M), ✓(D) | ✗(M), ✓(D) | $\lambda^{D}_{dyn}$ | intermediate | stable transition from M to D | Fig. S7F |
| | | | | high | D (stable, slightly away from the cell cortex) | |
| | | | | low | M (stable) | |
| ✓(M), ✓(D) | ✓(M), ✓(D) | ✓(M), ✓(D) | $L^{MT}_{av}$ | intermediate | D(stable), **close to septin ring** | Fig. 5C |
| | | | | high | ” | |

Table S4: Analysis of parameter sensitivity in spindle positioning (analytical model). M, D stand for mother and daughter bud respectively. * Spindle collapsed onto the cell cortex. The term 'collapse' means that the spindle is pulled to a very close proximity of the cell cortex. † Below/above this particular value of dynein density, spindle jumps sharply to the stable point at mother (daughter) bud cell cortex from stable point at daughter (mother) bud cell cortex with no unstable point in between the crossover.

| forces | origin | direction |
|---|---|---|
| instantaneous push ($F^{cell-mem}_{push-inst}$) | MT tip hitting the cell wall | directed along the MT, away from the cell wall |
| dynein mediated cortical pull ($F^{cor}_{dyn}$) | MT sliding in cell cortex | directed along the MT, toward the cell cortex |
| Bim1 bias ($F^{cor}_{Bim1}$) | cortical sliding of MTs toward septin ring | directed toward the septin ring |
| MT buckling ($F^{cor}_{buckle}$) | MTs impinged on the cell boundary | directed along the MT, away from the cell cortex |
| inter cMT coupling | overlapping antiparallel MTs grazing the NE | directed along the MT, away from the cell cortex |
| push due to MT polymerization in cortex ($f^{MT-poly}_{cor}$) | cortical resistance on polymerizing MT tip | directed along the MT, away from the cell cortex |
| force at ipMT-ipMT overlap ($f_{ipMT}$) | collective activity of kinesin-5 motors at ipMT-ipMT overlap | directed along the MT, leading to the separation of the SPBs |
| push by growing kMTs ($f^{growth}_{push}$) | kMT tip penetrating into the KT | directed along the MT, away from the KT |
| pull by shrinking kMTs ($f^{shrinkage}_{pull}$) | separation between kMT tip and KT | directed along the MT, pulling the KT |

Table S5: Description of MT based forces present in the *in silico* model

### SUPPORTING REFERENCES


1. Varshney, N., S. Som, S. Chatterjee, S. Sridhar, D. Bhattacharyya, R. Paul, and K. Sanyal, 2019. Spatio-temporal regulation of nuclear division by Aurora B kinase Ipl1 in Cryptococcus neoformans. *PLoS Genet* 15(2):e1007959.

2. Sutradhar, S., V. Yadav, S. Sridhar, L. Sreekumar, D. Bhattacharyya, S. K. Ghosh, R. Paul, and K. Sanyal, 2015. A comprehensive model to predict mitotic division in budding yeasts. *Molecular Biology of the Cell* 26:3954–3965.

3. Gittes, F., E. Meyhöfer, S. Baek, and J. Howard, 1996. Directional loading of the kinesin motor molecule as it buckles a microtubule. *Biophys J* 70:418–429.

4. Som, S., S. Chatterjee, and R. Paul, 2019. Mechanistic three-dimensional model to study centrosome positioning in the interphase cell. *Phys. Rev. E* 99:012409.

5. Ferenz, N. P., R. Paul, C. Fagerstrom, A. Mogilner, and P. Wadsworth, 2009. Dynein antagonizes Eg5 by crosslinking and sliding antiparallel microtubules. *Curr Biol.* 19(21):1833–1838.

6. Chatterjee, S., A. Sarkar, J. Zhu, A. Khodjakov, A. Mogilner, and R. Paul, 2020. Mechanics of multi-centrosomal clustering in bipolar mitotic spindles. *Biophysical Journal* 119(2):434–447.

7. Bertalan, Z., Z. Budrikis, C. A. M. L. Porta, and S. Zapperi, 2015. Role of the Number of Microtubules inChromosome Segregation during Cell Division. *PLOS ONE* 10.1371/journal.pone.0141305.

8. Wollman, R., E. Cytrynbaum, J. Jones, T. Meyer, J. Scholey, and A. Mogilner, 2005. Efficient Chromosome Capture Requires a Bias in the 'Search-and-Capture' Process during Mitotic-Spindle Assembly. *Current Biology* 15:828–832.

9. Sarkar, A., H. Rieger, and R. Paul, 2019. Search and Capture Efficiency of Dynamic Microtubules for Centrosome Relocation duringIS Formation. *Biophysical Journal* 116:2079–2091.

10. Yadav, V., and K. Sanyal, 2018. Sad1 Spatiotemporally Regulates Kinetochore Clustering To Ensure High-Fidelity Chromosome Segregation in the Human Fungal Pathogen Cryptococcus neoformans. *mSphere* 3(4):e00190–18.

11. Kozubowski, L., V. Yadav, G. Chatterjee, S. Sridhar, M. Yamaguchi, S. Kawamoto, I. Bose, J. Heitman, and K. Sanyal, 2013. Ordered Kinetochore Assembly in the Human-Pathogenic Basidiomycetous Yeast Cryptococcus neoformans. *mBio* 4(5):00614–13.

12. Muller, M. J. I., S. Klumpp, and R. Lipowsky, 2008. Tug-of-war as a cooperative mechanism for bidirectional cargo transport by molecular motors. *Proc Natl Acad Sci USA* 105(12):4609–14.

13. Ghanti, D., R. W. Friddle, and D. Chowdhury, 2018. Strength and stability of active ligand-receptor bonds: A microtubule attached to a wall by molecular motor tethers. *PHYSICAL REVIEW E* 98:042415.

14. H.A.Kramers, 1940. Brownian motion in a field of force and the diffusion model of chemical reactions. *Physica* 7(4)):284–304.

15. Bell, G., 1978. Models for the specific adhesion of cells to cells. *Science* 200(4342):618–627.

16. Kramer, A., B. Maier, and J. Bartek, 2011. Centrosome clustering and chromosomal (in)stability: A matter of life and death. *Molecular Oncology* 5(4):324–335.

17. Singh, M. P., R. Mallik, S. P. Gross, and C. C. Yu, 2005. Monte Carlo modeling of single-molecule cytoplasmic dynein. *PNAS* 102(34):12059–12064.

18. King, S. J., and T. A. Schroer, 2000. Dynactin increases the processivity of the cytoplasmic dynein motor. *Nature Cell Biology* 2:20–24.

19. Reck-Peterson, S. L., A. Yildiz, A. P.Carter, A. Gennerich, N. Zhang, and R. D.Vale, 2006. Single-Molecule Analysis of Dynein Processivity and Stepping Behavior. *Cell* 126:335–348.

20. Mallik, R., B. C. Carter, S. A. Lex, S. J. King, and S. P. Gross, 2004. Cytoplasmic dynein functions as a gear in response to load. *Nature* 427:649–652.

21. Lee, I.-J., N. Wang, W. Hu, K. Schott, J. Bahler, T. H. G. Jr., J. R. Pringle, L.-L. Du, and J.-Q. Wu, 2014. Regulation of spindle pole body assembly and cytokinesis by the centrin-binding protein Sfi1 in fission yeast. *Mol Biol Cell* 25(18):2735–2749.

22. Seybold, C., and E. Schiebel, 2013. Spindle pole bodies. *Current Biology* 23(19):R858–R860.

23. G.Civelekoglu-Scholey, D.J.Sharp, A. Mogilner, and J. Scholey, 2006. Model of Chromosome Motility in Drosophila Embryos: Adaptation of a General Mechanism for Rapid Mitosis. *Biophysical Journal* 19(11):3966–3982.

24. Fink, G., I. Schuchardt, J. Colombelli, E. Stelzer, and G. Steinberg, 2006. Dynein-mediated pulling forces drive rapid mitotic spindle elongation in Ustilago maydis. *EMBO J.* 25(20):4897–908.

25. Finley, K. R., K. J. Bouchonville, A. Quick, and J. Berman, 2008. Dynein-dependent nuclear dynamics affect morphogenesis in Candida albicans by means of the Bub2p spindle checkpoint. *J Cell Sci* 121:466–76.

26. Letort, G., F. Nedelec, L. Blanchoin, and M. Thery, 2016. Centrosome centering and decentering by microtubule network rearrangement. *Mol Biol Cell* 27(18):2833–43.

27. Dogterom, M., and B. Yurke, 1997. Measurement of the force-velocity relation for growing microtubules. *Science* 278(5339):856–60.

28. Soppina, V., A. K. Rai, A. J. Ramaiya, P. Barak, and R. Mallik, 2009. Tug-of-war between dissimilar teams of microtubule motors regulates transport and fission of endosomes. *Proc Natl Acad Sci USA* 106(46):19381–6.

29. Markus, S. M., K. M. Plevock, B. J. S. Germain, J. J. Punch, C. W. Meaden, and W. Lee, 2011. Quantitative Analysis of Pac1/LIS1-mediated Dynein Targeting: Implications for Regulation of Dynein Activity in Budding Yeast. *Cytoskeleton (Hoboken, NJ)* 68(3):157–74.

30. Joglekar, A. P., and A. J. Hunt, 2002. A simple, mechanistic model for directional instability during mitotic chromosome movements. *Biophys J* 83(1):42–58.

31. Sau, S., S. Sutradhar, R. Paul, and P. Sinha, 2014. Budding yeast kinetochore proteins, Chl4 and Ctf19, are required to maintain SPB-centromere proximity during G1 and late anaphase. *PLoS One* 9(7):e101294.

32. Rodal, A., L. K. L, B. Goode, D. D. DG, and J. Hartwig, 2005. Actin and septin ultrastructures at the budding yeast cell cortex. *Mol Biol Cell* 16:372–384.
